# Deep Generative Models of Protein Structure Uncover Distant Relationships Across a Continuous Fold Space

**DOI:** 10.1101/2022.07.29.501943

**Authors:** Eli J. Draizen, Stella Veretnik, Cameron Mura, Philip E. Bourne

## Abstract

Our views of fold space implicitly rest upon many assumptions that impact how we analyze, interpret and understand protein structure, function and evolution. For instance, is there an optimal granularity in viewing protein structural similarities (e.g., architecture, topology or some other level)? Similarly, the discrete/ continuous dichotomy of fold space is central, but remains unresolved. Discrete views of fold space bin ‘similar’ folds into distinct, non-overlapping groups; unfortunately, such binning can miss remote relationships. While hierarchical systems like CATH are indispensable resources, less heuristic and more conceptually flexible approaches could enable more nuanced explorations of fold space. Building upon an ‘Urfold’ model of protein structure, here we present a deep generative modeling framework, termed ‘DeepUrfold’, for analyzing protein relationships at scale. DeepUrfold’s learned embeddings occupy high-dimensional latent spaces that can be distilled for a given protein in terms of an amalgamated representation uniting sequence, structure and biophysical properties. This approach is structure-*guided*, versus being purely structure-based, and DeepUrfold learns representations that, in a sense, ‘define’ superfamilies. Deploying DeepUrfold with CATH reveals evolutionarily-remote relationships that evade existing methodologies, and suggests a new, mostly-continuous view of fold space—a view that extends beyond simple geometric similarity, towards the realm of *integrated* sequence***↔***structure***↔***function properties.

## Introduction

The precise historical trajectory of the protein universe [1] remains quite murky, and ikely corresponds to an evolution from (proto-) peptides, to protein domains, to multidomain proteins [2]. Presumably, the protein universe—by which we mean the set of all unique protein sequences (known or unknown, natural or engineered, ancestral or extant)—did not spontaneously arise with intact, full-sized domains. Rather, smaller, sub-domain–sized protein fragments likely preceded more modern domains; the genomic elements encoding these primitive fragments were subject to natural evoutionary processes of duplication, mutation and recombination to give rise to extant domains found in contemporary proteins [2–6]. Our ability to detect common polypeptide fragments, shared amongst at least two domains (in terms of either sequence or structure), relies upon having (i) a similarity metric that is sensitive and accurate, and (ii) a suitable random/background distribution (i.e., null model) for distances under this metric; historically, such metrics have been rooted in the comparison of either amino acid sequences or three-dimensional (3D) structures, often for purposes of exploring protein fold space. The recent advent of high-accuracy structure prediction 7, 8], enabled by deep learning, presents new opportunities to explore fold space; to do so effectively requires new methods to accurately and sensitively detect weak/distant relationships.

### Fold Space, Structural Transitions & Protein Fragments

Fold space, as the collection of all unique protein folds (Supplementary Note 1), corresponds to a many-to-one mapping: vast swaths of sequence space map to fold ***𝒜***, another vast swath maps to fold *ℬ*, a narrower range might map to fold ***𝒞***, and so on. Two proteins that are closely related (evolutionarily) might adopt quite similar folds (***𝒜,*** *ℬ*), leading to their proximity in this high-dimensional space. Traditionally, fold space has been examined by hierarchically clustering domains based upon 3D structure comparison (Fig. 1A); in such approaches, whatever metric is used for the comparison can be viewed as organizing or structuring the space. The transition of a protein sequence from one fold to another, whether it be nearby (***𝒜*** ⟶ *ℬ*) or more distant (***𝒜*** ⟶ ***𝒞***), and be it naturally (via evolution) or artificially (via design/engineering), ikely occurs over multiple intermediate steps. These mechanistic steps include processes such as combining or permuting short secondary structural segments or longer regions (such as whole secondary structural elements [SSEs]), or mutating individual residues via nonsynonymous substitutions [5, 13–16]. In general, each such step may yield a new 3D structure, and that structure may correspond to the same or a different fold’. Similarities across these transitional states, ***𝒜*** ⟶ ***𝒜’*** ⟶ ***𝒜”*** ⟶ … ⟶ *ℬ*, blur the boundaries that delineate distinct groups: increasing or decreasing a relatively arbitrary and heuristic quantity, such as an RMSD or other similarity threshold, effectively alters the granularity of groupings in this space, and can change which structures belong to which groups. In this sense, the discrete versus continuous duality of fold space can be viewed largely as a matter of semantics or thresholding, versus any ‘real’ (intrinsic or fundamental) feature of the space itself [17].

**Fig. 1.**
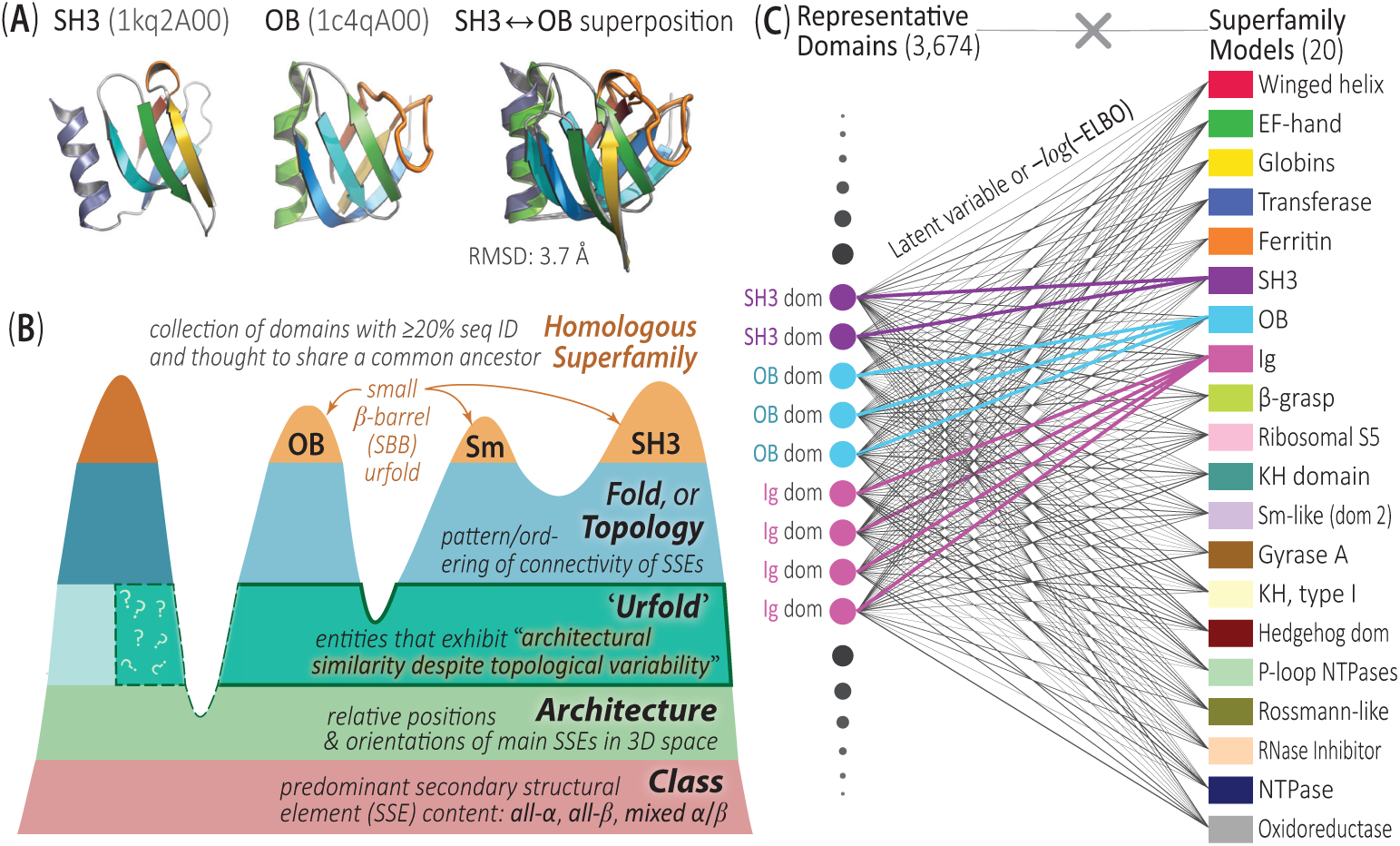
Overview of the Urfold model and DeepUrfold approach to identify domains that might reflect the phenomenon of “*architectural similarity despite topological variability* ”. (A) The SH3 and OB domains are prototypical members of the small *β*-barrel (SBB) urfold because they have the same barrel architecture, yet different strand topologies: though they have strikingly similar 3D structures and share extensive functional similarities (e.g., PPI binding on the same edge-strand, involvement in nucleic acid–binding and processing pathways [9, 10]), these similarities were historically obscured by the SH3 and OB superfolds having been classified differently. In the case of the SBB urfold, the loops linking the strands are permuted in the SH3 and OB (Fig. 2), yielding the different topologies seen in their 3D superposition. (B) If the Urfold phenomenon is viewed in terms of CATH, it is hypothesized to be a discrete structural entity or ‘level’ that lies between the Architecture and Topology strata, as schematized here. Note that entities from a finer-grained stratum (e.g., Topology) may impinge either partially or fully (‘?’ symbols) on a coarser-grained one (e.g., Architecture); the Urfold model unites the resultant discontiguous sets, thereby capturing similarities amongst proteins which are otherwise classified as different ‘folds’ (cf. Fig. 1 in [11]). (C) DeepUrfold, which applies deep learning to the Urfold conceptualization of protein structure, identifies potential new urfolds by creating 20 SF-specific VAE neural network models and comparing output scores (from inference calculations) for all representative domains from those SFs (totaling 3,674) to every other SF model; this process yields a fully-connected bipartite graph, as illustrated here. As a metric to compute initially, we can imagine comparing the latent variables (Fig. 3) from domain representatives using models trained on the same SF (colored lines in the graph). Then, all-by-all comparisons with all SFs can be performed to begin mapping fold space, which we view as being organized as mixed-membership communities, versus hierarchically-clustered, mutually-exclusive bins; as detailed below and shown in Fig. 4, such communities can be computed via stochastic block models (SBMs; reviewed in [12]).

Despite their limitations, it was pairwise similarity metrics in structure space that first indicated remote connections in a continuous fold space, the linkages potentially being mediated via shared fragments (see [18] and references therein). In an early landmark study, Holm & Sander [19] created an all-by-all similarity matrix from 3D structural alignments and discovered that the protein universe harbors five peptide ‘attractors’, representing frequently-adopted folding motifs (e.g., the *β*-meander).

Nearly a decade later, and armed with vastly more 3D structures, similar pairwise analyses across protein structure space showed that ‘all-*α*’ and ‘all-*β*’ proteins are separated by ‘*α/β*’ proteins [20]. All-by-all similarity metrics applied to full domains (or fragments thereof) can be equivalently viewed as a graph-theoretic adjacency matrix, thus enabling the creation of a network representation of fold space. Such networks have been found to be “nearly connected”, linking various domains (graph nodes) in ≈ 4-8 hops [21–23].

Graph-based representations of individual proteins have also motivated the study of common short (sub-domain) fragments. In pioneering studies, Harrison et al. [24, 25] found maximal common cliques of connected SSEs in a graph-based protein representation; their model took SSEs (helices, strands) as vertices and used the pairwise geometric relationships between SSEs (distances, angles, etc.) to decorate the graph’s edges. In that work, 80% of folds were found to share common cliques with other folds, and these were quantified by a new concept termed ‘*gregariousness*’ [24].

Although short, sub-domain–sized peptide fragments have been thoroughly studed, relatively few approaches have taken an evolutionary perspective, in the context of a continuous fold space. Goncearenco et al. [26] identified common loop fragments flanked by SSEs, called ‘*elementary functional loops*’ (EFLs), that couple in 3D space to perform enzymatic activity. Youkharibache [6] noticed that peptide fragments, called ‘*protodomains*’, are often composed (with *C*_2_ internal symmetry) to give a larger, full-sized domain. More recently, Bromberg et al. identified common fragments between metal-binding proteins using ‘*sahle*’, a new length-dependent structural alignment similarity metric [4]. These studies underscore the functional (and thus evolutionary) roles of sub-domain structural fragments.

Two state-of-the-art, evolution-based fragment libraries that are currently available, namely ‘*primordial peptides*’ [2] and ‘*themes*’ [27], involved creation of a set of common short peptide fragments based on HHsearch [28] profiles for proteins in SCOP and ECOD, respectively. The sizes of the libraries created by these two sequence-driven approaches (40 primordial peptides, 2195 themes) vary greatly, reflecting different stringencies of thresholds (and, ultimately, their different goals).

Another approach to study shared, commonly-occurring sub-domain fragments is to represent a protein domain as a vector of fragments. For example, the ‘*FragBag* ’ method [29] describes a protein by the occurrence of fragments in a clustered fragment library [30]. A recent and rather unique approach, ‘*Geometricus*’ [31], creates protein embeddings by taking two parallel approaches to fragmentation: (i) a *k*-mer– based fragmentation runs along the sequence, yielding contiguous segments, while (ii) a distance-based fragmentation uses the method of spatial moment invariants to compute (potentially non-contiguous) geometric ‘fragments’ for each residue position and its neighborhood within a given radius, which are then mapped to ‘*shape-mers*’. Conceptually, this allowance for discontinuous fragments is a key step in allowing an algorithm to bridge more of fold space, as similarities between such non-contiguous fragments can imply an ancestral (contiguous) polypeptide that duplicated and lost one or more *N ^′^*- or *C ^′^*-terminal SSEs, perhaps in a “creative destruction” process 5, 16] that yields two different folds (i.e., different topologies) despite the preserval of similar architectures.

### Limitations of Hierarchical Systems, and the Urfold

The conventional view of fold space as the constellation of all protein folds, grouped by their ‘similarities’ to one another, largely rests upon hierarchically clustering domains based upon 3D structure comparison, as exemplified in pioneering databases such as CATH [32], SCOP [33, 34], and ECOD [35]. Despite being some of the most comprehensive and useful resources available in protein science, these databases have intrinsic limitations that stem from their fundamental structuring scheme [36], reflecting assumptions and constraints of any hierarchical system (e.g., assigning a given protein sequence to one mutually exclusive bin versus others); in this design schema, domains with the same fold or superfamily (SF) cluster discretely into their own independent ‘islands’. The difficulty in smoothly traversing fold space, at least as it is construed by these databases—e.g., hop from island-to-island or create ‘bridges’ between islands in fold space—implies that some folds have no well-defined or discernible relationships to others. That is, we miss the weak or more indeterminate (but nevertheless *bona fide*) signals of remote relationships that link distantly-related folds. In addition to the constraints imposed by mutually exclusive clustering, the 3D structural comparisons used in building these databases generally rely upon fairly rigid spatial criteria, such as requiring identical topologies for two entities to group together at the finer (more homologous) classification levels (Fig. 1B). *What relationships might be detectable if we relax the constraints of strict topological identity?* As described below, this question is addressed by a recently proposed ‘Urfold’ model of protein structure [9, 11], which allows for sub–domain-level similarity.

Motivated by sets of striking *structure* ⟷ *function* similarities across disparate superfamilies, and even superfolds, we recently identified relationships between several SFs that exhibit architectural similarity despite topological variability (Fig. 1A), in a new level of structural granularity that allows for discontinuous fragments and that we have termed the ‘*Urfold* ’ (Fig. 1B; Supplementary Note 2; [9, 11]). Urfolds were first described in the context of small *β*-barrel (SBB) domains (Fig. 1A), based on patterns of *structure* ⟷ *function* similarity (as well as sequence signatures in MSAs, albeit more weakly) in deeply-divergent collections of proteins that adopt either the SH3/Sm or OB superfolds [9]. Notably, the SH3 and OB are two of the most ancient protein folds, and their antiquity is reflected in the fact that they permeate much of information storage and processing pathways (i.e., the transcription and translation apparatus) throughout all three domains of cellular life [16, 37, 38].

### DeepUrfold: Motivation & Overview

Can we rigorously ‘define’ the Urfold? The advent of deep learning [39], including the application of such approaches to protein sequences and structural representations [40, 41], affords opportunities to study protein interrelationships in wholly new and different ways—e.g., via quantitative comparison of ‘latent space’ representations of a protein in terms of lower-dimensional ‘embeddings’. Such embeddings can be derived from representations at arbitrary levels of granularity (e.g., atomic) and can subsume virtually any types of properties, such as amino acid type, physicochemical features (e.g., electronegatitivty), geometric attributes (e.g., surface curvature), phylogenetic conservation of sites, and so on. Two powerful benefits of such approaches are that (i) models can be formulated and developed in a statistically well-principled manner (or at least strive to be clear about their assumptions), and (ii) models have the capacity to be *integrative*, by virtue of the encoding (or ‘featurization’) of structural properties alongside phylogenetic, chemical, etc. characteristics of the data (in this case, augmenting purely 3D structural information about a protein). The methodology presented here explores the idea that viewing protein fold space in terms of feature embeddings and latent spaces (what regions are populated, with what densities, etc.)—and performing comparative analysis via such spaces (versus in direct or real’ 3D/geometric space)—is likely to implicitly harbor deep information about protein interrelationships, over a vast multitude of protein evolutionary timescales. Such distant timescales are likely to be operative at the Urfold level of structure [11].

Here, we present a deep learning–based framework, called ‘*DeepUrfold* ’, to systematically identify urfolds by using a new alignment-free, topology-agnostic, biochemically-aware similarity metric of domain structures, based on deep generative models, together with mixed-membership community detection algorithms. From a probabilistic perspective, our metric is rooted in the variational Bayesian inference that underpins variational autoencoders (VAEs [42]). From a deep learning perspective, our algorithm leverages embeddings and similarities in latent-space representations rather than simple (purely-geometric) 3D structures directly, enabling us to encode any sort of biophysical or other types of properties and thereby allowing more subtle patterns of similarities to be detected—such as may correspond to architectural simiarities among (dis-)contiguous fragments from different folds, or even superfolds, that are related only at great evolutionary distances (Fig. 1).

The DeepUrfold framework consists of four distinct methodological stages: *(**i**) Dataset Construction*, whereby protein 3D structures are prepared, featurized and allocated into suitable training/test splits for machine learning; *(**ii**) Training of SF-specific Models*, using featurized protein structural data and a hybrid 3D-CNN/VAE-based deep network; *(**iii**) All-by-all Inference Calculations*, computing VAE-derived ELBO-based scores to assess the ‘fit’ of each 3D structural domain representative to each SF, as schematized in Fig. 1C (i.e., subject each SF representative, *i*, to each SF-specific VAE model, *j*); *(**iv**) Elucidation of Any Community Structure* in these protein ↭ SF mappings, via stochastic block modelling (SBM) of the patterns of scores.

## Results

### The DeepUrfold Generative Modelling Framework

Conventionally, two protein structures that have similar architectures but varying topologies (i.e., folds) might be viewed as having resulted from convergent evolution. However, as in the case with the SH3 and OB superfolds, the *structure* ⟷ *function* similarities [9], and even *sequence* ⟷ *structure* ⟷ *function* similarities [16], can prove to be quite striking, suggesting that these domain architectures did not arise independently [6, 16] but rather as echoes of a (deep) homology. To probe what may be even quite weak 3D similarities, in DeepUrfold we model the evolutionary processes giving rise to proteins as an integrated 3D structure/properties ‘generator’. In so doing, we seek to learn probability distributions, *p*(*x θ*), that describe the specific geometries and physicochemical properties of different folds (i.e., features that largely define protein *function*), where the random variable *x* denotes a single structure drawn from (*x ɛ* **x**) a set of structures labelled as “having the same fold” (**x**), and *θ* denotes the collection of model parameters describing the variational distribution over the background (i.e., latent) parameters. We posit that folds with similar latent space embeddings and learned probabilistic distributions—which can be loosely construed as “structure ⟷ function mappings”, under our feature-set—likely have similar geometries/architectures and biophysical properties, regardless of potentially differing topologies (i.e., they comprise an urfold), and that, in turn, may imply a common evolutionary history.

Using the principles of variational inference [43], DeepUrfold learns the background distribution parameters ***θ_i_*** for superfamily distributions, i.e. models *p_i_*(*x_ij_|****θ_i_***), by constructing a variational autoencoder (VAE) model for each superfamily *i* (the VAE is trained on domain structures *j* annotated as being from superfamily *i*). In the current work, DeepUrfold is developed using 20 highly-populated SFs from CATH (see Fig. 1C, Supplementary Discussion 5.1 and Supplementary Table 1). The original/underlying likelihood distribution, *p_i_*(*x_ij_|****θ_i_***), is unknown and intractable, but it can be estimated by considering an easier-to-approximate posterior distribution of latent space parameters, *q_i_*(*z_ij_|***x***_i_*), where *z* denotes the latent variables we wish to infer and, again, **x** is our data (protein structures); in our case, the approximating distribution *q*(*z|***x**) is taken as sampling from a Gaussian. To ensure that *q_i_*(*z_ij_|***x***_i_*) optimally describes *p_i_*(*x_ij_|****θ_i_***), one can seek to maximize an *evidence lower bound* (ELBO) quantity as a variational objective, which supplies a lower bound of the marginal log-likelihood of a single structure *j*, ln[*p_i_*(*x_ij_*)]. The ELBO inequality can be written as:

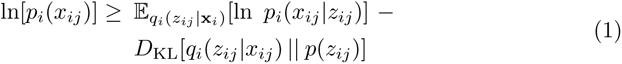

where *p_i_*(*x_ij_*) is the likelihood, E is the expectation value of *q* in terms of *p*, and *𝒟*_KL_[*q p*] is the Kullback-Leibler divergence, or relative entropy, between the two probability distributions *q* and *p*. In other words, maximizing the ELBO corresponds to maximizing the expected log-likelihood of our learned model and minimizing the entropy or ‘distance’ (*𝒟*_KL_) between (i) the true/exact underlying prior distribution of the data given a model, *p*(*x|****θ***), and (ii) our learned/inferred approximation, as a posterior distribution of latent parameters given the data, *q*(*z|***x**). Pragmatically, DeepUrfold’s variational objective is formulated as a minimization problem (Supplementary Discussion 5.2), so we compute –(ELBO) values (Supplementary Note 3).

In a similar vein, part of DeepUrfold’s testing and development (detailed below) involved training ‘joint models’ using a bag of SFs with intentionally *different* topologies—e.g., a mixed SH3 OB set, as well as a single ‘Global Model’ that was trained across all 20 SFs. The training of these mixed models took into account the *class imbalance* [44, 45] that stems from there being vastly different numbers of available 3D structural data for different protein SFs (e.g., immunoglobulin [Ig] structures, which are disproportionately abundant). Further details of the multi-loop permutation analyses used to test and develop DeepUrfold, particularly as regards –(ELBO) distributions and potential class imbalance issues, can be found below (e.g., Fig. 2) and in Supplementary Figs. 9-10. In particular, Supplementary Fig. 10 considers three alternative approaches to addressing class imbalance in a joint SH3 OB model, by examining the impacts on –(ELBO) distributions of training models with either (i) over-sampling of SH3s (to match the number of OBs), (ii) under-sampling of OBs (to match the number of SH3s), or (iii) default(/imbalanced) sampling, wherein all SH3 and OB domains are used. Intriguingly, class imbalance was not found to be an issue n the case of training these joint SH3 OB models (recall that the SH3 and OB comprise the putative SBB urfold [9]).

**Fig. 2.**
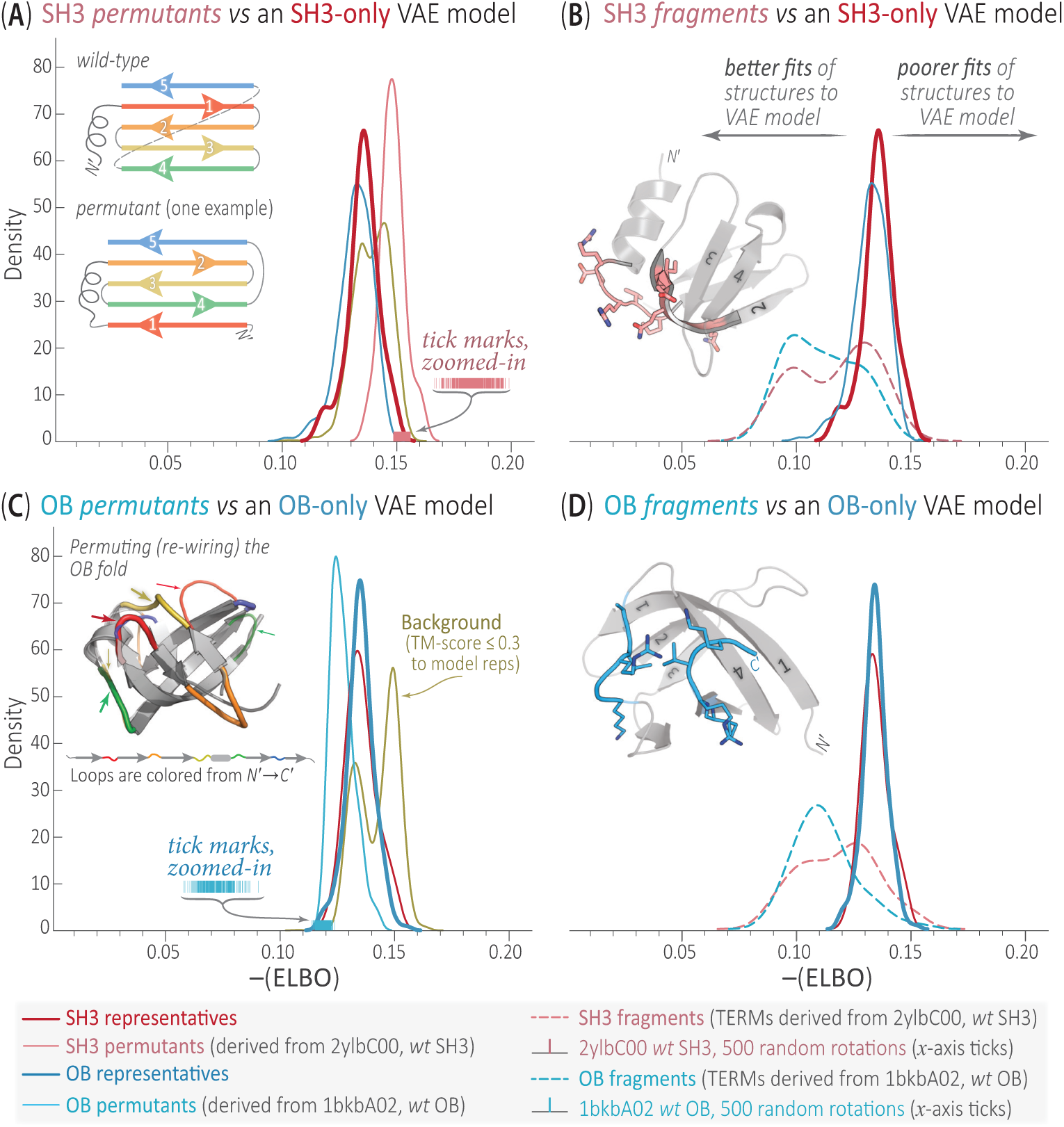
Likelihood-based ELBO values can quantify similarities among multi-loop permuted structures and can discern structural fragments. To gauge the sensitivity of Deep-Urfold’s VAE-based metric to loop orderings (topology), we generated a series of fictitious folds, both SH3-based (A, B) and OB-based (C, D), and analyzed their patterns of scores. Specifically, we implemented a multi-loop permutation algorithm [46] to scramble the SSEs in wild-type SH3 (2ylbC00) and OB (1bkbA00) domains, ‘rewiring’ the loops by stitching together SSEs and energetically relaxing the resultant structures. Each permuted structure was subjected to a VAE model trained on either an (A, B) SH3-only or (C, D) OB-only domain set. (Joint SH3 *∪* OB models can also be trained [Supp. Fig. 10].) Fits to trained models were approximated by the –(ELBO) score, viewed here as a similarity or ‘goodness-of-fit’ measure between an individual structure and the SF-level VAE. In reference to a given model, a given permutant or fragment query structure with a –(ELBO) value smaller (less positive) than the corresponding wild-type structure for that model can be considered as structurally more ‘similar’ (a better fit; see arrows in B) to the consensus model (Supp. Figs 8, 10; Supp. Discussion 5.2). To compare the ELBO-based score distributions of permutants to the corresponding wild-types, and to demonstrate the rotational invariance of the trained models, we subjected 500 random rotations of each wild-type SH3 and OB domain to inference passes through its corresponding VAE model, yielding the sets of *x*-axis tick marks; the tightness of these distributions is consistent with rotational invariance of DeepUrfold models. ELBO-based score distributions for structurally-dissimilar domains (TM-Score *≤* 0.3 to any SH3 or OB domain representatives), which we call the ‘background’ (gold traces), are included here to assess if a particular model is distinguishable from the structurally random background. We show how well the (B) SH3-only and (D) OB-only VAE models score potentially discontiguous sub-domain fragments, as captured by ‘tertiary structural motifs’ (TERMs) derived from each wild-type SH3 and OB domain structure. These broad fragment traces are leftward-shifted for both the SH3 and OB cases, implying better fits to the consensus VAE models.

As input to a DeepUrfold VAE, we encode the 3D structure of a protein domain by representing it as a 3D volumetric object (Supplementary Fig. 1), akin to the nput used in 3D convolutional neural networks (CNNs). Indeed, DeepUrfold’s neural network architecture can be viewed as a hybrid/stacked 3D-CNN–based VAE. In our discretization, atoms are binned into volumetric elements (voxels), each of which can be tagged or labeled, atom-wise, with arbitrary properties (biophysical, phylogenetic, etc.). A critical point is that this representation scheme is agnostic of polypeptide chain topology, as the covalent bonding information between residues, and the order of SSEs, is not explicitly retained; note, however, that no information is lost by this representation, as such information is implicit in the proximity of atom-occupied voxels n a model (and can be used to unambiguously reconstruct a 3D structure). The above preparatory and featurization steps utilized ‘`Prop3D`’, a computational toolkit that we have developed for machine learning with protein structures [47].

Note that we do not use VAEs to generate new samples from a given SF per se (Supplementary Note 6). Rather, the role of the VAE in DeepUrfold can be viewed as that of an anomaly detection tool, to robustly and quantitatively address the question: “Based on learned, superfamily-specific latent space representations, what is the ikelihood that a given domain structure (from any SF, *i*) arose from (or, alternatively, *was generated by*) a particular SF-specific VAE model, *j*?”. Does such information lluminate the types of protein similarities that DeepUrfold can sensitively detect?

### DeepUrfold Models Are Sensitive to Topological Variation

Can DeepUrfold models detect similarities amongst topologically-distinct, architecturally-similar proteins? To initially assess our DeepUrfold VAE models, and to examine the properties of the Urfold model more broadly, we directly probed the Urfold’s core concept of “architectural similarity despite topological variability” [11]. These tests were performed by considering sets of artificial protein domains that have identical architectures but with specifically introduced loop permutations. We obtained these systematically engineered perturbations of a 3D structure’s topology by ‘rewiring’ the SSEs (scrambling the loops), while retaining the overall 3D structure/shape (i.e., architecture). The SH3 and OB folds, which are related by a (non-circular) permutation, offer a rich system for these tests, as shown in Fig. 2 and delineated in [9]. Specifically, (i) we systematically created permuted (fictitious) 3D structures startng with representative SH3 and OB domains (Supplementary Fig. 9a) via structural modeling (including energetic relaxation), and (ii) we then subjected each of these rewired structures, in turn, to DeepUrfold models that were trained on either the SH3 alone (Fig. 2A), OB alone (Fig. 2C), or a joint SH3 OB set (Supplementary Fig. 10). The SH3/Sm and OB superfolds comprise the first-identified urfold [9], namely the small *β*-barrel (SBB). While SBBs typically have six SSEs (five strands and a helix), there are four ‘core’ *β*-strands, meaning an SBB’s *β*-sheet can theoretically adopt one of at least 96 distinct loop permutations [9]; note that, based on the operational definitions/usage of the terms ‘topology’ and ‘fold’ in systems such as SCOP, CATH, etc., such engineered permutants almost certainly would be annotated as being from different homologous superfamilies, implying no evolutionary relatedness. Thus, the loop-scrambling approach described here is a systematic way to gauge DeepUrfold’s ability to discern similarities at the levels of architecture and topology, in a self-contained manner that is agnostic of preexisting classification schemes such as CATH.

In general, we find that the synthetic/permuted domain structures have similar distributions of –(ELBO) scores as the corresponding wild-type domains (Fig. 2), though there are subtleties: SH3 permutants against an SH3-only model are shifted to poorer fits (Fig. 2A; though not so in the context of a joint SH3 OB model [Supplementary Fig. 10]), while for the OB the permutants are shifted to better fits relative to ts representatives (Fig. 2C). Those rewired structures with –(ELBO) scores smaller (less positive) than the wild-type indicate that those permutants better fit the VAE model than does the wild-type domain itself (i.e., distributions that shift leftward in Figs 2A, C, and Supplementary Fig. 10). In a sense, these permuted variants can be interpreted as being more compatible (structurally, biophysically, etc.) with the DeepUrfold-learned variational model (a ‘consensus’ model, of sorts [Supplementary Note 4]), and thus perhaps more thermodynamically stable or structurally robust were they to exist in reality—an interesting possibility as regards protein design and engineering. In terms of more conventional structural similarity metrics, the TM-scores [48] for permuted domain structures against the corresponding wild-type topolog (Supplementary Fig. 9a) were typically found to lie in the range ≈ 0.3−0.5—i.e., values which would indicate that the permutants and wild-type are not from identical folds, yet are more than just randomly similar (Supplementary Fig. 9b).

Another approach to explore potential similarity amongst proteins that exhibit architectural similarity despite topological variability is to examine the behavior of small domain fragments, such as short ‘tertiary structural motifs’ (‘TERMs’ [49, 50]), under the DeepUrfold model. Therefore, TERMs for representative SH3 and OB domains were computed and subjected to both the SH3-only and OB-only VAE models, with the structures of the highest-scoring fragments shown as insets in Fig. 2B, D. Many fragments derived from representative SH3 and OB domains scored better than the full, extant domain structures from which they were derived (smaller –(ELBO) scores), indicating DeepUrfold models may be able to identify shared common fragments (Fig. 2B, D), at least in terms of the ELBO-based scoring metric used here. Note that for both the SH3 and OB models the –(ELBO) distributions of these TERM fragments are left-shifted (i.e., less positive values, better fits to the VAE model), as would be intuitively expected were one measuring the ‘fit’ of a fragment to a larger-sized entity (e.g., a full domain; an underdetermined problem), versus fitting a domain-sized entity to a VAE model derived from training on domain-sized structural data.

The findings from the loop-permutation tests and fragment (TERMs) calculations suggest that the DeepUrfold model can be used in a topology-insensitive, architecture-sensitive approach: Note, for example, (i) the extensive overlap between the distribution of OB representative scores (blue curve) when subjected to the SH3-only trained VAE model (red curve, Fig. 2A), and, complementarily, (ii) the extensive overlap between the SH3 representatives (red curve) under the OB-only VAE model (Fig. 2C). In short, the differing SH3 and OB topologies behave similarly to one another (and different from the background) under the DeepUrfold VAE models shown in Fig. 2 as well as the joint SH3 ∪ OB models in Supplementary Fig. 10. Moreover, putative fragments of the SH3 and OB folds (TERMs) behave in an expected manner, in terms of their –(ELBO) distributions being broadened and shifted leftwards in Fig. 2B, D). We suspect that these behaviors are possible because, as mentioned above, our encoding is intentionally agnostic to topological connectivity information and, rather, is sensitive only to 3D shape/architecture. The generality of this approach is useful in detecting similarities amongst sets of seemingly dissimilar 3D structures—and thereby identifying specific candidate urfolds—because two sub-domain portions from otherwise rather (structurally) different domains may be quite similar to each other, even if the domains of which they are a part have different (domain-level) topologies but identical overall architectures. This concept can be represented symbolically: for an arbitrary subset of SSEs, d, drawn from a full domain D, the Urfold model permits relations (denoted by the ‘∼’ symbol) to be detected between two different ‘folds’, *i* and *j* (i.e. *d_i_* ∼ *dj)*, at the sub-domain level, without requiring that the relation also be preserved with the stringency of matched topologies at the higher ‘level’ of the full domain. That is, *d_i_* ∼ *d_j_* ⇏ *𝒟_i_* ∼ *𝒟_j_*, even though *d_i_* ⊂ *𝒟_i_* and *d_j_* ⊂ *𝒟_j_* ; this is in contrast to how patterns of protein structural similarity are traditionally conceived, at the domain level (and with *d_i_* ∼ *d_j_* ⇒ *𝒟_i_* ∼ *𝒟_j_*). Here, we can view the characteristic stringency or ‘threshold’ level of the Urfold, ‘*d*’, as being nearer that of architecture (Fig. 1B), while *𝒟* reflects both architecture and topology (corresponding to the classical usage of the term ‘fold’).

### Superfamily Latent Spaces and the Continuity of Fold Space

Do DeepUrfold’s latent-space representations capture gross structural properties across superfamilies and, if so, are there implications for the continuity of fold space? We find that the latent space of each superfamily-level DeepUrfold model offers a new, nuanced view of that superfamily, and examining patterns of similarities among such models (e.g., via a graph such as in Fig. 1C) may offer a uniquely informative view of fold space. Each SF-specific model captures an amalgamation of different 3D geometric and physicochemical properties—which uniquely characterize that individual SF, versus others—as a single ‘compressed’ data point or embedding; in this way, the learned latent space representation (or ‘distillation’) is more comprehensible than is a full 3D domain structure (or superimpositions thereof). In a sense, the DeepUrfold approach—and its inherent latent space representational model of protein SFs, with featurized proteins—can reconcile the dichotomy of a continuous versus discrete fold space because the Urfold model (i) begins with no assumptions about the nature of fold space (i.e., patterns of protein interrelationships), and (ii) does not restrictively enforce full topological ordering as a requirement for a relation to be detected (even a rather weak one) between two otherwise seemingly unrelated domains (e.g., 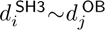 is not forbidden, using the terminology introduced above). We posit that DeepUrfold can detect these weak similarities (i.e., exhibit high sensitivity) because it operates on protein domains that are featurized beyond purely 3D spatial coordinates; our rationale here is that molecular evolution acts on proteins holistically, not on merely their 3D geometries, patterns of electrostatic charges, or any other single attribute taken in solation.

Can the DeepUrfold framework help us ‘visualize’ fold space? A rigorous analysis of fold space, through the lens of DeepUrfold, can begin by constructing an all-by-all bipartite graph of 3,674 domain representatives × the 20 most-populated CATH SFs, as schematized in Fig. 1C. While that dataset is the basis for our further analyses of fold space, using SBMs (below), we can also seek to visually explore the latent space representations learned in DeepUrfold’s VAE models. As an initial step in exploring DeepUrfold’s learned representations, we analyzed the latent spaces of representative domains for highly populated SFs by mapping the latent space embeddings into two dimensions, including the use of various schemes to combine/aggregate embeddings (Fig. 3, Supplementary Figs. 11-25, Supplementary Discussion 5.3). Proteins that share similar geometries and biophysical properties should have similar embeddings, and would be expected to lie close together in DeepUrfold’s latent-space representation, regardless of the annotated ‘true’ SF. Though this initial picture of the protein universe is limited to 20 highly populated CATH SFs, in this work (Supplementary Table 1, Supplementary Discussion 5.1), already we can see that these SF domains appear to be grouped and ordered by secondary structure composition (Fig. 3)—a result that is consistent with past analyses which used approaches such as multidimensional scaling to probe the overall layout of fold space (e.g., [20]). Variable degrees of intermixing between SFs can be seen in UMAP projections such as illustrated in Fig. 3—at the evel of an individual SF-trained model (the SH3-only VAE in Fig. 3A), for a Concatenated Model (Fig. 3B), and for a Global Model (Fig. 3C). This is a compelling finding, with respect to the Urfold and its relaxed notion of allowing for intermixed superfamilies [11]. In addition to this mixing, the latent space projections generally are not punctate, at the SF-level (Fig. 3A, Supplementary Fig. 18): rather than consist of clearly demarcated, well-separated ‘islands’, we instead find that they are fairly compact’ (in a loose mathematical sense) and well-connected, with only a few disjoint outlier regions. Manual inspection of the outlier domain structures shows that some of them are incomplete sub-domains or, intriguingly, a single portion of a larger domain-swapped region [51]. Together, these findings support a rather continuous view of fold space, at least for these 20 exemplary superfamilies.

**Fig. 3.**
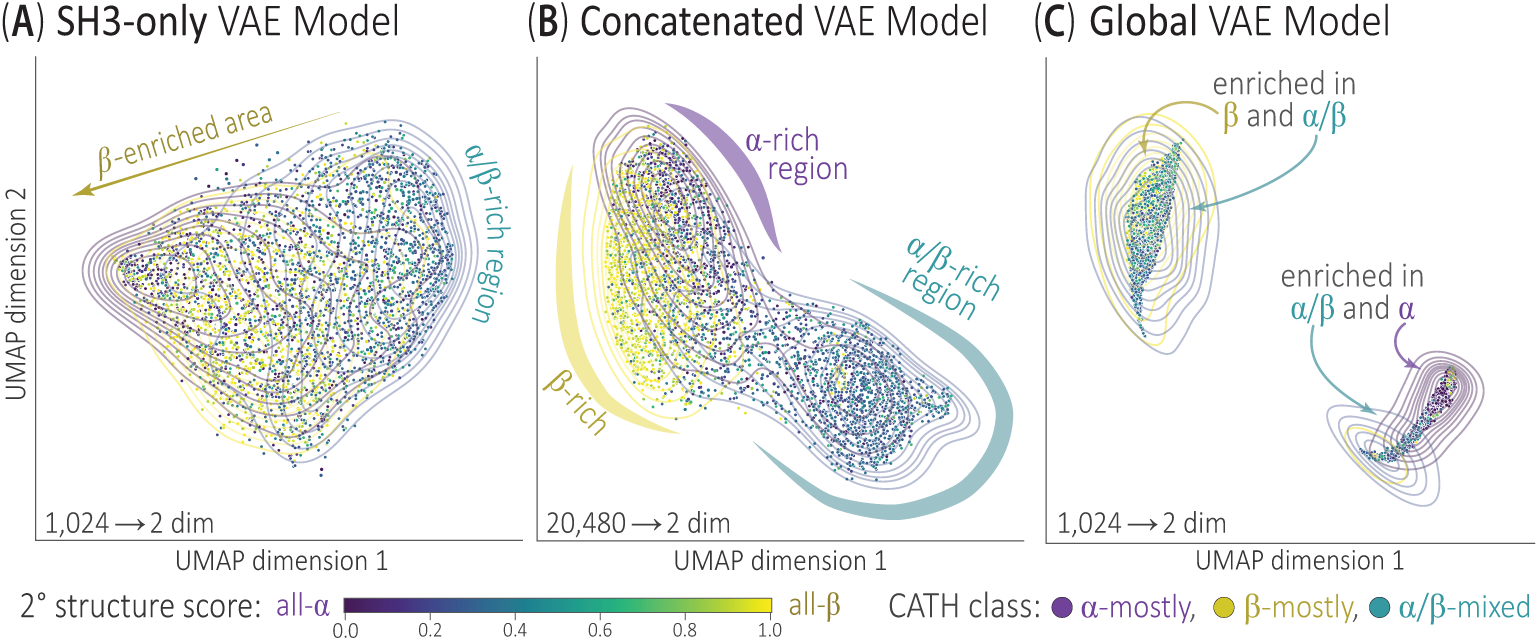
Dominant variables of DeepUrfold’s latent-space models capture gross structural properties and indicate a highly continuous fold space. In a pilot study of fold space, we used DeepUrfold to develop 20 distributions/models for 20 CATH homologous SFs. Representatives from each SF were subjected to VAEs that were pre-trained on domains from the same SF, and then the latent space variables for each structural domain were examined via the uniform manifold approximation and projection (UMAP) method; this process reduced the 1,024 dimensions of the actual domain-level VAE models to the two-dimensional UMAP projections shown here (and, for all 20 SFs, in Supplementary Fig. 18). In these diagrams each domain is colored by a secondary structure score; computed as 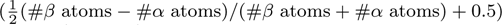, this score ranges from zero (for all-*α*) to unity (for all-*β*). As a visual aid, kernel density estimates (isodensity contour lines) surround domains belonging to the same annotated CATH *Class*. Our exploratory visualization considered (A) embeddings from an ‘SH3-only’ VAE model, as well as latent space representations obtained via (B) a ‘Concatenated Model’ aggregation strategy, whereby embeddings for each domain from the 20 SF VAE models were concatenated in the feature dimension (note the input dimensionality of 1,024 *×* 20 = 20,480), and (C) embeddings from a monolithic ‘Global Model’ VAE, derived by training on a pooled set of domains from all SFs. In each of these visualizations the protein domains, as learned by DeepUrfold, can be seen to largely group together by secondary structure composition; moreover, some of these regions are roughly ordered in panels A, B and C. For instance, distinct regions of mostly-*α*, mostly-*β* and *α/β* domains are clear in (B), with some overlap. Similarly, a gradient of domain classes (with more extensive overlap) can be seen in the SH3-only VAE in (A). Intriguingly, a far more disperse/punctate layout occurs for the Global Model in (C).

While each superfamily model is trained independently, with different sets of domain structures (SH3, OB, etc.), we find that the distributions that the VAE- based SF models each learn—again, as ‘good’ approximations to the true likelihood, *p_i_*(*x_ij_* ***θ_i_***)—are similar, in terms of the dominant features of their latent spaces. In other words, the multiple VAE models (across each unique SF) each learn a structurally ow-level, ‘coarse-grained’ similarity that then yields the extensive overlap seen in the SH3-only VAE model in Fig. 3A. When colored by a score that measures secondary structure content, for many SFs there are clear directions along which dominant features of the latent-space can be seen to follow, as a gradient from ‘all-*α*’ domains to ‘all-*β*’ domains, separated by ‘*α/β*’ domains. These findings are consistent and reassuring with respect to previous studies of protein fold space (e.g., [20]), as well as the geometric intuition that the ‘similarity’ between two domains would roughly track with their secondary structural content (e.g., two arbitrary all-*β* proteins are more likely to share architectural similarity than would an all-*β* and an all-*α*).

Can further information be gleaned via other visualizations of DeepUrfold’s embeddings? We obtained an initial view of fold space, in the DeepUrfold framework, from a single-superfamily model—namely, the SH3-only VAE (Fig. 3A). Extending this, we created 20 different views, one for each of the top 20 CATH superfamilies; as a supplement to the illustrative SH3-only data (Fig. 3A), Supplementary Fig. 18 provides this comprehensive suite of SF-level embeddings for all 20 CATH SFs. In addition, other latent space visualization results were obtained by, e.g., coloring domains by physicochemical properties, conducting dimensionality reduction via PCA and t-SNE, etc. (Supplementary Figs. 15-17). In order to create views of fold space that incorporate all 20 SF-specific models into a *single* representation, we pursued experiments wherein we ‘combined’ embeddings from the different superfamily-specific VAEs in various ways, taking into account that each embedding occupies a potentially quite different mathematical space. We did so via several approaches: (i) by subjecting all domain representatives to inference passes through each of the 20 superfamily models and concatenating the embeddings for each domain from each model into a single, ultra-wide embedding (Fig. 3B’s ‘Concatenated Model’); (ii) by developing a monoithic ‘Global Model’ VAE trained on a set of all domains from every superfamily, accounting for class imbalance (Fig. 3C); and (iii) by using an Optimal Transport (OT) algorithm to align each embedding, before concatenation as in (i) (a UMAP reduction for OT-aligned embeddings is in Supplementary Fig. 19).

Taken in its entirety—in Fig. 3, Supplementary Figs. 11-25, and Supplementary Discussion 5.3—two themes emerge in our exploratory views of DeepUrfold-derived protein embeddings: (i) domains are found to coalesce into different regions of latent space, roughly delineated by secondary structural content (i.e., CATH classes), and (ii) significant continuity exists between these regions. How extensive are the observed clusterings, and what do they imply about protein relationships?

### Protein Interrelationships Defy Discrete Clusterings

Our initial finding that fold space is rather continuous, at least under the DeepUrfold model, implies that there are, on average, webs of interconnections (similarities, relationships) between a protein fold and its neighbors in fold space (*𝒜^′^*, *𝒜^′′^*, *ℬ, …*). Therefore, we posit that an optimally realistic view of the protein universe will not entail hierarchically clustering proteins into mutually exclusive bins, regardess of whether such binning is based upon their folds (giving *fold space*) or any other relatively simple (standalone) geometric feature/criterion. Alternatives to discrete clustering could be such approaches as *fuzzy clustering*, *multi-label classification*, or *mixed-membership community detection* algorithms. DeepUrfold’s strategy is to detect communities of similar protein domains, at various levels of stringency, based on the quantifiable similarities of their latent-space representations (versus, e.g., hierarchical clustering based on RMSD or other purely-geometric measures). Again, this s possible because we are armed with a battery of ELBO-based scores (Fig. 2) of the fit’ of each SF domain representative to each of the top 20 SF VAE models (Fig. 1C).

In DeepUrfold, we formulate this labeling/classification/grouping task as a problem n nonparametric Bayesian stochastic block modelling (SBM; [52, 53]). In particular, we fit an edge-weighted [54], degree-corrected, mixed-membership [55, 56], hierarchical [57] SBM to a fully connected bipartite graph that is built from the similarity scores between (i) the VAE-based SF-level models (one side of the bipartite graph) and (ii) representative structural domains from these SFs (the other side of the bipartite graph), as schematized in Fig. 1C. We chose to capture the ‘fit’ between a domain representative and a particular SF’s VAE model by weighting each edge by the quantity —log (—(ELBO)) (see Fig. 1C, Eq 1, Supplementary Fig. 8, and Supplementary Discussion 5.2). The motivation for this approach is that the full, global collection of –(ELBO)-weighted protein interrelationships (again, between SFs and domain representatives) most naturally corresponds to a bipartite graph, or network, which can be represented by its adjacency matrix, ***A****_d×sfam_*; this matrix features covariate edge weights ***x*** that link vertices from the two ‘sides’ of the bipartite graph, where *sfam* ɛ 20 highly-populated SFs and *d* ɛ 3,674 representative domains from the 20 SFs. Following Peixoto [54], we can write the full joint probability of a given bipartite graph/network occurring by chance—with precisely the same vertices connected by the same edges, with the same weights—as the following product over distributions of data and model parameters:

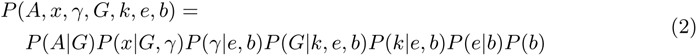

where ***b*** is the overlapping partition that represents the numbers of blocks (protein communities) and their group memberships (which nodes map to which blocks), ***e*** is a matrix of edge counts between the groups (thus allowing for mixed-membership between blocks), ***k*** is the labelled degree sequence, and ***G*** is a tensor representing the labeling of half-edges (each edge end-point *r, s*) to account for mixed-membership, satisfying the constraint 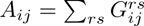 . The edge covariate parameters ***x*** (e.g., ELBO-based scores) are sampled from a microcanonical distribution, *P* (*x|G, γ*), where *γ* imposes a hard constraint such that 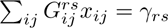 (Sec. VIIC of [52] and personal communication with T. Peixoto). We seek an SBM that best captures ***A***, where ‘best’ is meant as the usual trade-off between model accuracy (to the observed data) and model simplicity (i.e., mitigating overparametrization). An optimal SBM is obtained by considering this as a nonparametric Bayesian inference problem, meaning that (i) model features (the number of groups/blocks, node membership in blocks, patterns of edges between nodes and between groups, etc.), as well as (ii) model parameters and hyperparameters that are sampled over (marginalized out, via integration), are not set *a priori* but rather are determined by the data itself.

We estimate the optimal parameters for a given SBM via Markov chain Monte Carlo (MCMC) methods. Several different models are created for different ***b*** and ***e*** in order to find the optimal number of blocks with overlapping edges between them, and these are evaluated using a posterior odds-ratio test [55, 56].

Armed with the above SBM methodology, we can now summarize DeepUrfold’s overall approach as consisting of the following four stages: (i) Dataset construction, e.g. via the aforementioned discretization of 3D structures and biophysical properties into voxelized representations [47]; (ii) Training of SF-specific models, using our hybrid stacked 3D-CNN/VAE-based deep networks; (iii) In an inference stage, calculation of ELBO-based scores for ‘fits’ obtained by subjecting SF representative *i* to the VAE models of all other SFs, *j*; (iv) To decipher any patterns amongst these scores, utilization of SBM-based analysis of ‘community structure’ within the complete set of similarity scores for the VAE-based SF-level models (i.e., the full bipartite network of Fig. 1C, SF*_i_ ×* model*_j_*).

Application of this DeepUrfold methodology to the 20 most highly-populated CATH superfamilies identifies many potential communities of domain structures and SFs (Fig. 4, Supplementary Fig. 26), as well as other interesting patterns amongst electrostatic potentials, partial charges, and so on (Supplementary Figs. 27-32). Subjecting all domain representatives to all 20 SF-specific models, in an exhaustive *all* _SF-models_ × *all* _SF-reps_ analysis, reveals the global community structure shown in Fig. 4A (cf. CATH groupings in Fig. 4B). We argue that two proteins drawn from vastly different SFs (in the sense of their classification in databases such as CATH or SCOP) can share other, more generalized (e.g., non-contiguous) regions of geometric/structural and biochemical/biophysical properties, beyond simple permutations of secondary structural elements. And, we believe that the minimally-heuristic manner in which the DeepUrfold model is constructed allows it to capture such ‘distant’ linkages. In particular, these linkages can be identified and quantitatively described as patterns of similarity in the DeepUrfold model’s latent space. Organizing protein domains and superfamilies’ based on this new similarity metric provides a new view of protein nterrelationships—a view that extends beyond simple structural/geometric similarity, towards the realm of integrated *sequence ↔ structure ↔ function* properties.

**Fig. 4.**
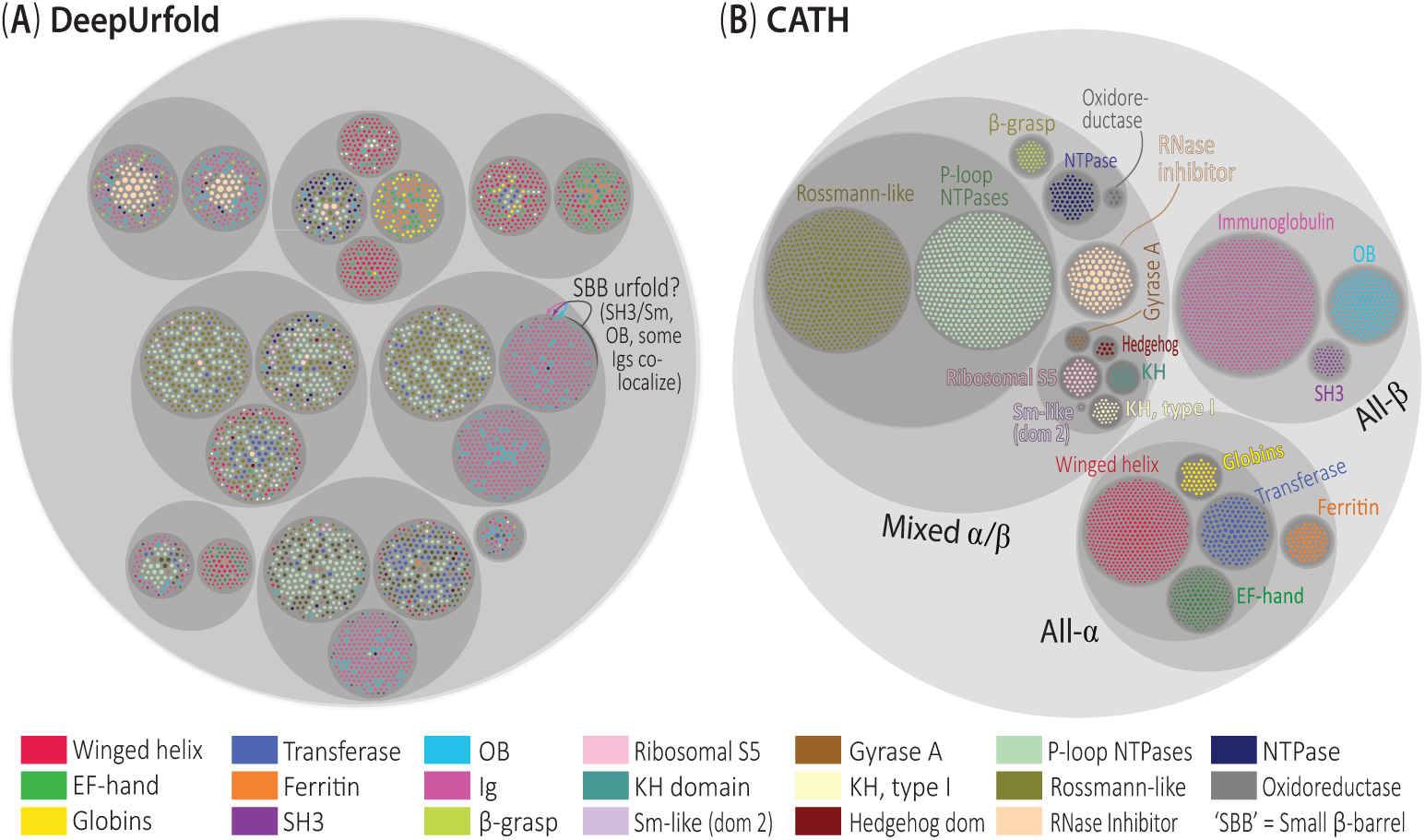
Protein interrelationships defy discrete clusterings: Stochastic block modelling of an all-vs-all comparison of domain structures and superfamily models. Here, we depict (A) the SBM communities predicted by DeepUrfold as a circle packing diagram, following a simlar representational scheme as for (B) the CATH hierarchy. While DeepUrfold avoids hierarchical clustering in favor of mixed-membership communities, we nevertheless display the groupings in this manner for the sake of visual representation and to facilitate comparison to CATH. Each domain representative is drawn as an innermost circle (corresponding to leaves in a hierarchical tree), colored by the annotated CATH SF and sized by the number of atoms. All of the SF-labelled nodes were ound to cluster together (Supplementary Fig. 26) and therefore were removed from the list used for this representation. Note that many SH3 and OB domains lie within the same lowest-level communities (labeled ‘SBB urfold?’ in (A)), showing that DeepUrfold can detect the link between these folds, as posited in the Urfold model [11]. Indeed, comparison of the patterns of groupings in (A) to the CATH hierarchy in (B) reveals that DeepUrfold is learning a rather different, non-hierarchical and ntermixed map of protein relationships.

We find that domains that have similar –(ELBO) scores against various superfamily models (other than the SF against which they were trained) are more likely to contain important biophysical/biochemical properties at particular—and, presumably, functionally important—locations in 3D space; these consensus regions/properties can be thought of as ‘defining’ the domain (Supplementary Note 5). Furthermore, if two domains map into the same SBM community, it is likely that both domains share the same scores when run through each SF model (i.e., an inference calculation), so we hypothesize that that community might contain an urfold that subsumes those two domains (again, agnostic of whatever SFs they are labeled as belonging to in CATH or other databases). We also expect that some domains (those which are particularly ‘gregarious’?) may be in multiple communities, which may reflect the phenomenon of a protein being constructed of a multifarious ‘urfold’ or of several sub-domain elements. Because of the conceptual difficulties and practical complexities of analyzing, visualizing and otherwise representing such high-dimensional data, in the present work we show only the single most likely cluster that each protein domain belongs to (e.g., in Fig. 4), while emphasizing that multi-class membership is a key element of DeepUrfold’s approach.

Given the stochastic nature of SBM methods, we computed six different replicates and chose the single most likely clustering for visualization. Fig. 4 displays the specific replica with 20 SFs and highest overlap score compared to CATH (simply to facilitate comparison to and reference to CATH). While each replica yielded slightly different hierarchies and numbers of clustered communities (ranging from 19-23), the communities at the lowest/finest level remained reassuringly consistent; in addition, we computed a ‘downsampled’ SBM in order to assess potential artefacts due to highly imbalanced classes, and found this not to be an issue (Supplementary Fig. 33). Overall, DeepUrfold’s SBM groupings exhibit varying degrees of intermixing. Specifically, in each replicate (see above) the SH3 and OB clustered into the same communities, as did the Rossmann-like and P-loop NTPases (Fig. 4A), instead of exclusively occupying their own individual clusters as in CATH (Fig. 4B). That members of the SH3 & OB superfolds aggregate in the same intermixed communities (and likewise for Rossmann folds & P-loop NTPases) is consistent with the Urfold model, as predicted for these specific SFs via manual/visual analysis [9, 11]. In general, each DeepUrfold community can be seen to contain domains from distinct CATH SFs (Fig. 4A, Supplementary Fig. 33), again conforming with the Urfold view of protein structural relationships. Most broadly, for the 20 protein SFs treated here we see that ‘mainly-*α*’ and ‘*α*/*β*’ domains preferentially associate, while ‘mainly-*β*’ and ‘*α*/*β*’ group together too (Fig. 4); interestingly, this result echoes the UMAP reduction of our Global Model, wherein domains were seen to aggregate into two distinct ‘*α* + *α*/*β*’ and ‘*β* + *α*/*β*’ islands (Fig. 3C).

In addition to coloring each domain (node) by its preexisting CATH SF label in circle-packing diagrams, such as that of Fig. 4, we also explored coloring domain nodes by other basic types of molecular properties. These additional properties include (i) secondary structure type, (ii) average electrostatic potential, (iii) average partial charge, and (iv) enriched gene ontology (GO) terms (Molecular Function, Biological Process, and Cellular Component), as shown in Supplementary Figs. 27-32. A navigable, web-based interface for exploring these initial DeepUrfold results is freely available at https://bournelab.org/research/DeepUrfold/. Interestingly, domains with simiar average electrostatic potentials and partial charges can be found to associate into similar groups in DeepUrfold (Supplementary Figs. 27a and 28a), whereas the corresponding CATH-based circle-packing diagrams, when colored by those same features, have no discernible order or structuring (Supplementary Figs. 27b and 28b); whether or not this phenomenon stems from any underlying, functionally-relevant ‘signal’ is a question of interest for further work.

In order to assess how ‘well’ our DeepUrfold model does, we compare and contrast our clustering results with CATH. However, we emphasize that there is no reliable, objective ground truth for a map of fold space, as there is no universally-accepted, correct’ description of fold space (and, it can be argued, even ‘fold’). Therefore, we cautiously compare our DeepUrfold results to a well-established system, like CATH, with the awareness that these are two conceptually different approaches to representing and describing protein structure relationships and, thus, the protein universe. Indeed, because our model uses a fundamentally different input representation of proteins, intentionally ignoring all topological/connectivity information, we expect that our model will deviate from CATH in terms of clustering-related measures such as *completeness*, *homogeneity*, *silhouette score*, and *partition overlap* [56]. Given all this, approaches that do differ from CATH—versus matching or recapitulating it—can be considered as representing an alternative view of the protein universe. Somewhat counterintuitively, we deem weaker values of our comparison metrics (e.g., less similarity to CATH) as providing stronger support for the Urfold model of protein structure. Simultaneously, we systematically compared how well other, independently-developed sequence– and structure–based models can reconstruct CATH (Fig. 5); in so doing, we include a random baseline model as a sort of ‘negative control’ in gauging the performance of the DeepUrfold framework (Fig. 5, Supplementary Fig. 34, and Supplementary Tables 2 and 3). Among all these methods, our DeepUrfold approach yields results that are the most divergent from CATH, consistent with DeepUrfold’s approach of taking a wholly new view of the protein universe and the domain-level structural similarities that shape it. We also see that many other protein comparison algorithms, both sequence-based (Fig. 5, left) and structure-based (Fig. 5, right), have difficulty reconstructing CATH (possibly due to extensive manual curation of CATH), but more closely reproduce it than does our method (save Struct2Seq, which exhibits metrics similar to those of DeepUrfold). We suspect that this largely occurs because of DeepUrfold’s low-level feature encoding, which intentionally incorporates and integrates more *types* of information than purely 3D structural geometry.

**Fig. 5.**
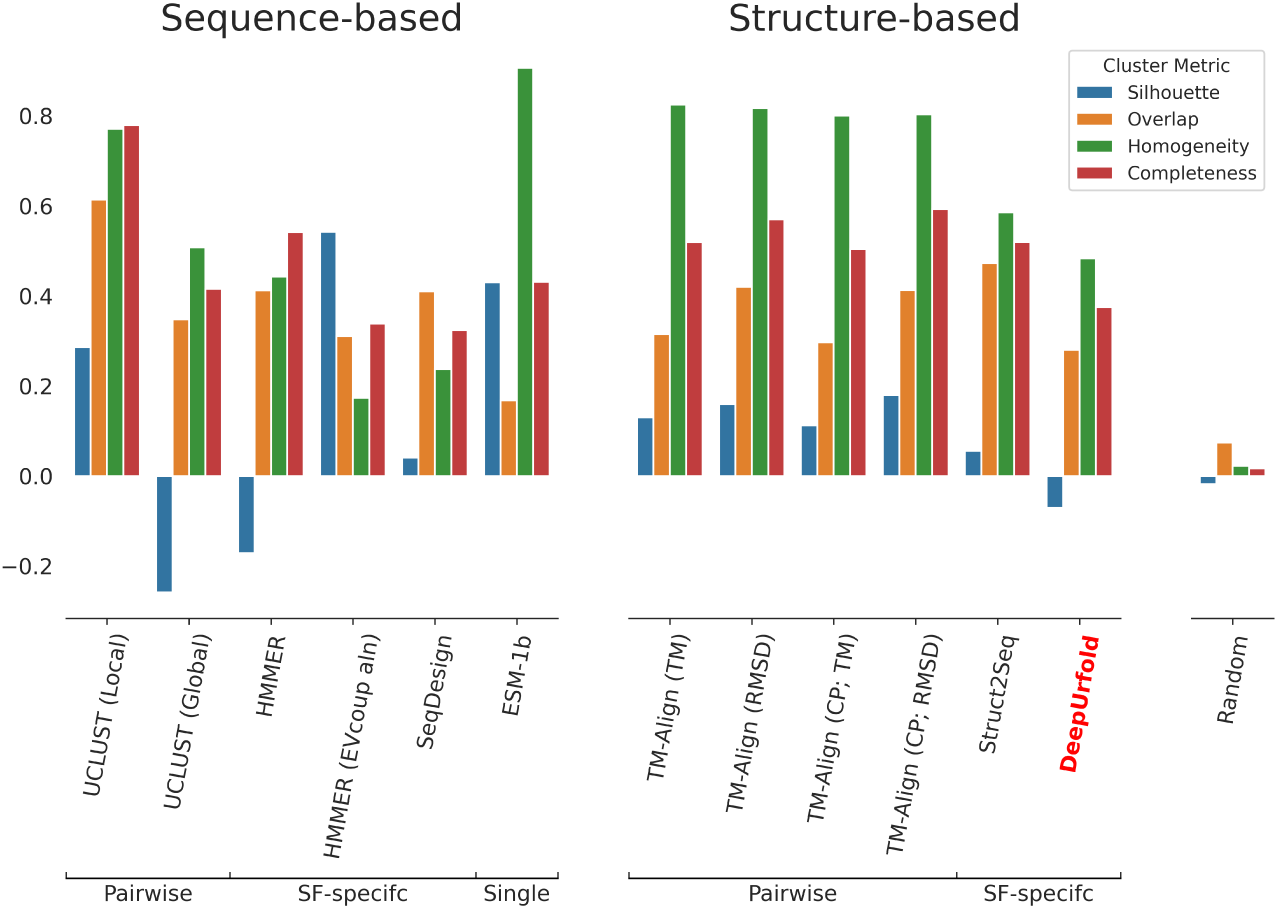
DeepUrfold and other comparative methods, relative to CATH. Here, we compare DeepUrfold to other sequence-based (left-half) and structure-based (right-half) protein similarity approaches by using each of them to attempt to reconstruct CATH’s organization of protein superfamilies. (For TM-Align, ‘CP’ stands for a circular permutation mode.) The scores from each of the algorithms, applied to the same protein dataset as used for DeepUrfold in this work, are used as edge weights to compute an SBM. In so doing, any score types that would increase with decreasing similarity (i.e., behave as a distance metric) were converted to a similarity metric by negation (*−x* or *−*log *x*). We take the communities at the lowest hierarchical level as clusters and employ various cluster comparison metrics to understand how well each algorithm/similarity metric can be used to recapitulate CATH. For each of these metrics (*silhouette*, *overlap*, *homogeneity* and *completeness*), a value of unity is deemed best. We also compared a uniform random grouping for 20 groups as a baseline/control experiment, as shown here in the right-most column and detailed in Supplementary Fig. 34 and Supplementary Table 3. Note that DeepUrfold does ‘poorly’ with these metrics because it does not produce the same clustering patterns—in other words, it is learning something quite different from these other algorithms, which more closely reproduce CATH.

## Discussion, Further Outlook

This work offers a new, structure-guided, community-based view of protein relationships, based on deep generative modelling. Using a deep learning-enabled framework that we term *DeepUrfold*, our approach aims to (i) explore and assess the Urfold model of protein structure relationships [11], in a rigorous/quantitative manner, and (ii) develop a platform for systematically identifying putative new urfolds. The following are key features of the DeepUrfold framework: (i) It is sensitive to 3D structure and structural similarity between pairs of proteins, but is minimally heuristic (e.g., it does not rely upon pre-set RMSD thresholds or the like) and, crucially, it is topology-agnostic and alignment-free (as it leverages latent space embeddings of featurized structures, versus direct 3D coordinates, for comparison purposes). (ii) Beyond the residue-level geometric information defining a 3D structure (i.e. coordinates), DeepUrfold is an extensible model insofar as it can incorporate any types of properties of interest, so long as such data can be encoded as part of the ‘featurization’ in a deep model—e.g. biophysical and physicochemical characteristics (electrostatic charge, solvent exposure, etc.), site-by-site phylogenetic conservation, and so on. (iii) The DeepUrfold method provides a quantitative metric, in the form of the deep neural network’s loss function (at the inference stage), that is amenable to approaches that are more generalized than brute-force hierarchical clustering; for instance, this work shows that we can use loss function scores in stochastic block modelling to construct mixed-membership communities of proteins. In the above ways, DeepUrfold can be viewed as an integrative approach that, while motivated by structural (dis)similarities across fold space, is also cognizant of *sequence*↭*structure*↭*function* interrelationships— even those which are quite weak. This is intentional: molecular evolution acts on the sequence/structure/function triad as its base ‘entity’, not on the purely geometric aspects of 3D structure alone. We suspect that any purely geometric/structure-based approach ultimately will be limited in its ability to accurately represent fold space (as also described in Supplementary Figs. 23-25 and Supplementary Discussion 5.3.2).

Using the DeepUrfold framework, we demonstrate (i) the general utility of a new type of similarity metric for representing and comparing protein domain structures, based on deep generative models and latent spaces, and (ii) that a mixed-membership community detection algorithm can identify what we previously suspected, via manual/visual analysis [11], to be putative urfolds. Finally, we emphasize that because DeepUrfold is agnostic of precise protein topology (i.e., order of connectivity of SSEs in 3D-space) it can readily detect levels of similarity ‘above’ the fold level (above CATH’s T’ level, below its ‘A’ level), including the potential of non-contiguous fragments. We believe that such spatially-compact groups of frequently recurring sub-domain fragments, sharing similar architectures (independent of topology) within a given group—which, again, we term an ‘urfold’—could correspond to primitive ‘design elements’ in the early evolution of protein domains [22]. We note that Kolodny [58] has made similar points.

Overall, the DeepUrfold framework provides a sensitive approach to detect and thus explore distant protein interrelationships, which we suspect correspond to weak phylogenetic signals (perhaps as echoes of remote/deep homology). Also notable, the embeddings produced by our VAE models and ELBO-based similarity scores provide new methods to visualize and interpret protein interrelationships on the scale of a full fold space. From these models, it is clear that there is a fair degree of continuity between proteins in fold space, and intermixing between what has previously been labeled as separate superfamilies; a corollary of this finding is that discretely clustering proteins, or their embeddings, is ill-advised [36] from this perspective of a densely-populated, smoother-than-expected fold space. An open question is the degree to which the extent of overlap between individual proteins (or groups of domains, as an urfold) in this fold space is reflective of underlying evolutionary processes, e.g. akin to Edwards & Deane’s finding [21] that “evolutionary information is encoded along these structural bridges [in fold space]”.

While the present work focused exclusively on developing DeepUrfold with CATH as a backdrop, it also would be intriguing to assess other classification schemes as contexts for DeepUrfold-based VAE models—specifically, SCOP, SCOP2 and ECOD. SCOP2 is particularly interesting because it aims to represent sub-domain-level similarities and evolutionarily-distant functional relationships by relaxing the strict constraints of hierarchical trees in favor of a graph-based approach to relationships [33]. A comparative analysis of DeepUrfold groupings (e.g., from an SBM) and SCOP2 groupings, in order to gauge any clear and readily identifiable points of concordance between these approaches, would be of great interest.

Another informative next step would be to use DeepUrfold to identify structural fragments that contain similar patterns of geometry and biophysical properties between proteins from quite different superfamilies. Notably, these fragments may be continuous or discontinuous, and pursuing this goal might help unify the ‘primordial peptides’ [2] and ‘themes’ [27] concepts with the Urfold hypothesis, allowing connections between unexplored (or at least under-explored) regions of fold space. Also, we suspect that ‘Explainable AI’ techniques, such as layer-wise relevance propagation (LRP; [59, 60]), can be used to elucidate which atoms/residues, along with their 3D locations and biophysical properties, are deemed most important in defining the various classification groups that are identified by DeepUrfold (i.e., the structural and physicochemical determinants of why a given protein falls into urfold versus urfold). This goal can be pursued within the DeepUrfold framework because we discretize full domain structures into voxels as part of the 3D-CNN data encoding scheme (Supplementary Fig. 1): thus, we can probe the neural network (i.e., trained model) to learn about specific voxels, or groups of specific voxels (e.g., amino acid residues), that contribute as sub-domain structural elements. Doing so would, in turn, be useful in finding common sub-domain segments from different superfamilies. We hypothesize that the most ‘relevant’ (in the sense of LRP) voxels would highlight important substructures; most promisingly, that we know the position, biochemical and biophysical properties, and so on about the residues would greatly illuminate the *physical* basis for the deep learning-based classification. In addition, this would enable us to explore in more detail the mechanistic/structural basis for the mixed-membership features of the SBM-based protein communities. Beyond helping to detect and define new urfolds, for use in areas like protein engineering or drug design, such communities of weakly-related proteins may offer a powerful new lens on remote protein homology.

## Methods

The following subsections describe the computational methodology that underlies the DeepUrfold framework.

### Datasets

Using ‘`Prop3D`’, a computational toolkit that we have developed for handling protein properties in machine learning and structural bioinformatics pipelines [47], we created a ‘`Prop3D-20sf`’ dataset. This dataset employs 20 highly-populated, diverse CATH superfamilies of interest (Fig. 1C); these SFs are tabulated in Supplementary Table 1, and annotated rationales for many of the SFs are delineated in Supplementary Discussion 5.1. Domain structures from each of the 20 SFs were ‘cleaned’ by adding missing residues with `MODELLER` (version 9.20) [61], missing atoms with `SCWRL4` [62], and protonating and energy minimizing (simple debump) with `PDB2PQR` (version 1.8) [63]. Next, we computed a host of derived properties for each domain in CATH4.3 [47], including (i) purely geometric/structural quantities, e.g. secondary structure labels 64] and solvent accessibility, (ii) physicochemical properties, e.g. hydrophobicity, partial charges, electrostatic potentials, (iii) basic chemical descriptors (atom and residue types), and (iv) phylogenetic conservation. As detailed in [47], these computations rely heavily on the `Toil` workflow engine [65], and data were stored using the `Hierarchical Data Format` (version 5) in a `Highly Scalable Data Service` (`HSDS`)–enabled environment. The domains from each SF were split such that all members of an `S35` 35% sequence identity cluster (pre-computed by CATH) were on the same side of the split; as described in [47], we constructed data splits so as to mitigate evolutionary data leakage’. We partition the protein data roughly as 80% training, 10% validation, and 10% test (https://doi.org/10.5281/zenodo.6873024; further technical details on creating and enacting `Prop3D` computational workflows can be found in [47]).

In our `Prop3D`-based dataset (`Prop3D-20sf`), each atom is attributed with the folowing seven groups of features, which are one-hot (Boolean) encoded: (i) Atom Type (`C`, `CA`, `N`, `O`, `OH`, `Unknown`); (ii) Residue Type (`ALA`, `CYS`, `ASP`, `GLU`, `PHE`, `GLY`, `HIS`, `LE`, `LYS`, `LEU`, `MET`, `ASN`, `PRO`, `GLN`, `ARG`, `SER`, `THR`, `VAL`, `TRP`, `TYR`, `Unknown`); (iii) Secondary Structure (`Helix`, `Sheet`, `Loop/Unknown`); (iv) Hydrophobic (or not); (v) Electronegative (or not); (vi) Positively-charged (or not); and (vii) Solvent-exposed (or not). For all of the DeepUrfold final production models reported here, the “residue type” feature was omitted because it was found to be uninformative, at least for this type of representation (see Supplementary Figs. 4-5, and more broadly Supplementary Figs. 2-7 for model training steps and classification metrics); interestingly, this finding about the dispensability of a residue-type feature was presaged in early work on this project (e.g., the receiver operating characteristic (ROC) curves in Fig. 2 of ref [66]).

### Protein 3D Structure Representation

We represent protein domains in DeepUrfold’s 3D-CNN by discretization into 3D volumetric pixels, or voxels, as described in Supplementary Fig. 1. Briefly, our method centers a protein domain in a 256^3^ Å^3^ cubic volume to allow for large domains, and each atom is mapped to a 1 Å^3^ voxel using a *k* D-tree data structure, with a query ball radius set to the van der Waals radius of the atom from a lookup table. If two atoms occupy the same given voxel—a possibility, as the solid diagonal of such a cube is 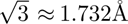 —then the maximum (element-wise) between their feature vectors is used for that voxel (justifiable because they are all binary-valued). Because a significant fraction of voxels in our representation domain do not contain any atoms, protein domain structures can be encoded in this way via a sparse representation; doing so, via an implementation using MinkowskiEngine [67], substantially reduces the computational costs of our deep learning workflow.

We sought to avoid learning (arbitrary) rotational information about domains in our 3D-CNN/VAE models, as there is no unique or ‘correct’ canonical orientation of a protein structure in R^3^. Therefore, we applied random rotations to each protein domain structure as part of the model training routine; these rotations were in the form of orthogonal transformation matrices randomly drawn from the Haar distribution, which is the uniform distribution on the 3D rotation group, i.e., SO(3) [68]. Tests of the rotational invariance of our various trained models can be seen in Supplementary Figs. 20-22.

### Stacked 3D-CNN/VAE Model Design and Training

A sparse 3D-CNN variational autoencoder was adapted from MinkowskiEngine [67, 69]. In DeepUrfold’s `Encoder`, there are seven blocks consisting of Convolution (n-*>*2n), BatchNorm, Exponential Linear Unit (ELU) activation functions, Convolution (2n-*>*2n), BatchNorm, and ELU, where n=[16, 32, 64, 128, 256, 512, 1024], or a doubling at each block. Finally, the tensors are pooled using a Global Pooling routine, and the model outputs both a normal distribution’s mean and log variance. Next, the learned distribution is sampled from (Supplementary Note 6) and used as input to the `Decoder`. The decoder also consists of seven blocks, where each block consists of ConvolutionTranspose(2n-*>*n), BatchNorm, ELU, Convolution(n-*>*n), BatchNorm, and ELU. Finally, one more convolution is used to output a reconstructed domain structure in a 256^3^ Å^3^ volume. A detailed layout of DeepUrfold’s model architecture can be found in Supplementary Methods 4.1.

In VAE training calculations, a well-established ‘reparameterization trick’ enables gradients to be computed for backpropagation steps despite the VAE’s latent space variables being sampled stochastically. This is achieved by making only the mean (*µ*) and variance (*σ*) differentiable, with a random variable that is normally distributed (𝒩 (0, **I**)). That is, the latent variable posterior **z** is given by **z** = ***µ*** + ***σ*** ⊙𝒩 (0, **I**), where ⊙ denotes the Hadamard (element-wise) matrix product and 𝒩 is an ‘auxiliary noise’ term [42].

In DeepUrfold, we optimize against the negative of the evidence lower bound, –(ELBO). As described in Eq 1, Supplementary Discussion 5.2 and Supplementary Fig. 8, this approach combines into a single quantity (i) the mean squared error (MSE) of the reconstructed domain and (ii) the difference between the learned distribution and the true distribution of the SF (i.e., the KL divergence, or relative entropy between the true/underlying distribution of the data given a model, *p*, and our learned/inferred posterior distribution of latent parameters given the data, *q* [42]).

We used stochastic gradient descent (SGD) as the optimization algorithm for parameter updates during NN model training, with a momentum of 0.9 and 0.0001 weight decay. We began with a learning rate of 0.2 and decreased its value by 0.9 every epoch using an exponential learning rate scheduler. Our final network has ≈110M parameters in total and all of the SF-level networks were trained for 30 epochs, using a batch size of 255 (Supplementary Fig. 2 provides an illustrative example of model training with Igs). We utilized the open-source frameworks `PyTorch` [70] and `PytorchLightning` [71] to simplify training and inference, and to make the models more reproducible.

To optimize/tune hyperparameters for DeepUrfold’s VAE, we used `Weights & Biases` Sweeps [72] to parameter-scan across the batch size, learning rate, convolution kernel size, transpose convolution kernel size, and convolution stride in the Ig model, while minimizing the –(ELBO). We used a Bayesian Optimization search strategy and ‘hyperband’ method with three iterations for early termination. We found no significant changes to parameters, and therefore used the following default values: convolution kernel size of 3, transpose convolution kernel size of 2, and convolution stride of 2.

Because of a large-scale class imbalance between the number of domains in each superfamily (e.g., over-representation of Igs), we follow the ‘one-class classifier’ approach, creating one VAE for each SF. As part of our ‘control experiments’, we also trained a joint SH3 OB model and compared random over- and under-sampling from ImbalancedLearn [45] on joint models of these two SFs (Supplementary Fig. 10). Finally, a ‘Global Model’ was also trained by combining all domains from all 20 SFs and using ImbalanceLearn under-sampling, with training for 100 epochs (Supplementary Fig. 3).

The early stages of DeepUrfold’s workflow, involving structure sanitization and featurization, creation of training data splits, etc., made extensive use of `Prop3D` and ts attendant `Prop3D-20sf` dataset (supplying 20 highly-populated CATH SFs); details on `Prop3D` pipelines and workflow performance can be found in [47]. All of the NN models developed and used throughout this work, including (i) the 20 individual SF- level models, (ii) the Global Model and (iii) joint SH3 OB models, were trained on a Lambda Labs Deep Learning platform, using between one and four NVIDIA RTX A6000 GPUs.

### Evaluation of Model Performance

We calculated the area under the ROC curve (AUROC) and the area under the precision-recall curve (AUPRC) for the 20 SFs. Representative domains, as defined by CATH, for each superfamily were subjected to their SF-specific VAE models and predicted values were micro-averaged to perform AUROC and AUPRC calculations. Immunoglobulins were chosen for purposes of display in this work (Supplementary Figs. 2, 4-7), and the results for all SFs can be found in the extended Supplementary Information. All SFs resulted in roughly similar metrics for each of the seven different groups of encoded features (Supplementary Figs. 4-5).

### Assessing Topological Sensitivity by Scrambling Loops

To gauge the sensitivity of our trained DeepUrfold models to loop orderings (i.e., topology), we subjected artificial protein structures, with systematically permuted secondary structural elements, to superfamily-specific VAEs. To do this, we generated a series of fictitious folds by implementing a multi-loop permutation algorithm [46], allowing us to systematically ‘re-wire’ the SSEs found in representative SH3 and OB domains in order to extensively sample possible topological orderings. Having stitched together the SSEs in various orders, we relaxed the conformations/energetics of each new 3D structure using the `MODELLER` suite (version 9.20) [61]. While 96 unique permutations are theoretically possible for the 4-stranded β-sheet core of the SBB, and 960 orderings for all 5 strands (*n*!·2*^n−^*^2^ unique topologies are possible for a sheet of *n* strands [9]), note that fewer than these numbers of permuted domains were able to be successfully modelled; presumably this is because only some geometries lie within the radius of convergence of `MODELLER` (e.g., the loop-creation algorithm did not have to span excessive distances in those cases, try to resolve knots, and so on).

Next, each novel permuted structure was subjected to an inference pass through a VAE model that was trained on all other (wild-type) domains from the SH3 homologous superfamily. Fits to the model were approximated by the log-likelihood score of the permuted structure, represented by –(ELBO) scores, which can be viewed as a similarity metric (goodness-of-fit of a given structure to the VAE model). We also calculated a ‘background’ distribution of each model by performing an all-vs-all `TM-align` calculation for all domains in our representative CATH domain set; in this step, we recorded any domains that have a TM-score ≤ 0.3, as that threshold quantity is thought to correspond to domains that have random 3D structural similarity (see also the description in Supplementary Fig. 9).

### DeepUrfold’s Ability to Discern Structural Fragments

To further assess and probe the behavior of DeepUrfold models, we examined if its trained VAEs are better at detecting similarities between shared common fragments, rather than global/domain-level structural similarity; note that such might be the case with potential urfolds, which we posit to share extensive sub-domain similarity. For the SBB urfold, consisting of SH3 and OB folds, we extracted tertiary structural motifs (TERMs [49, 50]) from representative domains of both the SH3 and OB superfamilies; this was achieved using the code `TERMANAL` [49] (version as of 2015), while preserving all intermediate fragment files [49]. Next, each set of TERM fragments (OB-derived and SH3-derived) was subjected to separate inference passes through both the SH3 and OB VAE models, and fits to each model were approximated by the log-likelihood scores of the TERM fragments, represented by –(ELBO) score distributions (Fig. 2B, D).

### Exploration of Latent-space Organization

We subjected each representative domain (numbering 3,674), drawn from all the individual SFs, to an inference pass through each of the 20 SF-specific DeepUrfold models, and visualized the 20 different latent space embeddings for all representatives from each separate model. These results are further detailed in Supplementary Discussion 5.3 and Supplementary Figs. 11-25: in particular, Supplementary Discussion 5.3.1 describes the individual, SF-level feature embeddings that we analyzed as 20 independent UMAP-learned manifolds, each of which are shown as individual panels in Supplementary Fig. 18. More concretely, the ‘latent space embedding’ for a given domain subjected to one SF-specific VAE corresponds to a 1,024-dimensional vector that describes the representative domain in its most ‘compressed’ or ‘distilled’ form in the feature space learned by the VAE model—as an amalgamation of the position of each atom, their biophysical properties (represented by the mean of the learned distribution), and any other features that were included in the model (e.g., phylogenetic properties; see above). As an example, the latent space organization of the SH3 VAE model is shown in Fig. 3A (as a UMAP reduction).

We computed individual UMAP reductions for each SF-level VAE latent space, as shown in Fig. 3A (for the SH3) and Supplementary Fig. 18’s panels (for all 20 SFs). In addition, we sought to visualize an *overall* DeepUrfold latent space, spanning the 20 SFs. However, DeepUrfold-learned embeddings from two different, independently-trained SF-specific VAE models are not necessarily directly comparable, as they can occupy disparate regions of the learned latent (hyper)space. This, in turn, makes it difficult to combine or ‘aggregate’ all 20 latent spaces, from the 20 SF VAE models, nto a single representation. We addressed this challenge by devising three alternative approaches to obtain an overall representation of DeepUrfold’s latent spaces: (i) creation of a ‘Concatenated Model’ (Fig. 3B), wherein we pooled the latent spaces for every domain (from each SF-specific VAE) into a single dataset by concatenating on the feature or column dimension, e.g. the shape of the dataset from a single SF model s (3674, 1024) and the combined dataset then becomes (3674, 20480) after concatenation; (ii) training of a single, monolithic ‘Global Model’ (Fig. 3C), using domains from all SFs and compensating for SF class imbalance; and (iii) ‘shifting’ the embedding vectors from each individual SF VAE model to a common region of latent space, via a method known as Optimal Transport (OT) for domain adaptation. As shown in Supplementary Fig. 19 and detailed in its accompanying caption, we applied the OT algorithm (using Sinkhorn-based transport with group LASSO L1L2 regularization) and then concatenated on the feature dimension; reassuringly, this process achieved similar results as our more näıve concatenation approach, inasmuch as SFs exhibited a clear dispersal in terms of SSE content (i.e., the non–OT-based approach [Fig. 3B] and OT-based approach [Supplementary Fig. 19] are roughly similar in this regard). Thus, in total we applied four strategies to handling VAE embeddings: (i) single– SF-level models (Fig. 3A, Supplementary Fig. 18), (ii) a Concatenated Model (Fig. 3B), (iii) a Global Model (Fig. 3C), and (iv) an OT-aligned & Concatenated Model (Supplementary Fig. 19).

Finally, in order to aid visualization of the learned embeddings, for each of the above four approaches we reduced the number of latent space dimensions to two, giving a (3674, 2)-sized matrix across all domains. We achieved this via three different dimensionality-reduction approaches—UMAP, t-SNE and PCA. As a manifold earning approach, the UMAP algorithm is more generally versatile than principal component analysis (PCA), the latter of which is a subspace projection method that assumes linearity in the data; nevertheless, for completeness we also applied PCA to the DeepUrfold embeddings for the Concatenated Model (Supplementary Fig. 13) and the Global Model (Supplementary Fig. 17). Also, UMAP more robustly captures ong-range structure/correlations in a dataset than does the common t-distributed stochastic neighbor embedding (t-SNE) approach, which we also applied to the DeepUrfold embeddings for the Concatenated Model (Supplementary Fig. 12) and the Global Model (Supplementary Fig. 16). Given our näıvety about DeepUrfold’s latent spaces, we utilized UMAP as a de facto reduction approach because it provides both (i) a well-formed metric notion of local distances (e.g., within-clusters) and (ii) better preserval (versus t-SNE) of the topological structure/relationships amongst more distant points in a dataset, e.g., more global-scale, between-cluster orderings.

### Mixed-membership Community Detection via SBMs

We performed all-vs-all comparisons of domains and superfamilies by subjecting representative protein domain structures from each of the 20 chosen SFs through each SF-specific one-class VAE model. The –(ELBO) loss score for each (*i, j*) pair 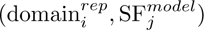 can be used to quantitatively evaluate pairwise ‘distances’ between SFs by treating the complete set of distances as a fully connected bipartite graph between domains *i* (one side of the graph) and SF models *j* (other side of the graph), defined by adjacency matrix ***A****_ij_*, with edges weighted by the −log(−(ELBO)) scores from the covariate matrix, ***x*** . Stochastic Block Models (SBM; [53]) offer a generative, nonparametric Bayesian inference–based approach for community detection in random graphs [54]. Therefore, we used SBM algorithms to partition the DeepUrfold-derived bipartite graph into communities of domains that have similar distributions of edge covariates between them. Using the SBM likelihood equation (Eq 2), inference is done via the posterior:

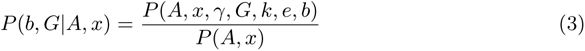

where ***b*** is the overlapping partition, ***e*** is the matrix of edge counts between groups, ***k*** is the labelled degree sequence, and ***G*** is a tensor representing half-edges (each edge end-point *r, s*) to account for mixed-membership, satisfying 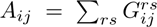. Edge covariates ***x*** are sampled from a microcanonical distribution, *P* (*x|G, γ*), where *γ* adds a hard constraint such that 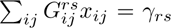 (personal communication with T. Peixoto and Sec. VIIC in [52]).

Using the same SBM approach as we did for post-processing the DeepUrfold-derived data (i.e., ELBO-quantified fits between domain representatives and SF-specific VAE models), we also compared our results to community analyses of data that we obtained by utilizing state-of-the-art sequence– and structure–based methods for comparing proteins (e.g., `HMMER`, `ESM-1b`, `SeqDesign`, etc. listed in Fig. 5 and Supplementary Table 3). All SBMs were created using fully-connected *n×m* bipartite graphs, linking *n* CATH S35 domains to *m* SF models. In our current work, we used 3,674 representative CATH domains from 20 SFs, yielding a 3,674 × 20-element similarity matrix for each of the various methods (`UCLUST`, `HMMER`, `SeqDesign`, etc.) that we sought to compare. Each SBM was degree-corrected, overlapping, and nested and fit to a real normal distribution of edge covariates. For those methods that give decreasing scores with increasing similarity (i.e., closer to zero is greater similarity), we log–transformed each score, whereas values from methods with a non-inverse relationship between the score metric and inferred similarity (i.e., higher values mean greater similarity) were unaltered.

While only ‘superfamily-specific’ methodologies/models would be directly comparable to the task performed by DeepUrfold (e.g., where *n×m* matrices are the original output created by subjecting *n* CATH representative domains without labels to *m* SF-specific models), for purposes of comparison in this work we have also included ‘pairwise’ and ‘single model’ methods (Fig. 5). This was accomplished in the following way: For pairwise approaches, an all-vs-all *n×n* similarity matrix was created and then converted to *n×m* by taking the median distance of a single CATH domain to every other domain in a given SF. What we are calling ‘single model’ approaches here are those wherein a single model is trained on all known proteins and outputs a single embedding score for each domain, creating an *n ×* 1 vector. To convert that data form into an *n × m* matrix, we took the median distance of a single CATH domain embedding to every other domain embedding from a given SF.

### Evaluating SBM Communities, and Comparing to CATH

Because we have no ground truth for the new Urfold view of protein structure simlarities (and the resultant protein universe), we applied cluster comparison metrics to evaluate each SBM community, both in a self-contained manner and as referred against the original CATH clusterings. The specific measures we considered include the *silhouette score*, *partition overlap*, *homogeneity* and *completeness*, for each of the various protein comparison approaches listed in Fig. 5 (and Supplementary Table 3):

- **Silhouette Score:** Provides a measure of how similar an object is to its own cluster (*cohesion*) compared to next-closest clusters (*separation*), with values ranging from −1 (poor grouping) to +1 (ideal).
- **Overlap:** Describes the maximum overlap between partitions, by solving an instance of the maximum weighted bipartite matching problem [56].
- **Homogeneity:** The optimal value (1) occurs when each cluster contains only members of a single class; this metric ranges from [0, 1].
- **Completeness:** Ranging from [0, 1], the optimal value (1) occurs when all members of a given class are (presumably correctly) assigned to the same (completed) cluster.

All of our comparisons start by using the sequence and structure representatives from CATH’s S35 cluster for each of the 20 superfamilies of interest. The code `USE- ARCH` (version 11.0.667) [73] was run twice with parameters -allpairs local and -allpairs global; both runs included the -acceptall parameter. `HMMER` (version 3.3.2) [74] models were built using (i) `MUSCLE` (version 3.8.31) [75] alignments from CATH’s S35 cluster, and (ii) a deep MSA created from `EVcouplings` (version v0.1.1) 76] using `jackhmmer` (via `HMMER` 3.3.2) [74] and `UniRef90` of the first S35 representative for each superfamily. Each `HMMER` model was used to search all representatives, reporting all sequences with bitscores 10^12^. `SeqDesign` (commit eb6308f) [77] was run using the same MSAs from `EVcouplings`. Finally, we also compared our DeepUrfold results against the `ESM-1b` pre-trained protein language model [78].

For other structure-based comparisons, we ran `TM-Align` (C++ version 2017.07.08) [79] on all representative domains, with and without allowing for circular permutations (‘CP’ in Fig. 5), and while saving the RMSD and TM-score values. `Struct2Seq` (commit 22e497a) [80] was executed with default parameters after first converting domain structure representatives into dictionaries in order to match the required form of input.

Finally, as a baseline, we also compare random groupings to CATH. To do so, we first created a uniform random grouping with 20 groups using numpy’s random.choice function. Next, we tried using the same SBM clustering as above, using random weights with numpy’s random.rand function. The random SBM eventually converged to a solution with only two groups: one for all domains and another for all VAE models (see Supplementary Fig. 34 and the details in its caption).

## Data Availability

The `Prop3D` framework, used to create, share and load datasets, as well as its associated `Prop3D-20sf` pre-built dataset, are available at https://prop3d.readthedocs.io/. `Prop3D` contains instructions for connecting to the public `Prop3D-20sf` HSDS endpoint (http://prop3d-hsds.pods.uvarc.io/about) and parsing the `Prop3D-20sf` raw hdf5 file (https://doi.org/10.5281/zenodo.6873024). The extended Supplementary material, including the 20 pre-trained SF models and raw output from the stochastic block modelling of DeepUrfold (as well as other tools used for comparative analysis) can be found at https://doi.org/10.5281/zenodo.13228646. We also provide an accompanying website to explore the SBM communities and the CATH hierarchy at https://bournelab.org/research/DeepUrfold/.

## Code Availability

All code used to build datasets and train models can be found at http://github.com/b ouralab/Prop3D [81] and http://github.com/bouralab/DeepUrfold [82], respectively.

## Acknowledgements

Portions of this work were supported by a University of Virginia Presidential Fellowship in Data Science (EJD) and NSF Career award MCB-1350957 (CM). We thank Luis Felipe Murillo and John Readey for support with HSDS, as well as Jaime Iranzo and Tiago Peixoto for discussions regarding SBMs; we also thank Menuka Jaiswal, Saad Saleem, and Yonghyeon Kweon for early efforts on this project. Finally, we thank the anonymous reviewers for insightful questions and substantive feedback, ultimately enabling this work to be much improved.

## Author Contributions

EJD designed and implemented DeepUrfold. CM and SV developed the initial Urfold model. EJD and CM led the manuscript preparation, and CM and PEB advised the project.

