## Supplementary Information for "Deep Generative Models of Protein Structure Uncover Distant Relationships Across a Continuous Fold Space"

**6 Supplementary References** **47**

### 1 Supplementary Figures

#### 1.1 Voxelizing & Featurizing Protein Structures

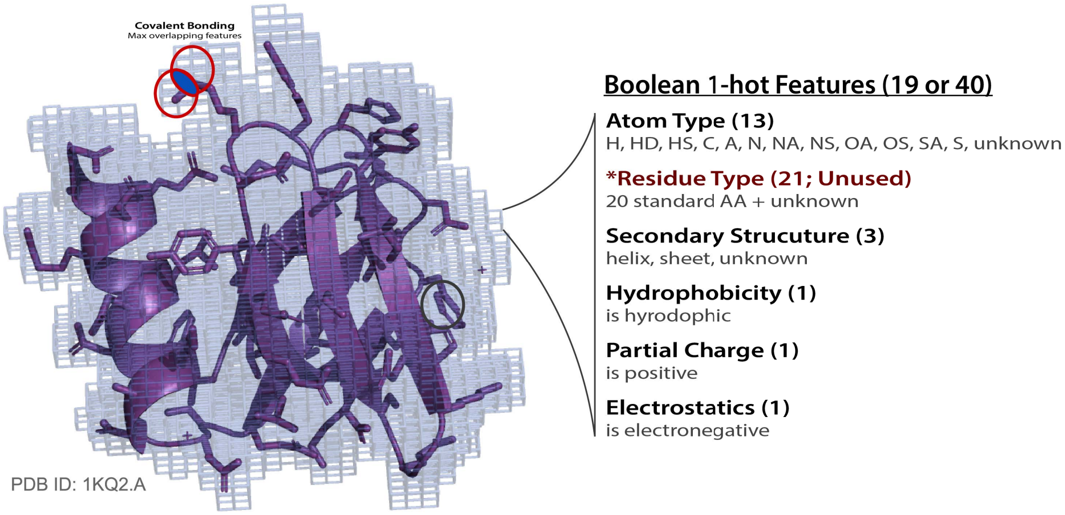

**Supplementary Figure 1:** In DeepUrfold, a protein domain structure is voxelized by (i) centering it in a $256^3$ Å$^3$ volumetric mesh, and (ii) discretizing each atom to fit a 1 Å$^3$ voxel, the assignment being based on searching a $k$D-tree with a radius equal to the current atom's van der Waals radius (and the $k$D-tree initialized by the full 256 Å$^3$ volume, with 1 Å$^3$ resolution). If two or more atoms occupy a single voxel, the maximum value over each feature is utilized, thereby accounting for covalent bonding. Each voxel contains a 1-hot feature vector with 19 or 40 elements, the actual number depending on if residue_type is included. Note that a residue_type feature was not included in the models shown in this work because (and interestingly) doing so was found to yield poorer reconstruction metrics, suggesting this as a dispensable feature for this type of model (see Supplementary Figures 4 and 5). Finally, note that many further details and considerations in the featurization process that we used for DeepUrfold can be found in our recent Prop3D article [1].

#### 1.2 Model Development & Metrics

##### 1.2.1 Training & Validation Loss

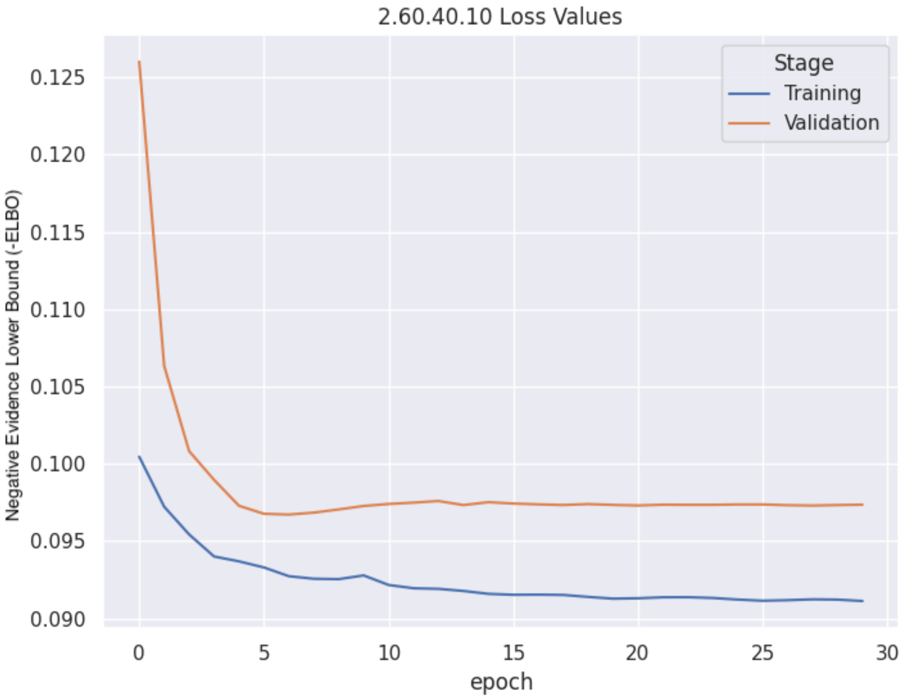

**Supplementary Figure 2:** As an example of the DeepUrfold training workflow at the single-SF level, the `2.60.40.10` Ig model was trained for 30 epochs using a 80%/10% split from CATH's S35 clusters (test set [10%] not shown).

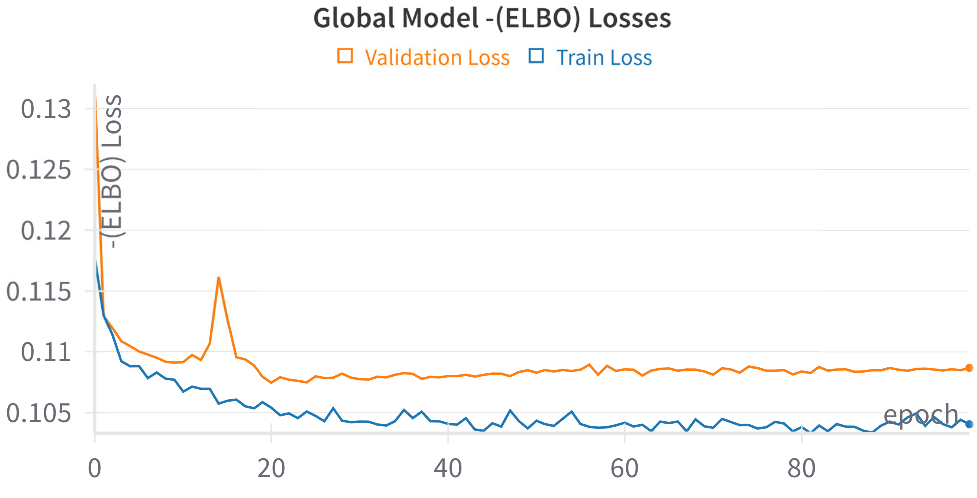

**Supplementary Figure 3:** We also trained a single, monolithic 'Global Model', wherein all 20 SFs were used as input, with training over 100 epochs. Each SF was split using a 80%/10% split from their CATH's S35 clusters (test set [10%] not shown), and class imbalance was accounted for via *ImbalancedLearn* down-sampling (see Methods section of the main text).

##### 1.2.2 Classification Metrics (with Residue Type)

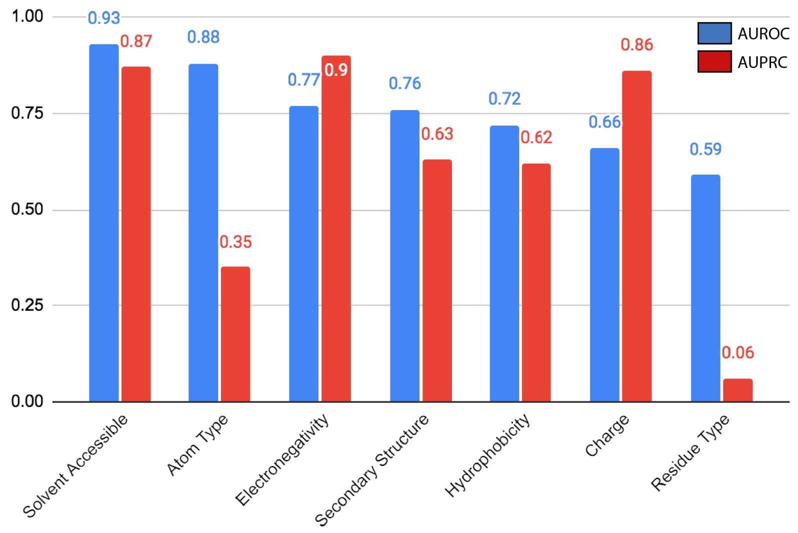

**Supplementary Figure 4:** We trained an Ig-specific model 7 different times for 7 different groups of features, excluding all of the other feature groups, yielding the results shown here. We compared the reconstructed values for each set of feature-values to the input using Receiver Operating Characteristic (ROC) and Precision-Recall Curves (PRC)[1], saving the AUC (Area Under the Curve) for each. Most features could be reconstructed well (AUC $\geq 0.6$) except for residue_type, so we removed that one from further models. We suspect that residue_type is not as decisive a feature in the all-atom models such as are being trained in the DeepUrfold work presented here, perhaps because it is too coarse-grained to be meaningful in this context.

##### 1.2.3 Classification Metrics (without Residue Type)

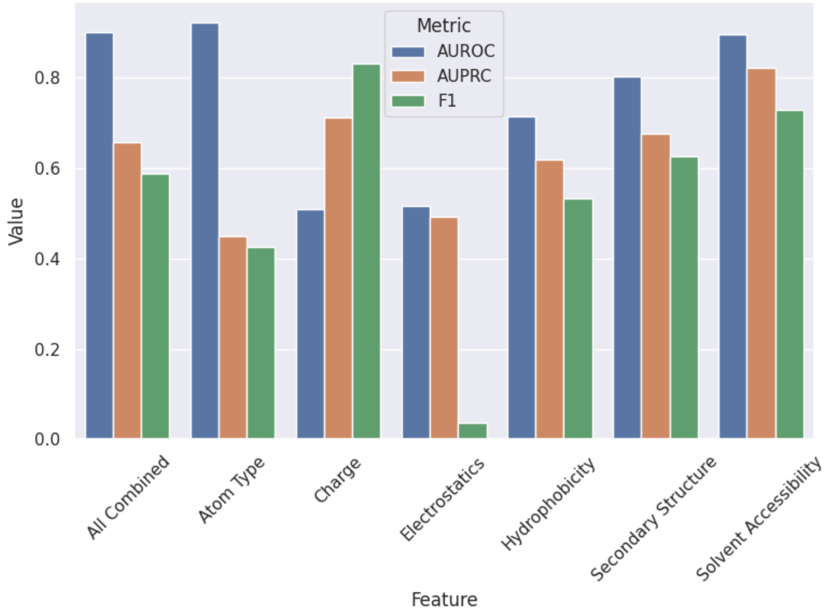

**Supplementary Figure 5:** Here, the `2.60.40.10` model was trained with all features, but individual feature groups were separated in order to perform micro-averaging of ROC, PRC, and F1 scores.

[1]The ROC curve plots the *true positive rate*, $\frac{TP}{TP+FN}$ (also known as *sensitivity* or *recall*), against the *false positive rate*, $\frac{FP}{FP+TN}$, for a series of discrimination thresholds; the PRC plots the *precision*, $\frac{TP}{TP+FP}$, versus the recall.

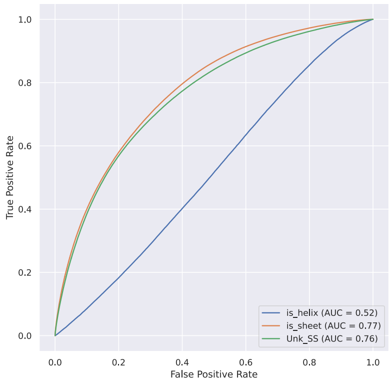

**(a)** Micro-averaged ROC Curve for separated features in the Secondary Structure feature group.

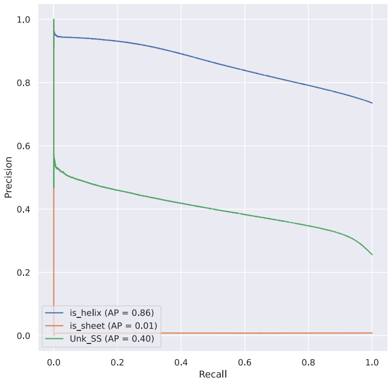

**(b)** Micro-averaged PRC Curve for separated features in the Secondary Structure feature group.

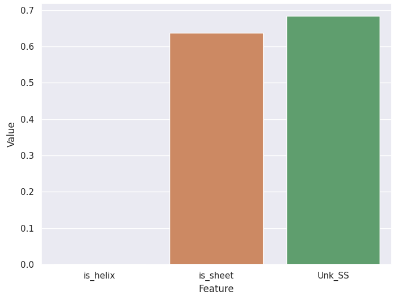

**(c)** Micro-averaged F1 values for separated features in the Secondary Structure feature group.

**Supplementary Figure 6:** Classification metrics for separated features in the Secondary Structure feature group.

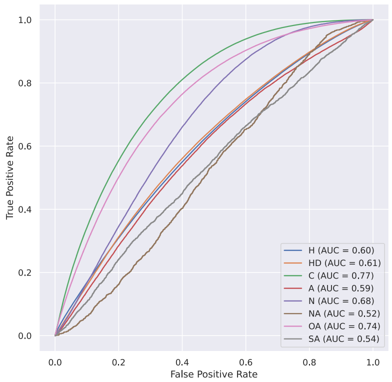

**(a)** Micro-averaged ROC Curve for separated features in the Atom Type feature group.

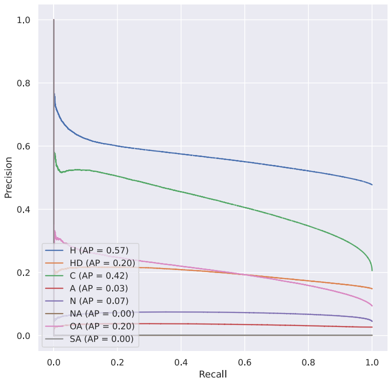

**(b)** Micro-averaged PRC Curve for separated features in the Atom Type feature group.

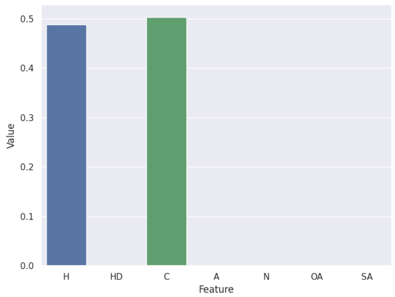

**(c)** Micro-averaged F1 values for separated features in the Atom Type feature group.

**Supplementary Figure 7:** Classification metrics for separated features in the Atom Type feature group.

#### 1.3 Interpreting DeepUrfold's ELBO-based Scores & Histograms

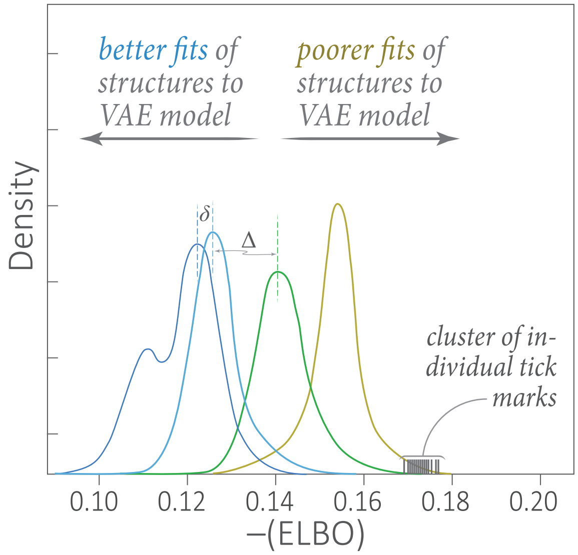

**Supplementary Figure 8:** This schematic aids the interpretation of results such as shown in Fig 2 of the text. Briefly, (i) the curves correspond to distributions (i.e., multiple inference calculations for a set of many structures), (ii) individual tick marks, lying along the abscissa, arise from a single pass (i.e., forward inference pass) through a trained DeepUrfold VAE model, and (iii) a rightward shift of a distribution relative to another, e.g., blue $\rightarrow$ green $\rightarrow$ yellow here, indicates progressively poorer fits to a given DeepUrfold VAE model. A tight cluster of ticks, such as shown here, might correspond to a series of systematically-rotated 3D structures. Note that differences between distributions can be significant and meaningful with respect to the learned models, even though such shifts may be relatively small (denoted by a '$\delta$') or large ($\Delta$).

#### 1.4 Multi-loop Permutation Experiments

##### 1.4.1 Sample Permutants

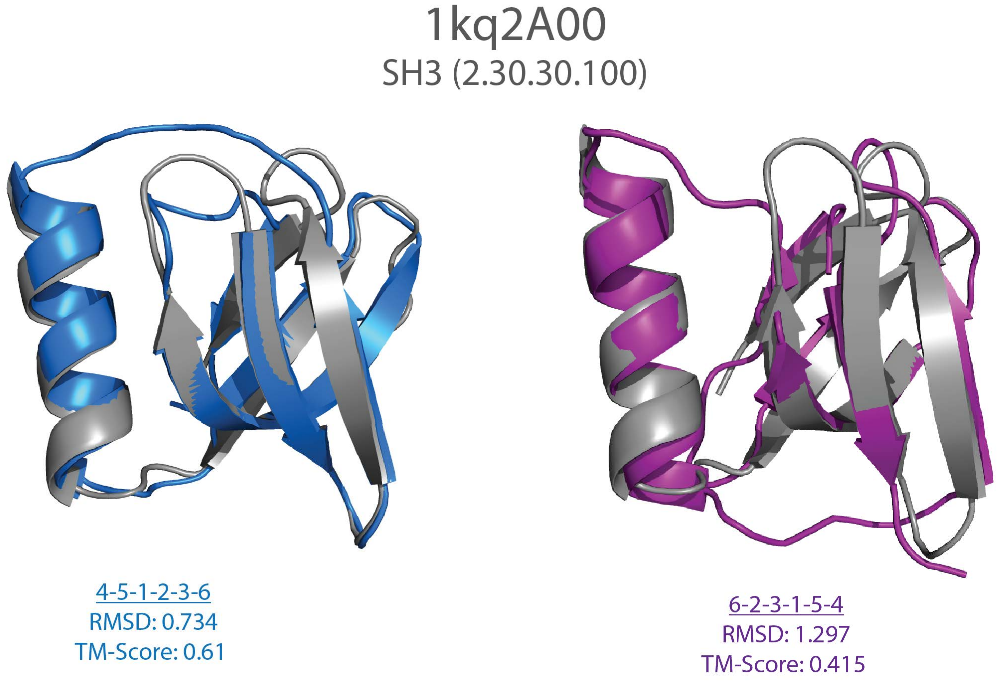

**(a)** Two exemplary SH3 permutants (blue and purple ribbon diagrams) are shown, overlaid on a wild-type SH3 scaffold in grey (PDB 1KQ2, from CATH `2.30.30.100`). Note that these two orderings of SSEs (4-5-1-2-3-6 [left] and 6-2-3-1-5-4 [right]), which are 'scrambled' with respect to the wild-type, are not related by a simple (circular) permutation.

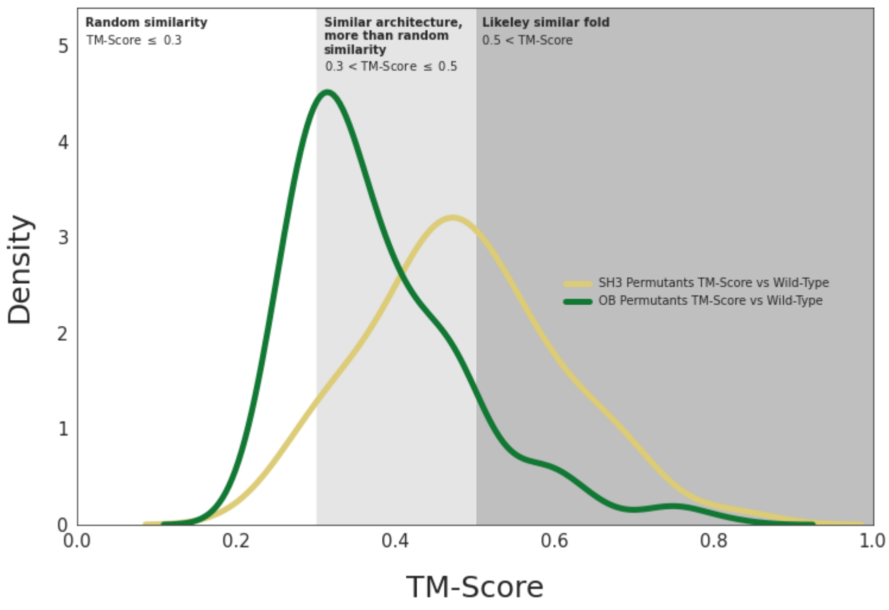

**(b)** TM-scores for multi-loop permuted structures. Many of the SH3 (yellow curve) and OB (green) permutant TM-scores fall between ≈0.3-0.5, indicating that the structures are more than randomly similar, though not identical folds.

**Supplementary Figure 9:** Shown here, with a focus on the SH3 domain, is an example of the systematic multi-loop permutation calculations that were performed in testing and developing DeepUrfold.

##### 1.4.2 Class Imbalance Scores

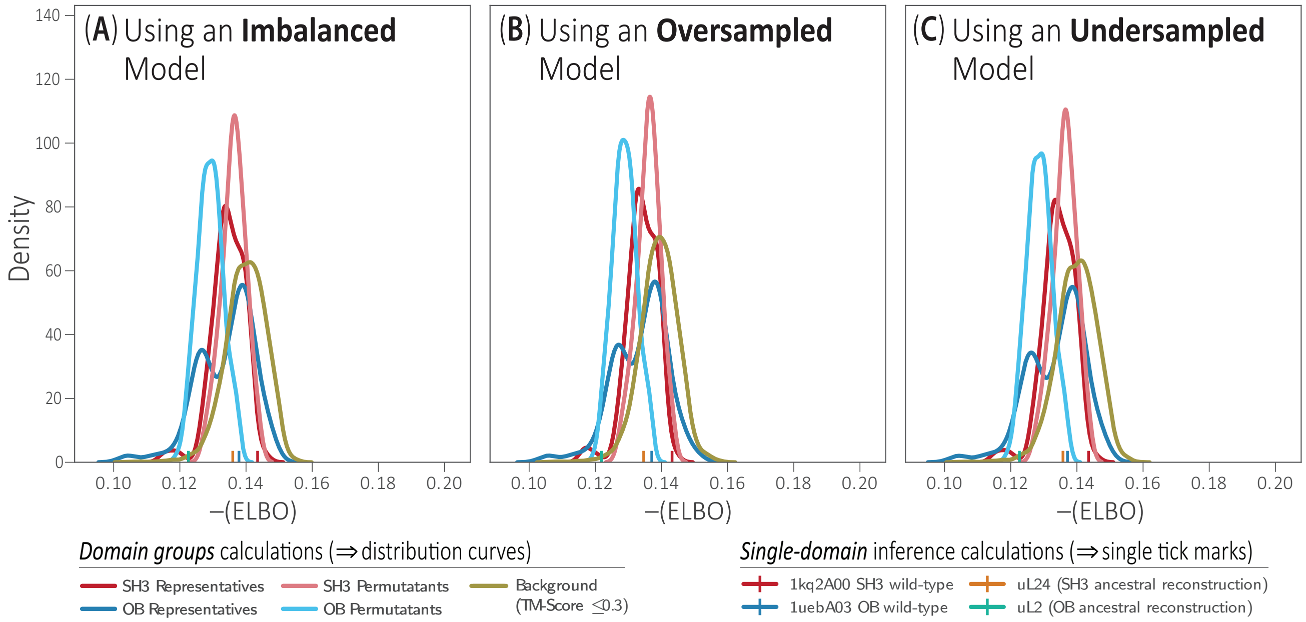

**Supplementary Figure 10:** In order to assess how class imbalance might affect our DeepUrfold models, we trained three joint SH3 and OB models: (A) using all domains from each superfamily; (B) over-sampling SH3 domains to match the number of OB domains; and (C) under-sampling OB domains to match the number of SH3 domains. We found no significant change between the results, in terms of –ELBO score distributions, when performing calculations with representative structures (cyan and pink traces), multi-loop permuted models (yellow and green traces), or ancestral versions of the SH3 and OB (uL2, uL24), suggesting that class imbalance is not a significant concern, at least for these SFs. Note that in each case the background distributions (gold traces) lie furthest towards the right (less negative –ELBO values), as would be expected if the DeepUrfold VAE models are learning real, SF-specific signals.

In terms of the potential significance of these results, we can elaborate a bit more on the single-tick entities shown here along the $x$-axis (similarly to the tick marks in Fig 2 of the main text, these values are from individual inference passes through a trained VAE model): Those specific single markings refer to DeepUrfold's –(ELBO) values computed for a possible 'ancestral' domain of the extant SH3 and OB superfolds, as examined in Alvarez-Carreño, Williams & colleagues' recent work "*Creative destruction: New protein folds from old*" [2]. For example, the datum labeled 'uL2' is a *universal large ribosomal subunit protein*, which is "*arguably one of the oldest ribosomal proteins*" and which can be interconverted between the SH3 and OB folds via simple circular permutation. Williams & colleagues have speculated as to which could have been the true precursor (SH3? OB?, etc.?). DeepUrfold offers a quantitative metric/tool by which we can attempt to distinguish between SH3-first vs OB-first possibilities: With our DeepUrfold models, we find that Williams' SH3 ancestral/precursor 3D structure indeed scores best against the SH3 VAE model, and likewise their ancestrally-reconstructed OB scores best against our OB VAE model (not shown). Intriguingly, for the joint SH3 ∪ OB models shown here we find the OB progenitor to score best (uL2 [cyan tick] lies most leftward, towards better fits to the DeepUrfold model), suggesting that the OB may be the historically most antecedent domain. Were that the case, then perhaps the progenitor evolved—e.g., via Alvarez-Carreño et al.'s "creative destruction" mechanism—into the contemporary OB and SH3 domains?

### 1.5 Latent Space Visualization and Exploration

#### 1.5.1 Concatenated Model

##### 1.5.1.1 UMAP

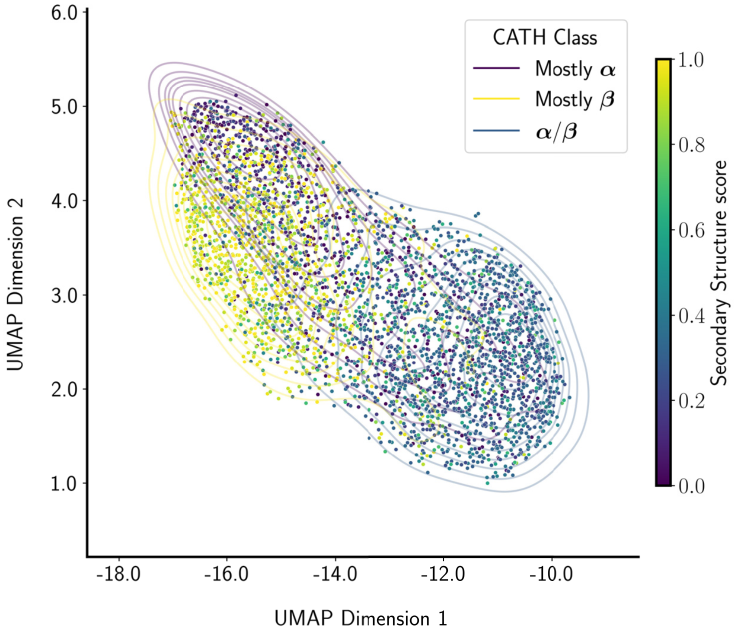

**(a)** Colored by secondary structure content

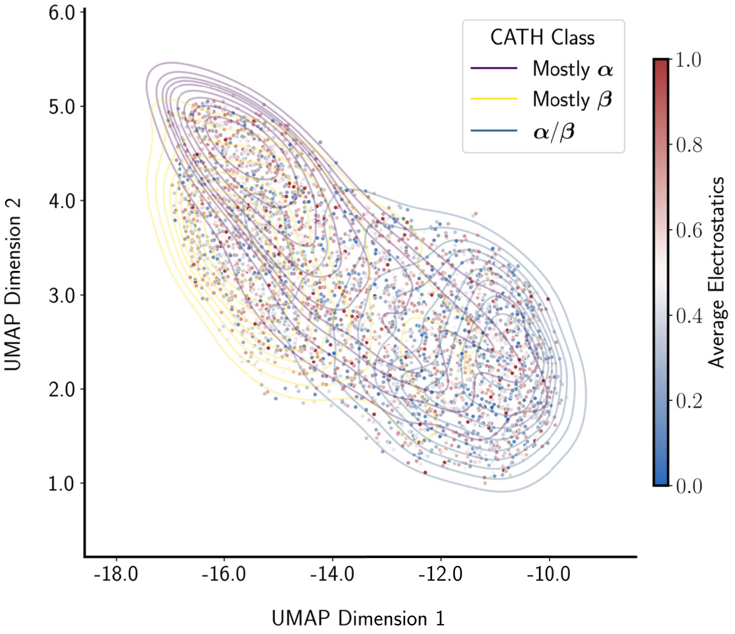

**(b)** Colored by averaged electronegativity values

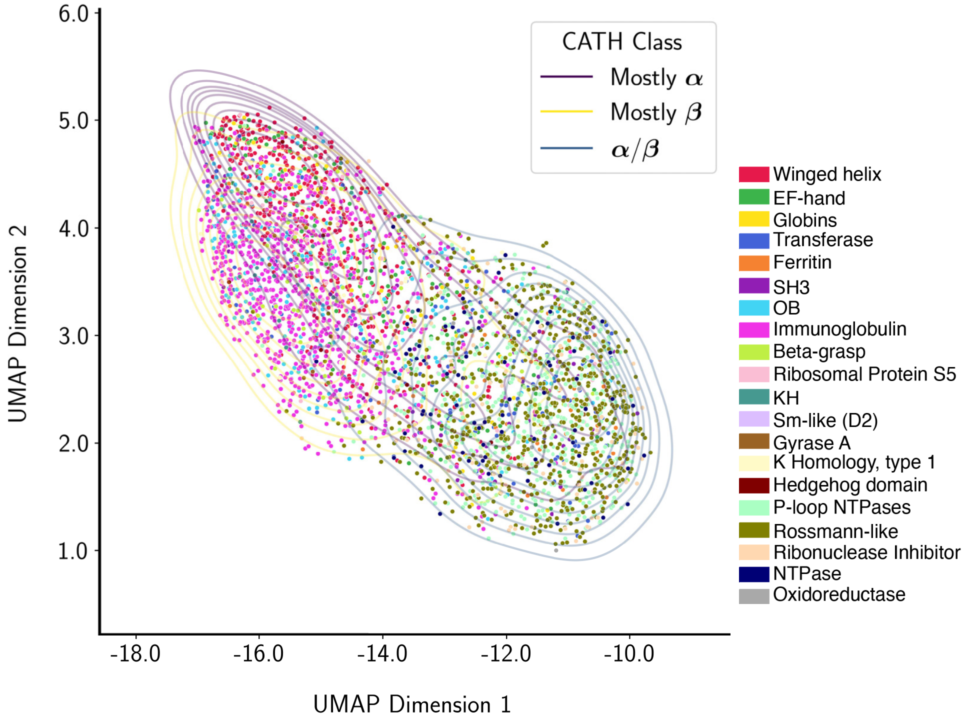

**(c)** Colored by true/correct CATH superfamily

**Supplementary Figure 11: Latent Space from UMAP**: Representatives from each of the 20 SFs were subjected to VAE models trained with domains from the same SF, saving the latent variable representing the mean for each representative domain. The latent variables for each different model were concatenated and reduced from 1,024 to 2 dimensions for visualization purposes as shown here. Domains are colored by (a) secondary structural content, (b) electronegativity, or (c) CATH superfamily.

##### 1.5.1.2 t-SNE

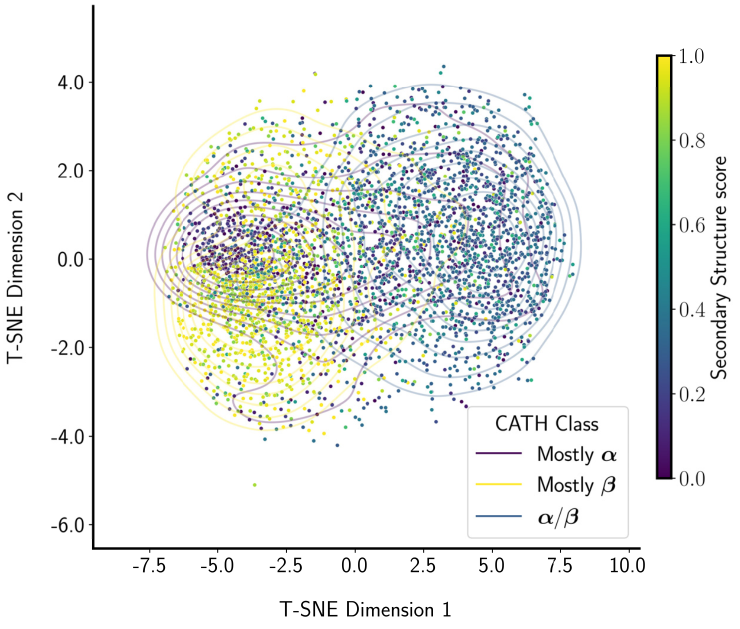

**(a)** Colored by secondary structure content

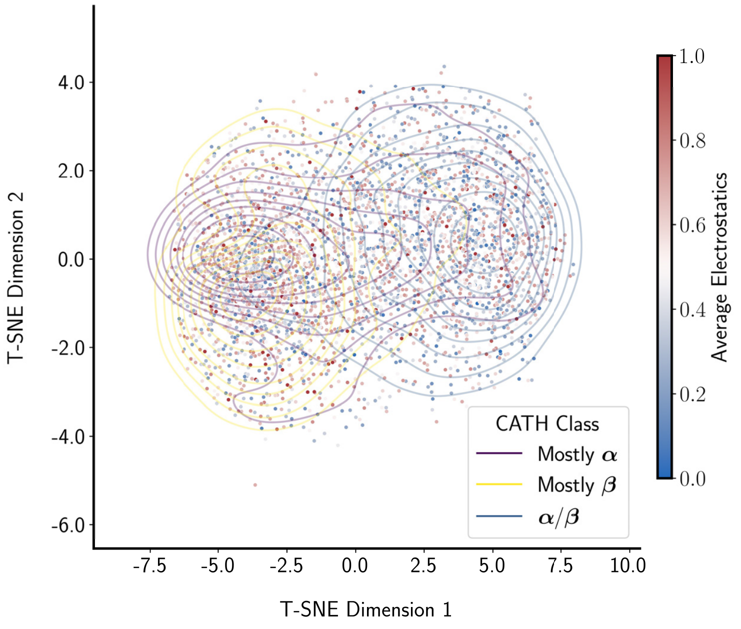

**(b)** Colored by averaged electronegativity values

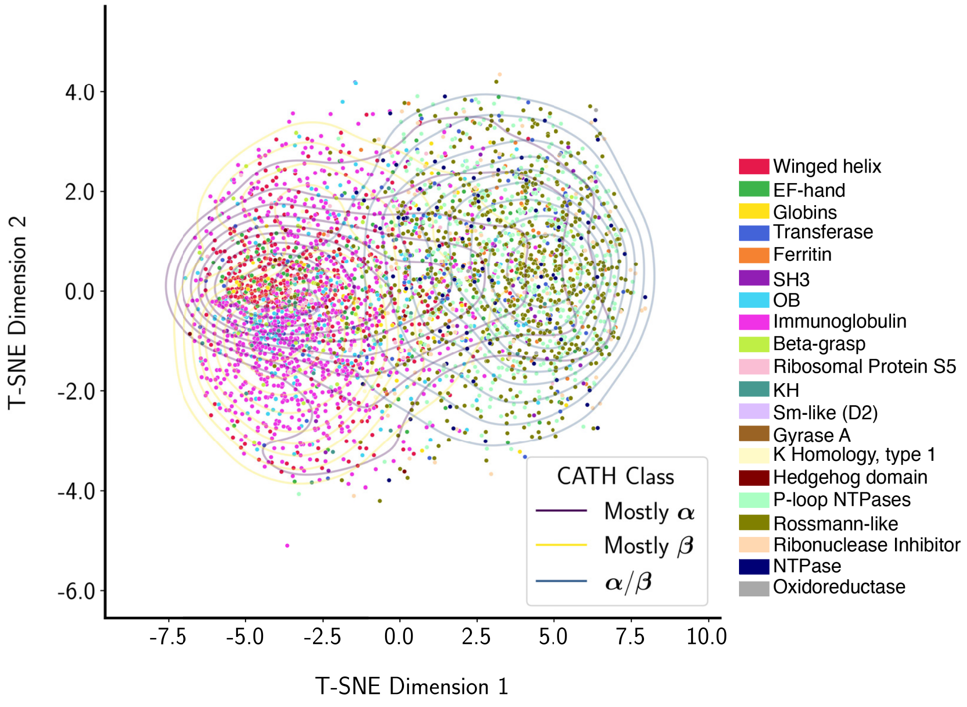

**(c)** Colored by true/correct CATH superfamily

**Supplementary Figure 12: Latent Space from t-SNE**: Analogous to the above UMAP data (Supplementary Figure 11), here we computed subspace projections via the t-distributed Stochastic Neighbor Embedding (t-SNE) algorithm; like UMAP, this dimensionality reduction approach tolerates nonlinearities in the data. As above, domains are colored in these diagrams by either (a) secondary structural content, (b) mean electronegativity, or (c) their CATH superfamily.

##### 1.5.1.3 PCA

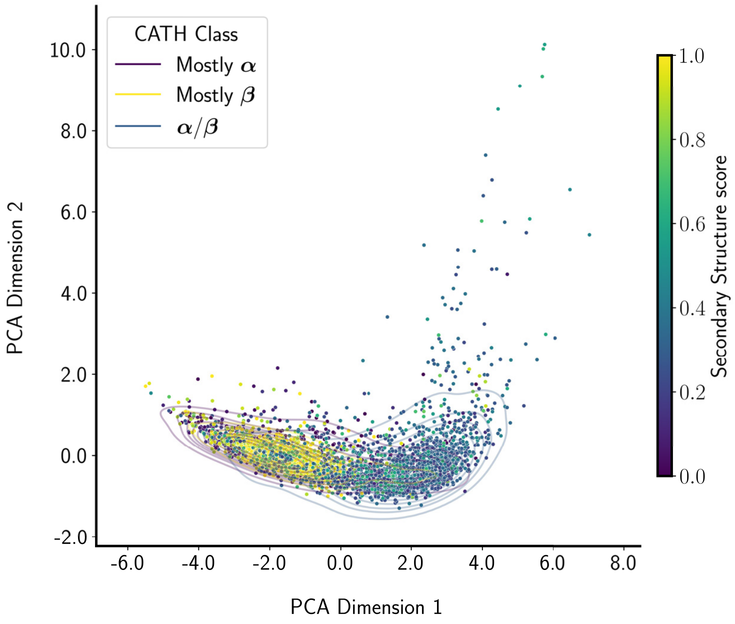

**(a)** Colored by secondary structure content

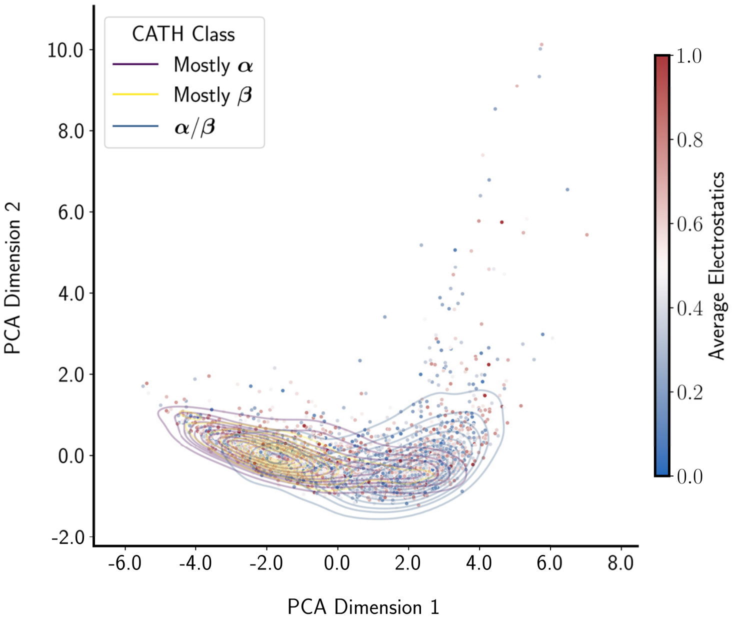

**(b)** Colored by averaged electronegativity values

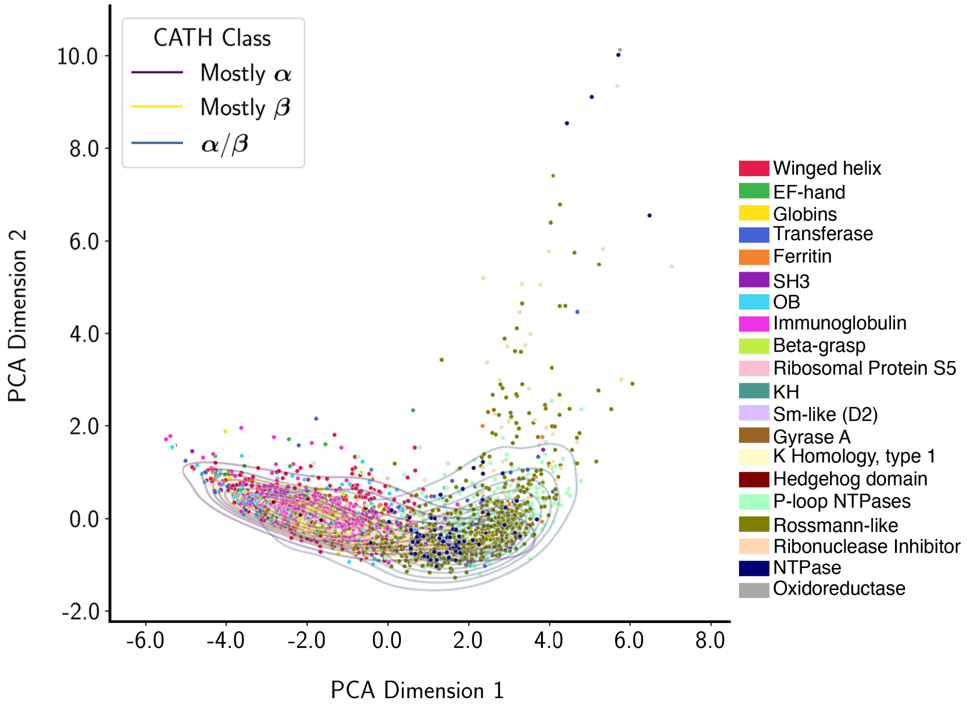

**(c)** Colored by true/correct CATH superfamily

**Supplementary Figure 13: Latent Space from PCA**: These panels, analogous to the preceding two figures, illustrate the results from performing principal component analysis (PCA) on the DeepUrfold latent space embeddings; note that, unlike UMAP and t-SNE, the PCA method cannot account for nonlinearities in the data.

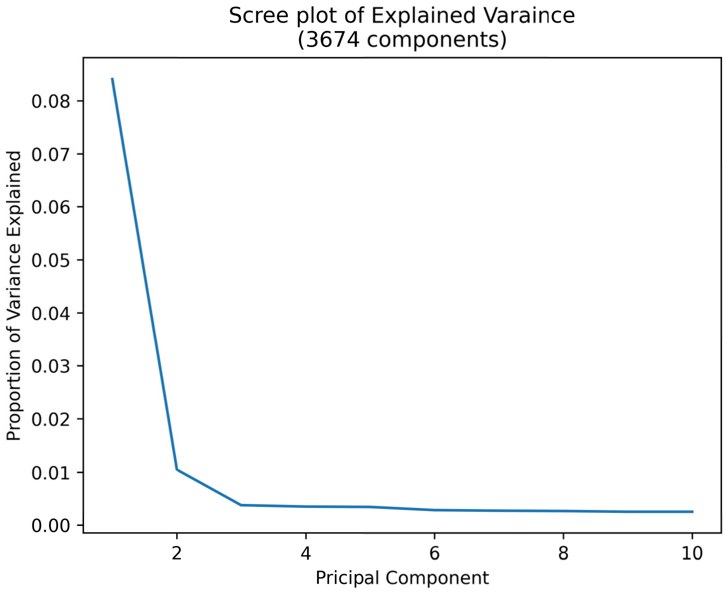

**(a)** Explained Variance of 10 random PCA inverse transforms

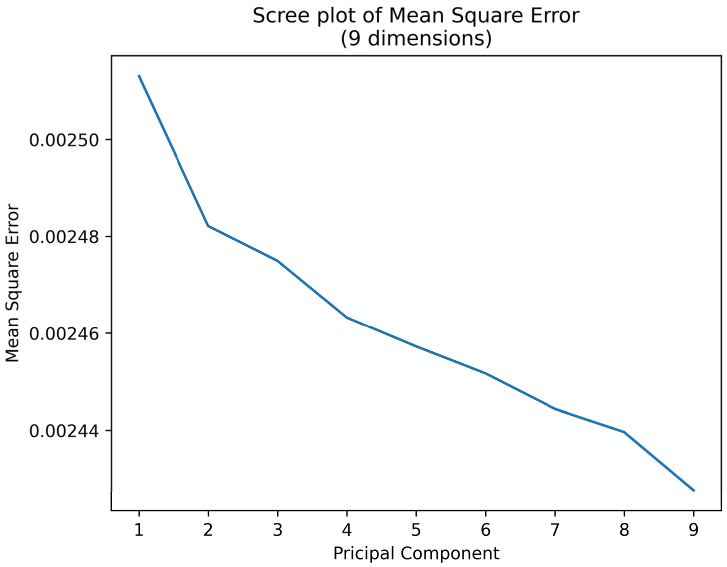

**(b)** Mean Square Error of 10 random PCA inverse transforms

**Supplementary Figure 14: Exploring the PCA results via 'scree' plots**: How much do the principal components account for the model variance? The rapid decay in the scree plot seen here—revealing just one to two components as accounting for the bulk of the variance—is consistent with the general 'streaky' appearance of the PCA projections (e.g., Supplementary Figure 17 for the Global Model). This is because a streaky appearance means that the underlying high-dimensional data can be mapped to an essentially one-dimensional manifold (e.g., Supplementary Figures 13 and 17 panels, "Latent Space from PCA", and to some degree the UMAP results as well [see the Supplementary Discussion §5.3.1, "Individual, Superfamily-level Feature Embeddings via UMAP", regarding some of the 'streaky' 2D mappings in Supplementary Figure 18's SF-level UMAP subpanels]).

##### 1.5.2 Global Model

###### 1.5.2.1 UMAP

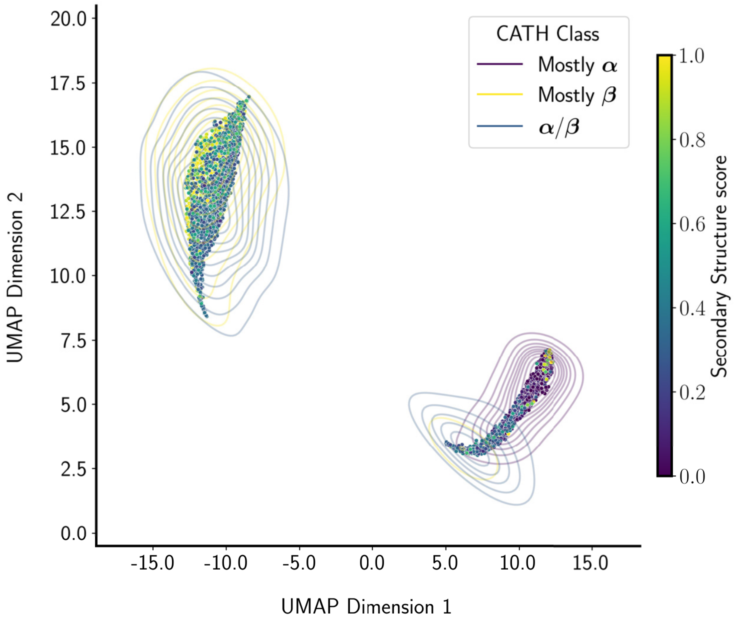

**(a)** Colored by secondary structure content

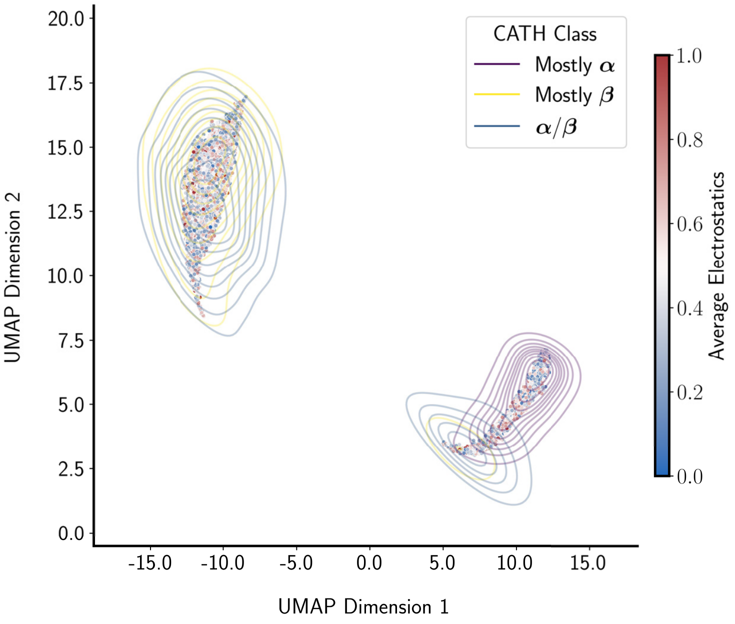

**(b)** Colored by averaged electronegativity values

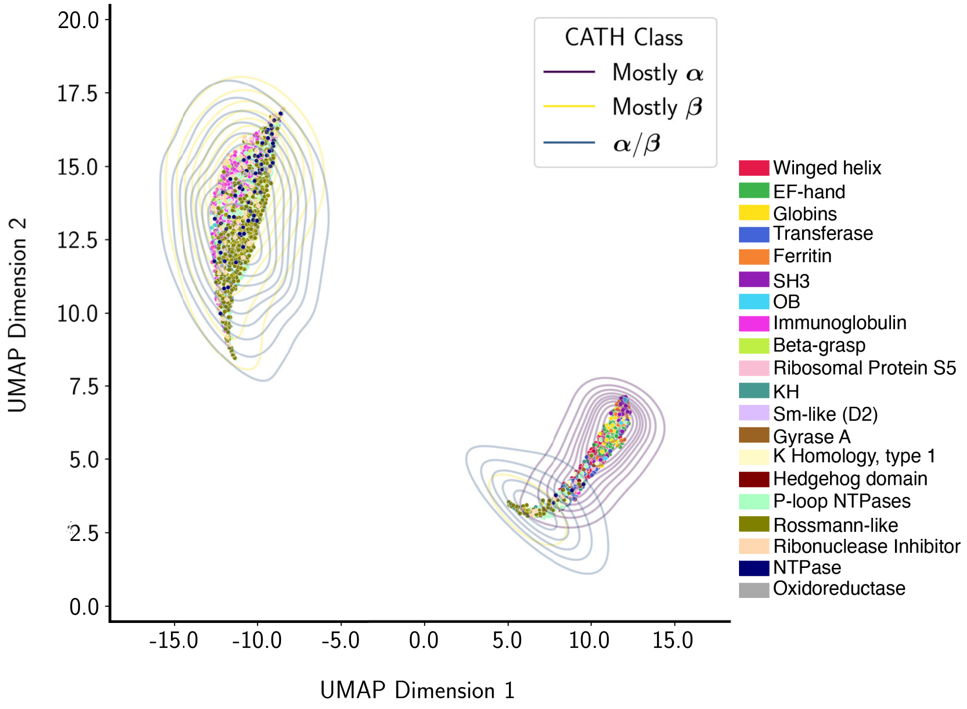

**(c)** Colored by true/correct CATH superfamily

**Supplementary Figure 15: Latent Space from UMAP**: Representatives from each SF were subjected to the Global Model, which was trained on a single set of all domains from every superfamily, saving the latent variable representing the mean for each representative domain. The latent variables for each different model were reduced from 1,024 to 2 dimensions for the visualization purposes illustrated here.

##### 1.5.2.2 t-SNE

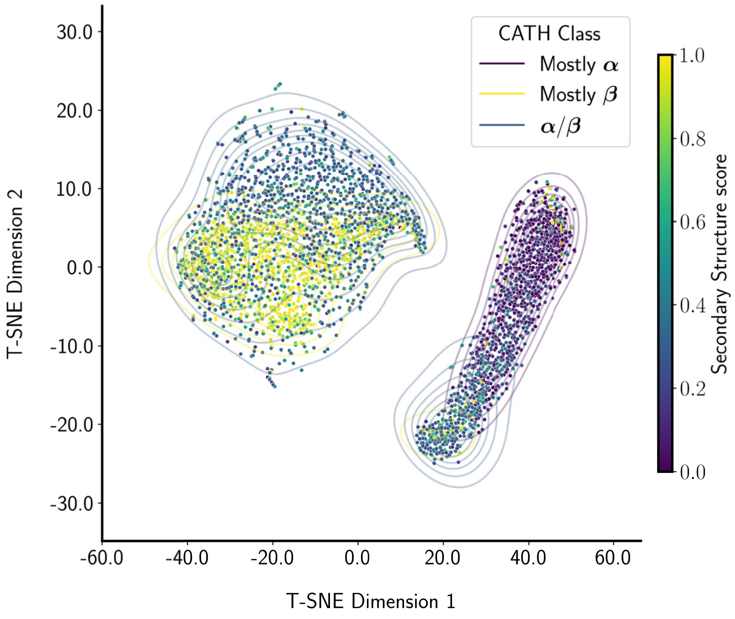

**(a)** Colored by secondary structure content

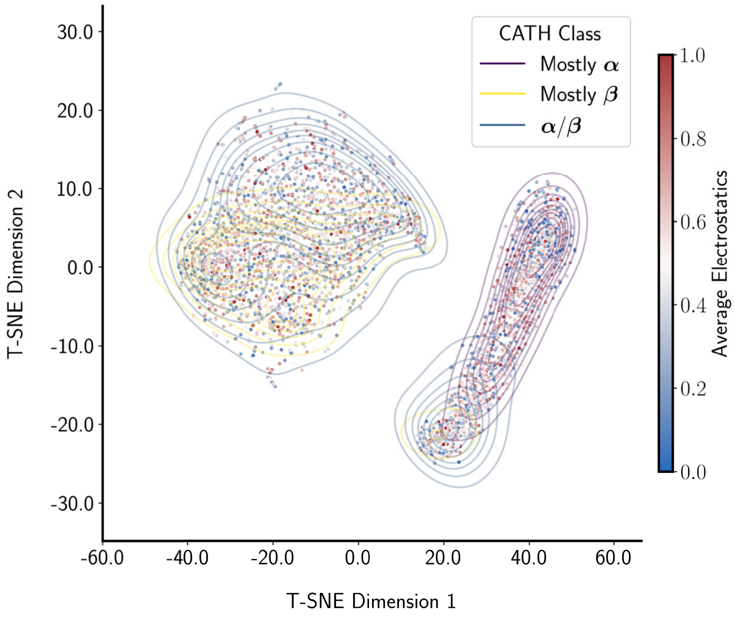

**(b)** Colored by averaged electronegativity values

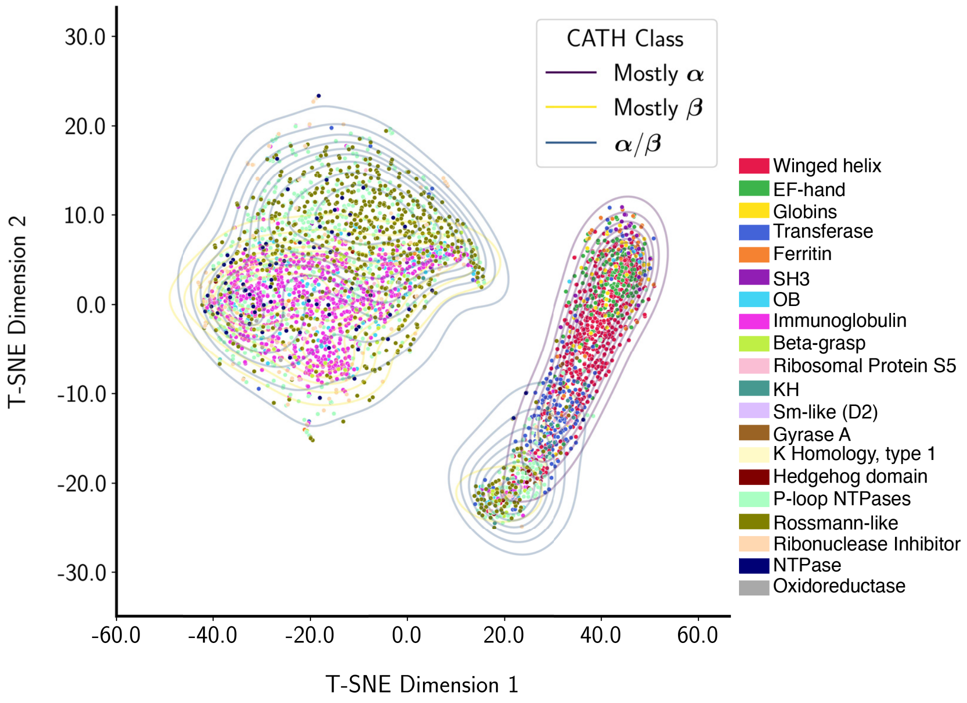

**(c)** Colored by true/correct CATH superfamily

**Supplementary Figure 16: Latent Space from t-SNE**: Analogous to the above UMAP data (Supplementary Figure 15), here we computed subspace projections via the t-distributed Stochastic Neighbor Embedding (t-SNE) algorithm; like UMAP, this dimensionality reduction approach tolerates nonlinearities in the data.

##### 1.5.2.3 PCA

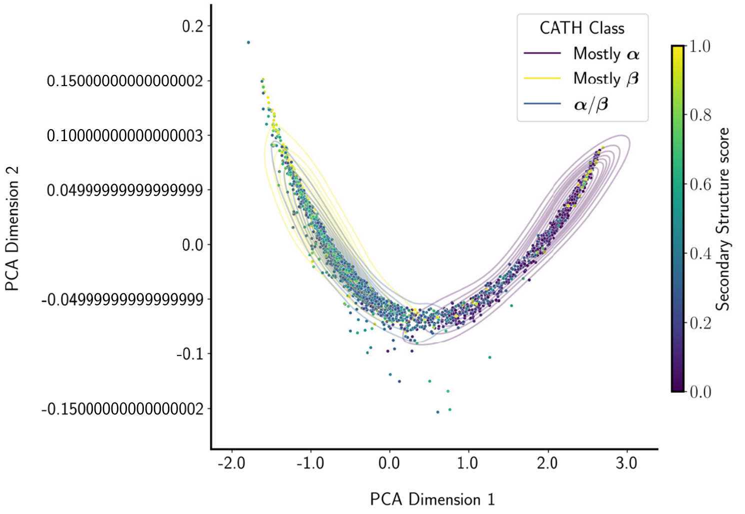

**(a)** Colored by secondary structure content

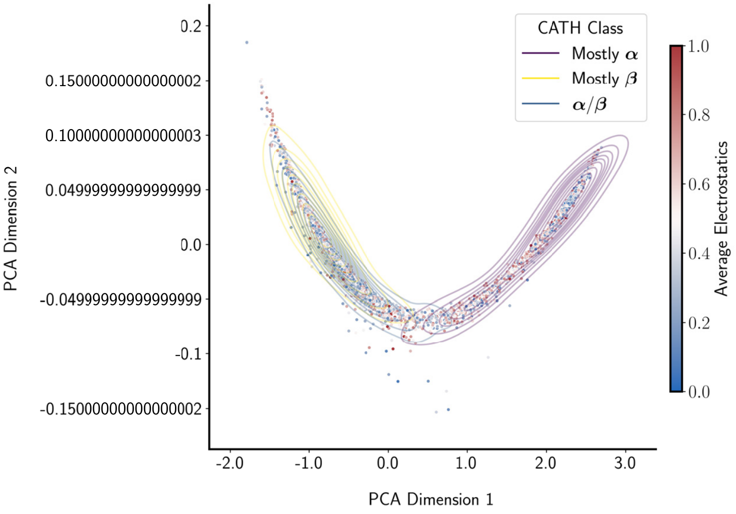

**(b)** Colored by averaged electronegativity values

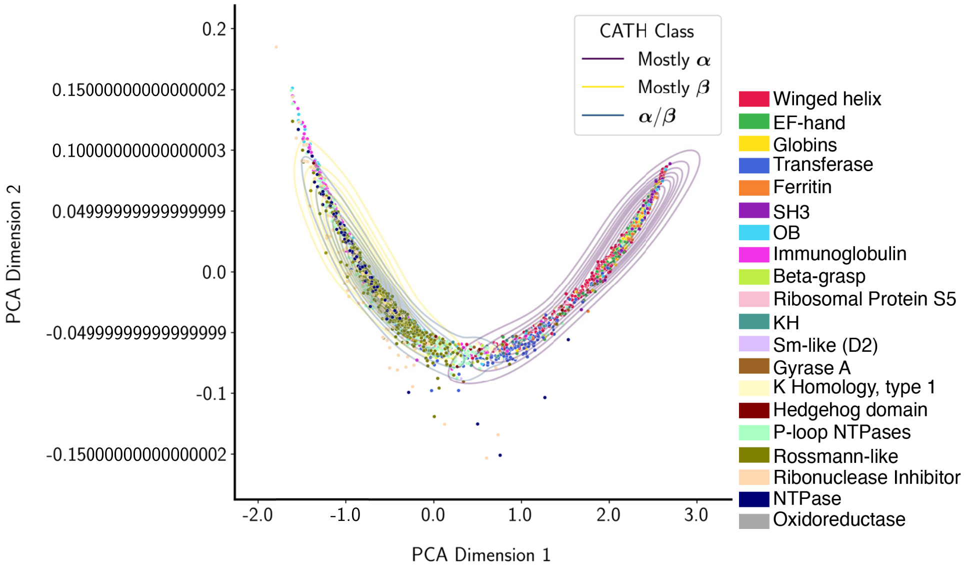

**(c)** Colored by true/correct CATH superfamily

**Supplementary Figure 17: Latent Space from PCA**: These panels, analogous to the preceding two figures, illustrate the results from performing PCA on the DeepUrfold latent space embeddings; as mentioned above, note that, unlike UMAP and t-SNE, PCA does not account for nonlinearities in the data.

##### 1.5.3 Individual, Superfamily-level Feature Embeddings via UMAP

**Supplementary Figure 18:** Instead of concatenating the latent space embeddings for each representative domain from the 20 SF-level models along the feature dimension, in the following panels we reduced the latent space with UMAP separately for each model. Note that these results also show similar gradients of secondary structure content for each learned model.

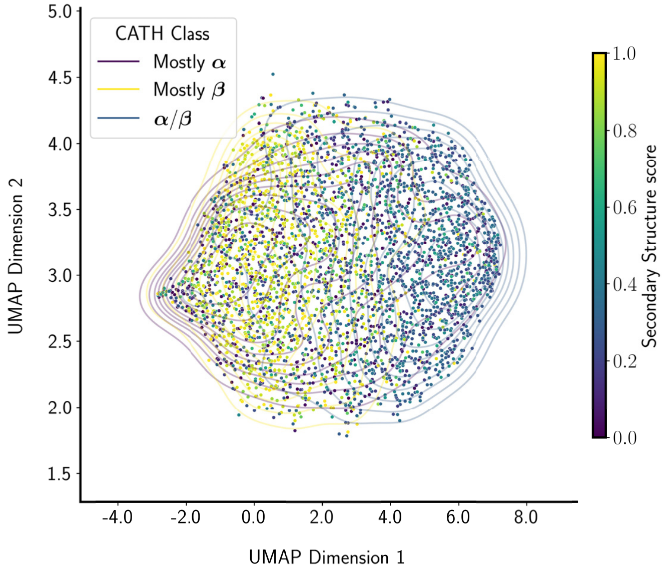

(a) 1.10.10.10

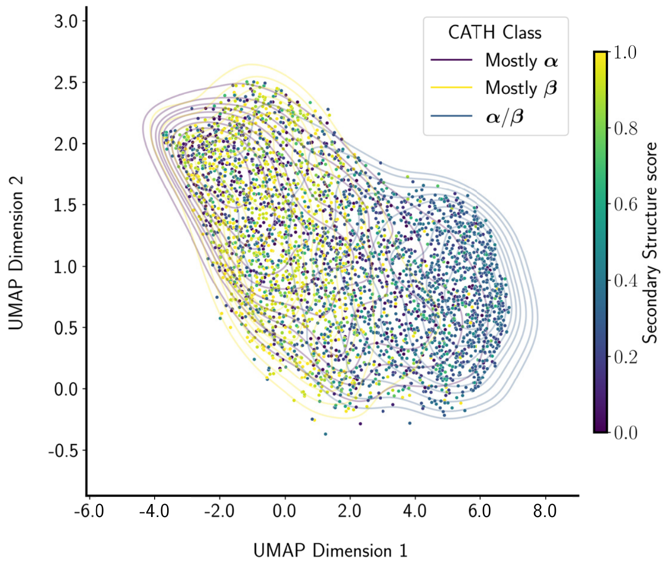

(b) 1.10.238.10

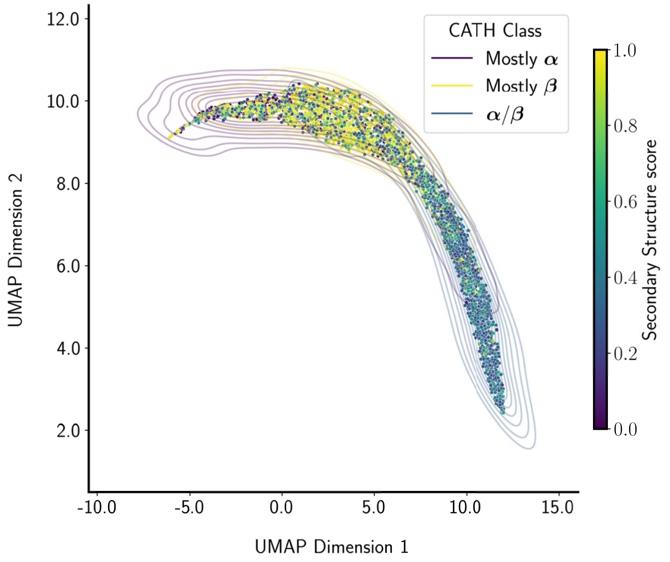

(c) 1.10.490.10

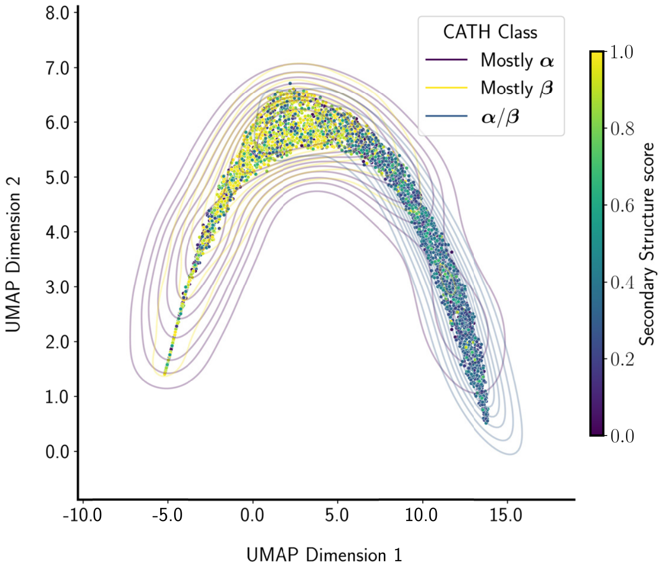

(d) 1.10.510.10

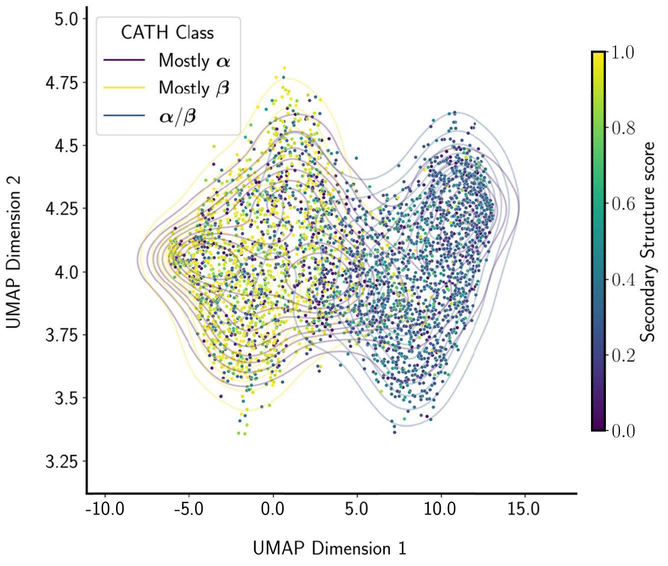

**(e)** `1.20.1620.10`

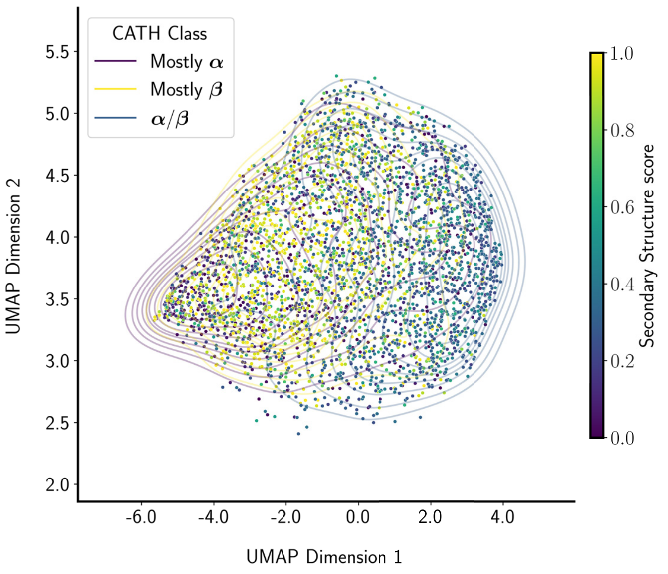

**(f)** `2.30.30.100`

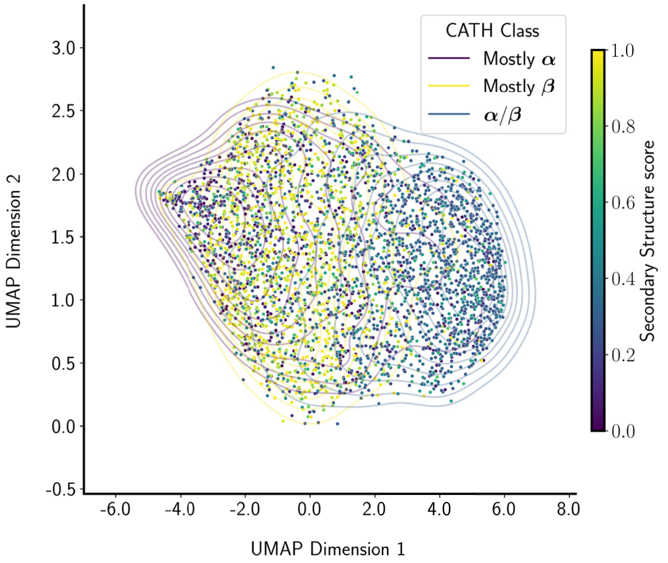

**(g)** `2.40.50.140`

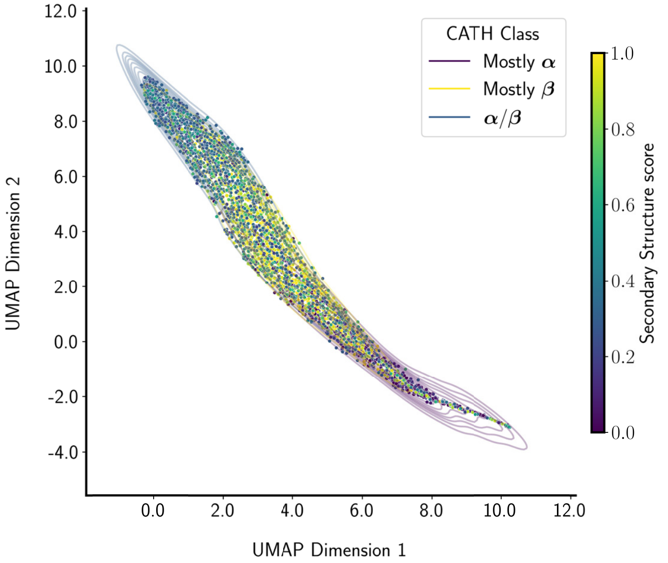

**(h)** `2.60.40.10`

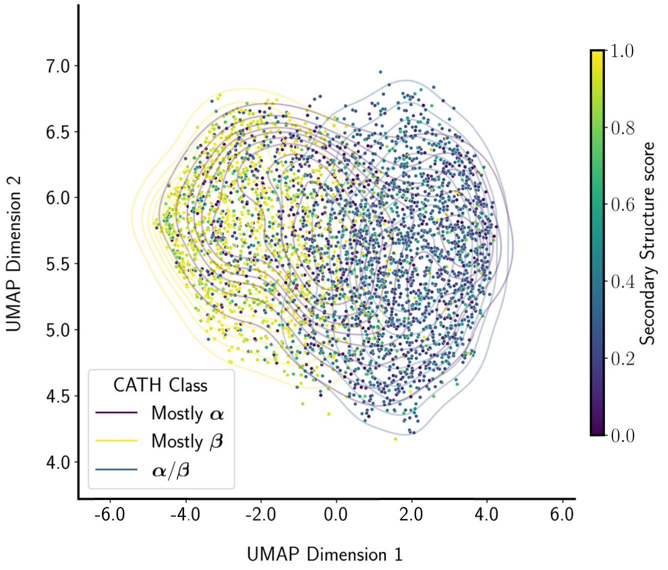

**(i)** `3.10.20.30`

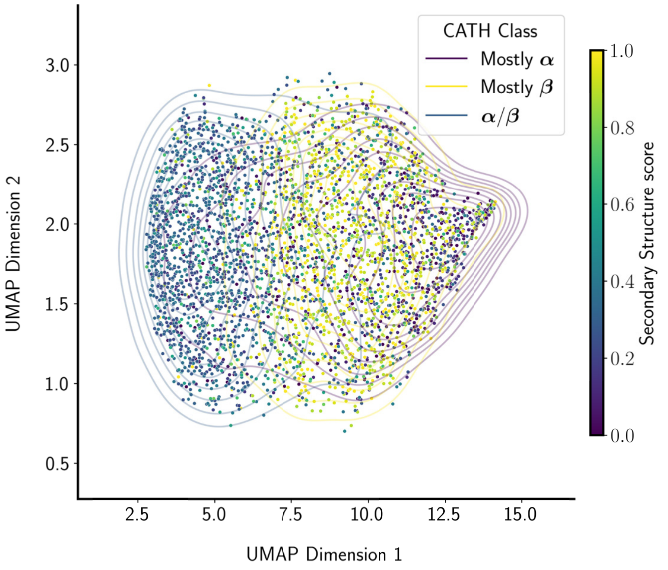

**(j)** `3.30.230.10`

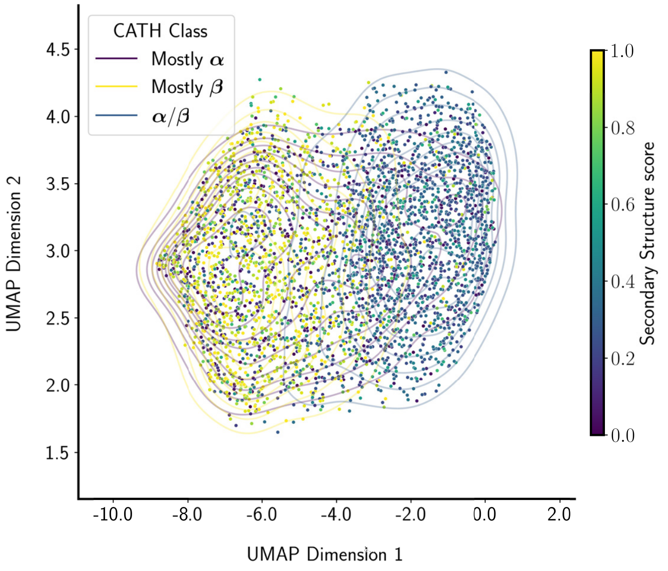

**(k)** `3.30.300.20`

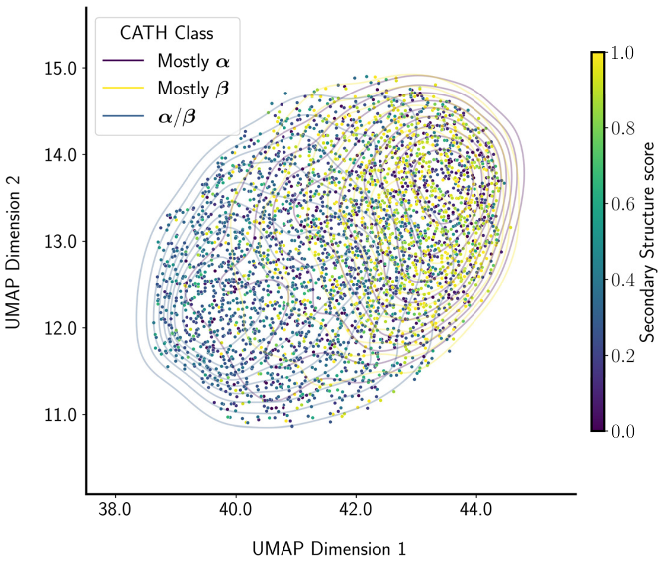

**(l)** `3.30.310.60`

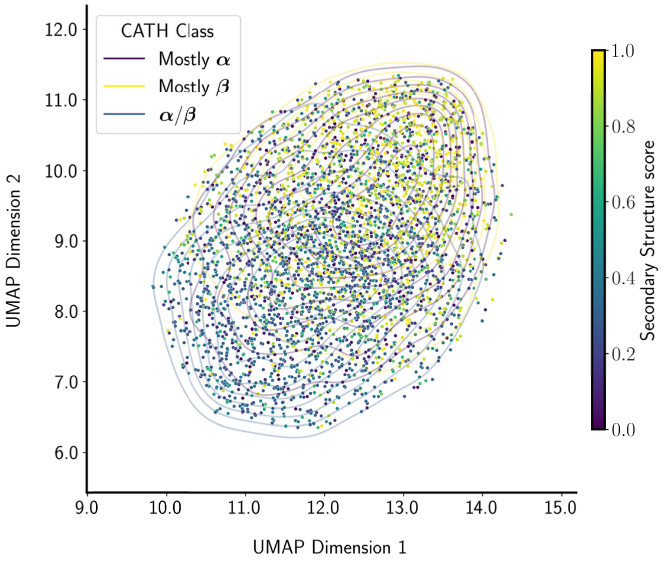

**(m)** `3.30.1360.40`

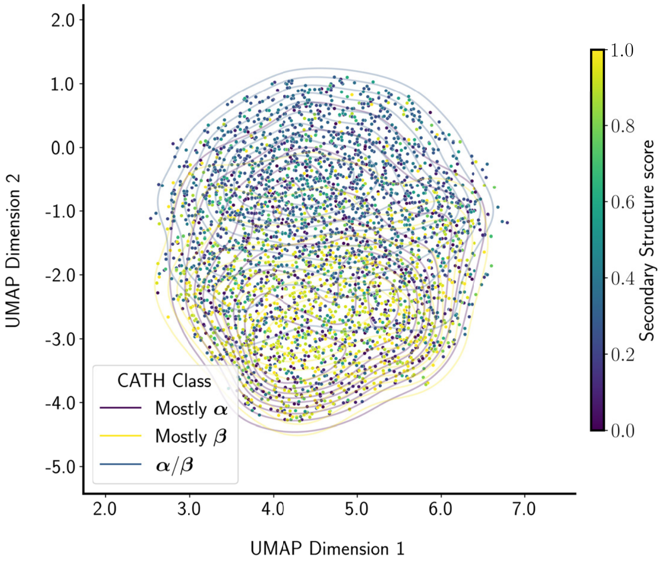

**(n)** `3.30.1370.10`

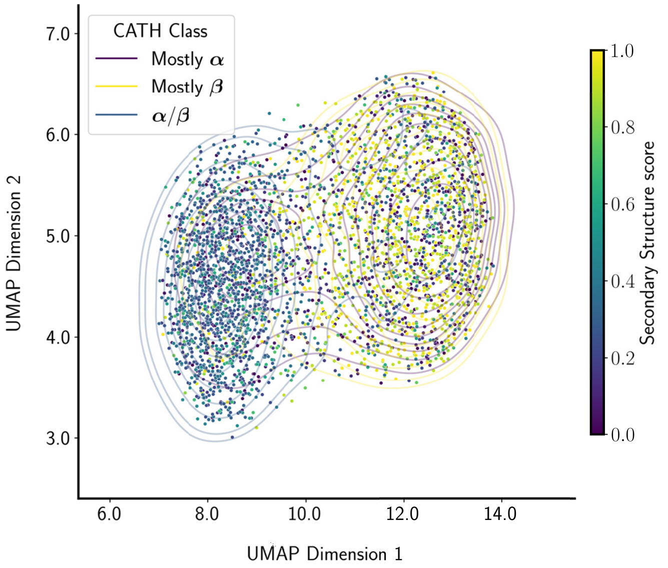

**(o)** `3.30.300.20`

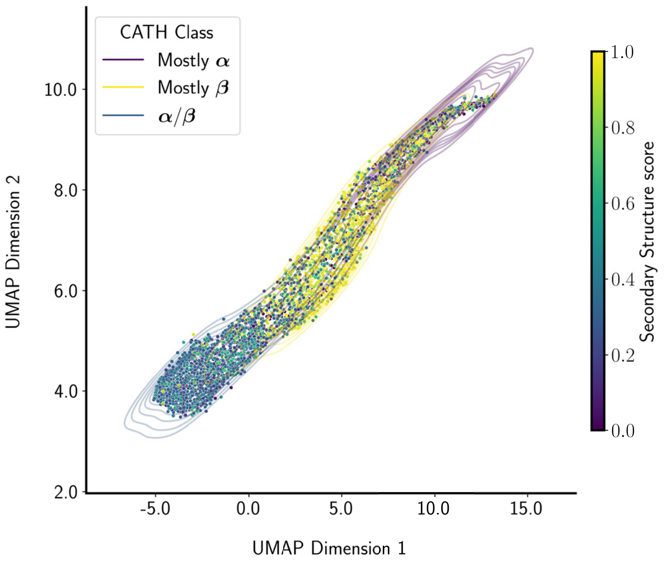

**(p)** `3.40.50.300`

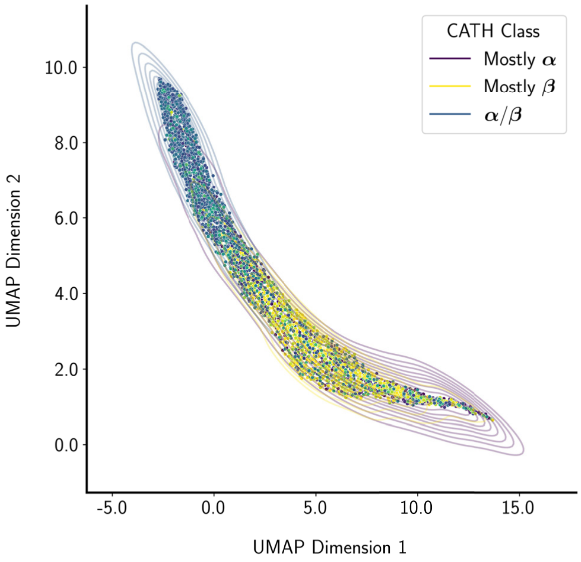

**(q)** `3.40.50.720`

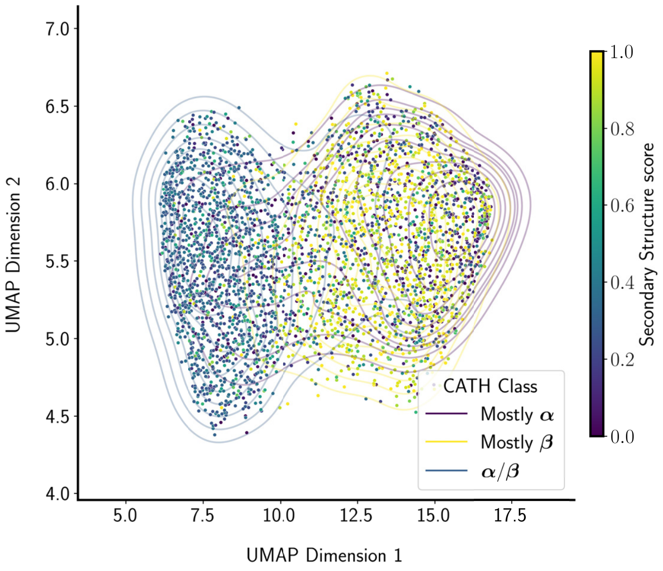

**(r)** `3.80.10.10`

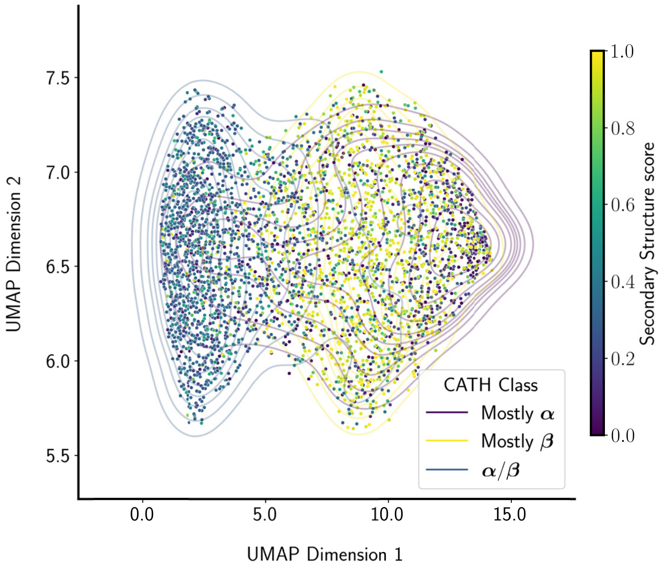

**(s)** `3.90.79.10`

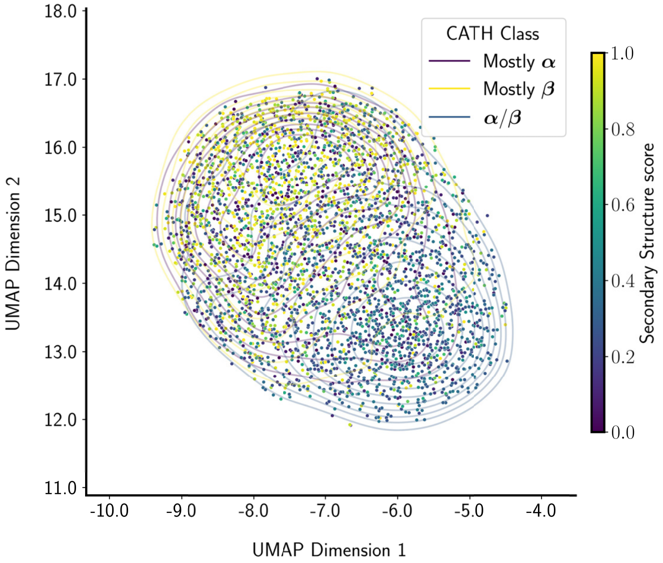

**(t)** `3.90.420.10`

##### 1.5.4 Projected Latent Space Embeddings after Alignment via Optimal Transport

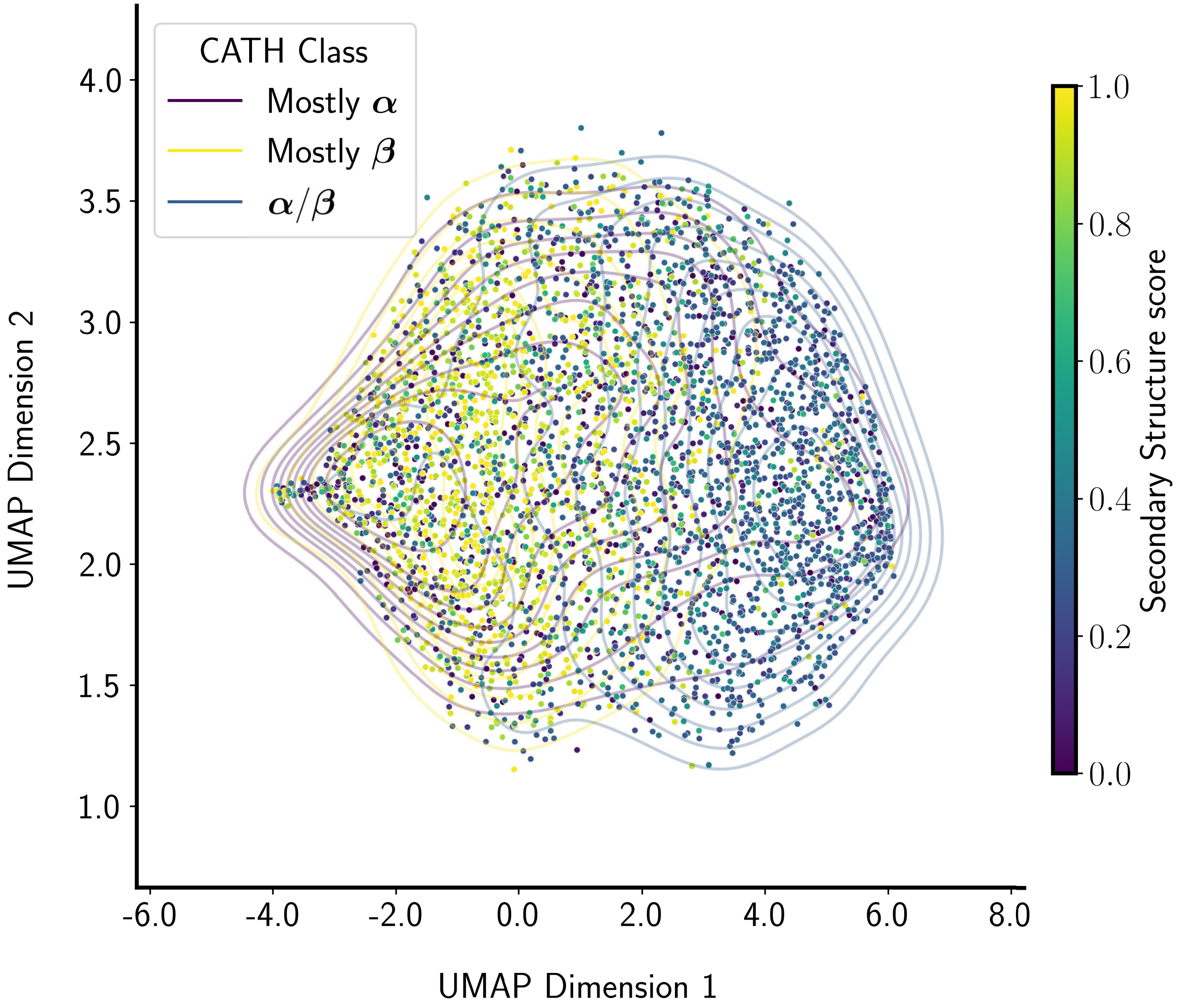

**Supplementary Figure 19:** The 20 different SF embeddings can each occupy distinct regions of latent space and therefore cannot necessarily be directly compared. To address this possible issue, in this approach we first transform ('shift') all 20 different SF latent space embeddings into a common region of the hyperspace via application of an Optimal Transport (OT) Domain Adaptation algorithm. In particular, we performed a Sinkhorn-based transport with group LASSO L1L2 regularization of OT distances (using POT, the Python Optimal Transport package), concatenating all embeddings in the feature dimension. Applying this OT algorithm enabled us to 'transport' each domain's latent space embedding to the single space of CATH SF `1.10.10.10`; that particular SF, the first in our list, was chosen simply because it was found to exhibit a fairly clean secondary structure gradient in our other (non–OT-based) UMAP calculations (see, e.g., Supplementary Figure 18a; also, note that this choice should be relatively immaterial, given the application of OT). Next, to these optimally transported embeddings we applied the same approach as described elsewhere in this text, wherein we concatenated all transported latent spaces on the column dimension for each representative domain. Finally, we computed a UMAP reduction on this monolithic transported-and-concatenated dataset. Note that OT has been shown to be effective in comparing different embedding spaces (see, e.g., Alvarez-Melis et al.'s "*Towards Optimal Transport with Global Invariances*" [3]).

##### 1.5.5 Rotational Invariance of Trained Models

###### 1.5.5.1 SH3-only Model (a single-SF model)

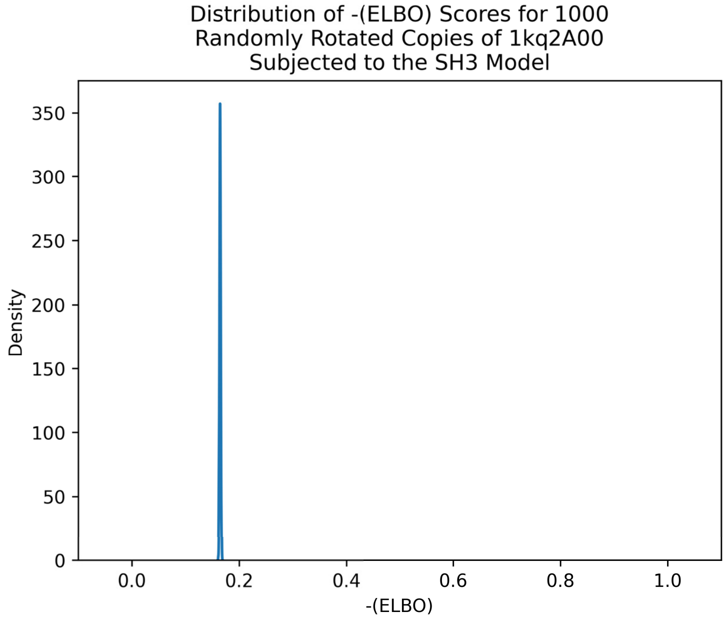

**(a)** Distribution of –(ELBO) scores for rotated copies

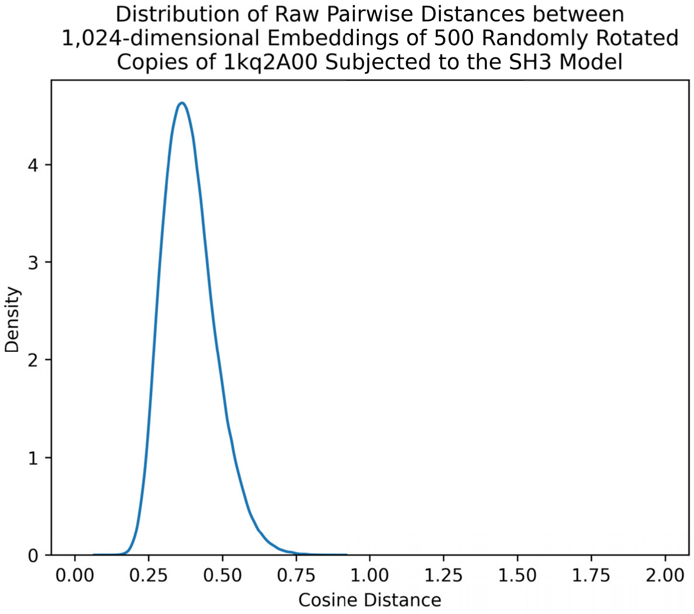

**(b)** Distribution of raw embedding distances between rotated structures subjected to inference passes through the SH3 model

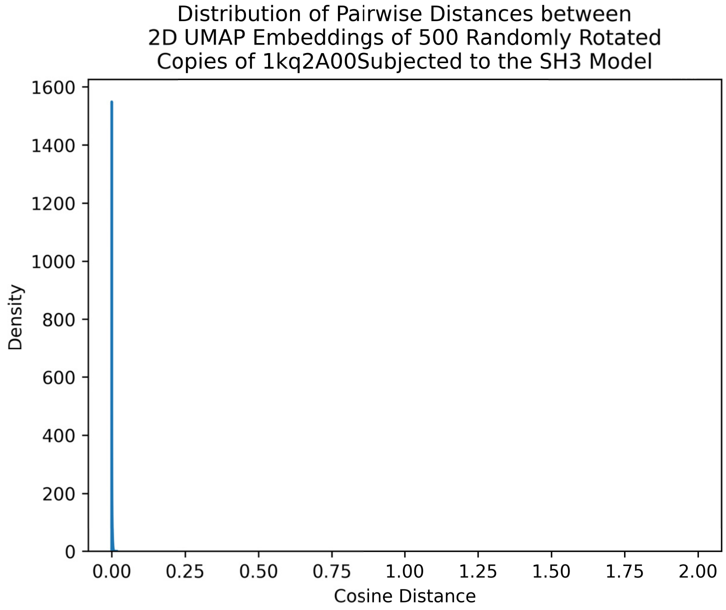

**(c)** Distribution of UMAP embedding distances between rotated structures subjected to the SH3 model

**Supplementary Figure 20:** To test if our single-superfamily trained VAE models (for the SH3, in this case) are rotation invariant, we subjected random rotations (drawn from the SO(3) group) of a canonical SH3 member (1kq2A00) to the VAE model trained on all SH3 domains. The distributions of (a) –(ELBO) scores, (b) pairwise distances between 1,024-dimensional embeddings (using the cosine distance), and (c) 2D UMAP cosine distances all exhibit small means and variances (i.e., tight distributions). This finding indicates that DeepUrfold's VAE training yielded a model that finds all angular orientations of the 3D structure to be similar (i.e., the model is not learning protein orientation).

###### 1.5.5.2 Concatenated Model

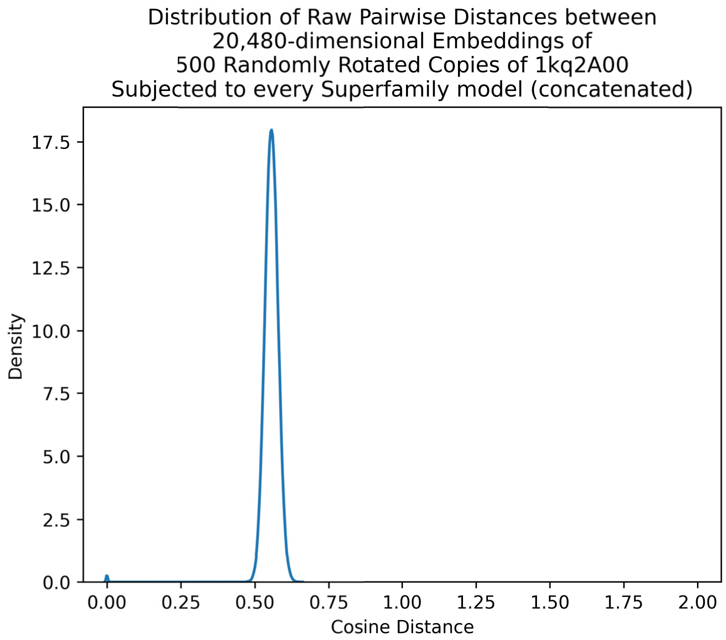

**(a)** Raw embedding distances between rotated structures subjected to the concatenated model

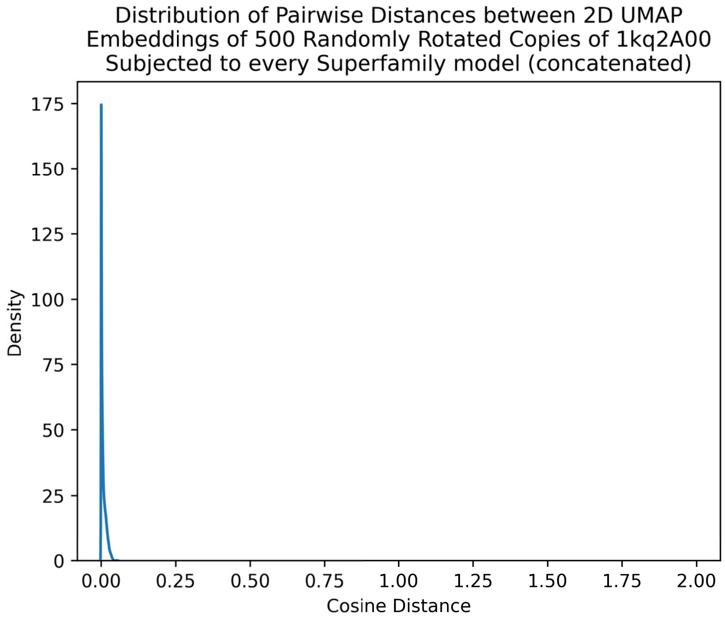

**(b)** UMAP embedding distances between rotated structures subjected to the concatenated model

**Supplementary Figure 21:** In order to test if our Concatenated Model is rotation invariant, we subject random rotations (from the SO(3) group) of 1kq2A00, a canonical SH3 member, to each superfamily model, followed by concatenating their embeddings as described in the Methods section. The distributions around the (a) pairwise cosine distances among the 20,480-dimensional embedding vectors and (b) 2D UMAP cosine distances both exhibit small means and variances (i.e., tight distributions), showing that the Concatenated Model scores all 3D structural rotations similarly—i.e., it is rotationally invariant. Note this figure's absence of a panel showing a 'Distribution of –(ELBO) scores for rotated copies', unlike for the SH3-only Model and the Global Model (cf. Supplementary Figures 20 and 22); this is the case because the embedding vectors are concatenated in a post-training manner (see Methods), meaning there are not raw –(ELBO) scores for this model formulation.

###### 1.5.5.3 Global Model

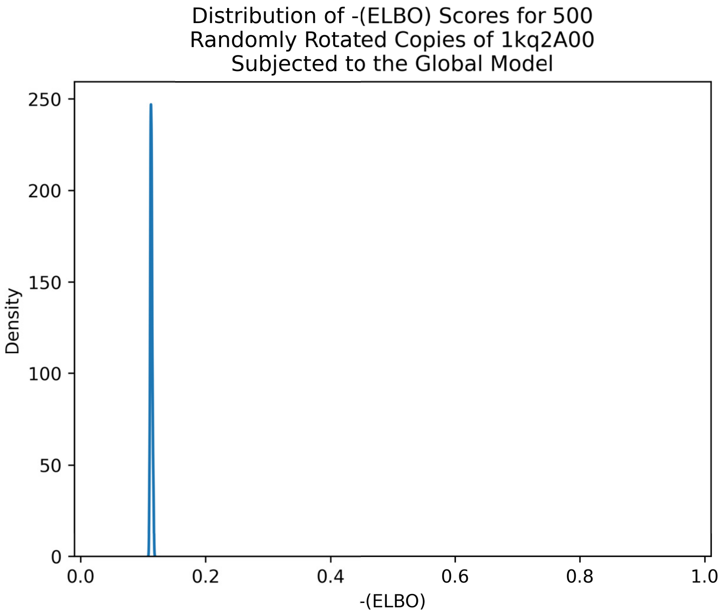

**(a)** Distribution of –(ELBO) scores for rotated copies

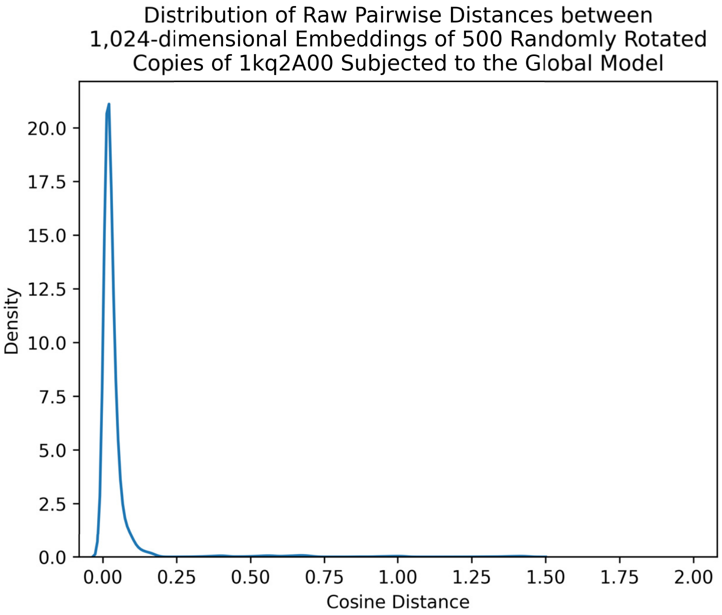

**(b)** Raw embedding distances between rotated structures subjected to the SH3 model

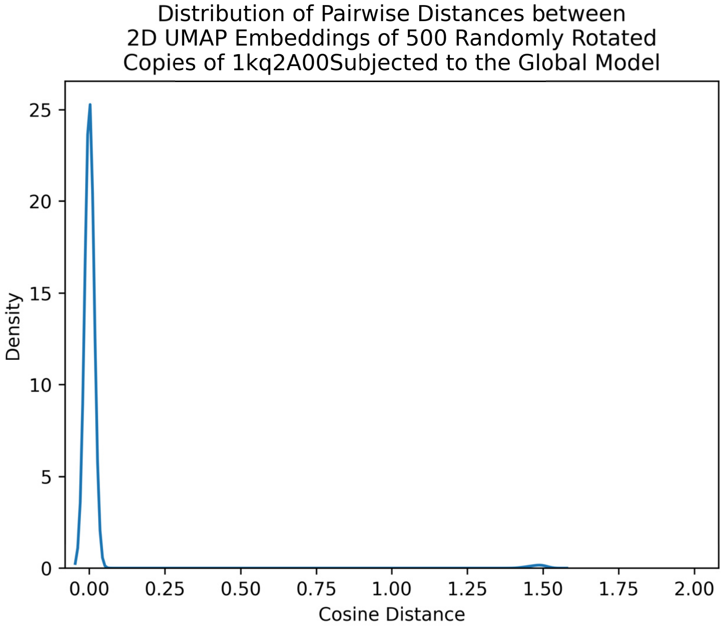

**(c)** UMAP embedding distances between rotated structures subjected to the SH3 model

**Supplementary Figure 22:** Finally, to test if our Global Model is also rotation-invariant, we rotated a canonical SH3 (1kq2A00) into 500 random orientations (drawn from the SO(3) group; see Methods) and subjected each of those rotated copies to a monolithic VAE model trained on all domains from every superfamily (i.e., the new Global Model). The distributions of (a) raw –(ELBO) scores, (b) 1,024-dimensional embedding cosine distances, and (c) 2D UMAP cosine distances all exhibit small means and variances (i.e., tight distributions), showing that all rotations are scored similarly by the Global Model, and implying that its learned representations are rotationally invariant.

##### 1.5.6 Patterns of Proximity among Latent Space Embeddings

###### 1.5.6.1 SH3-only Model

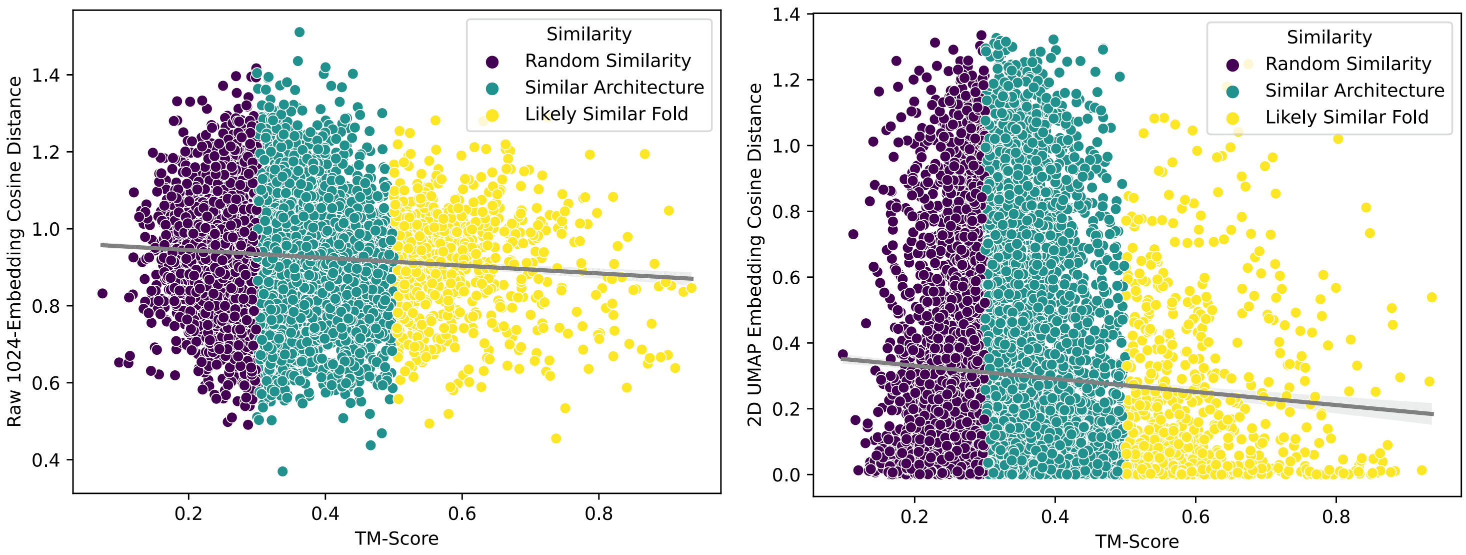

**(a)** Raw 1,024-dimensional Embedding Distances vs TM-score

**(b)** 2D UMAP Embedding Distances vs TM-score

**Supplementary Figure 23:** In both (a) raw and (b) UMAP-reduced embeddings distances, there is a somewhat weak negative correlation between (i) pairwise distances between embeddings ($y$-axis) and (ii) pairwise TM-scores for the SH3-only model ($x$-axis).

###### 1.5.6.2 Concatenated Model

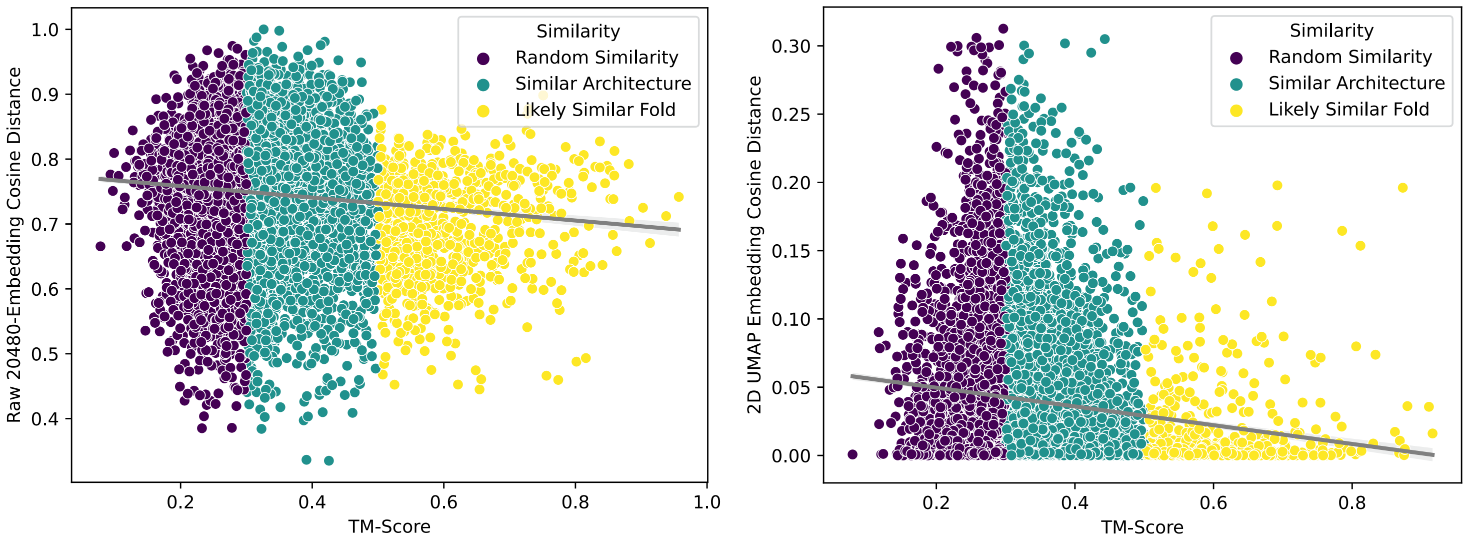

**(a)** Raw 20,480-dimensional Embedding Distances vs TM-score

**(b)** 2D UMAP Embedding Distances vs TM-score

**Supplementary Figure 24:** In both (a) raw and (b) UMAP-reduced embeddings distances, there is a somewhat weak negative correlation between embedding distances and TM-score values for the Concatenated Model (see Methods for its construction).

###### 1.5.6.3 Global Model

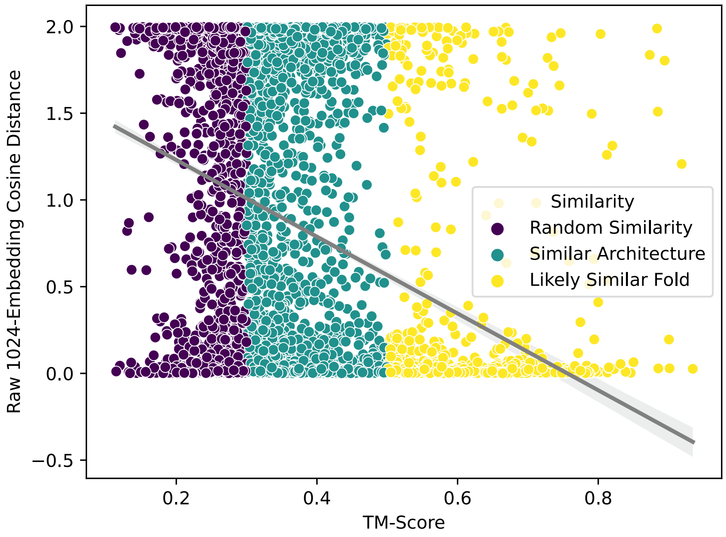

**(a)** Raw 1,024-dimensional Embedding Cosine Distances vs TM-score

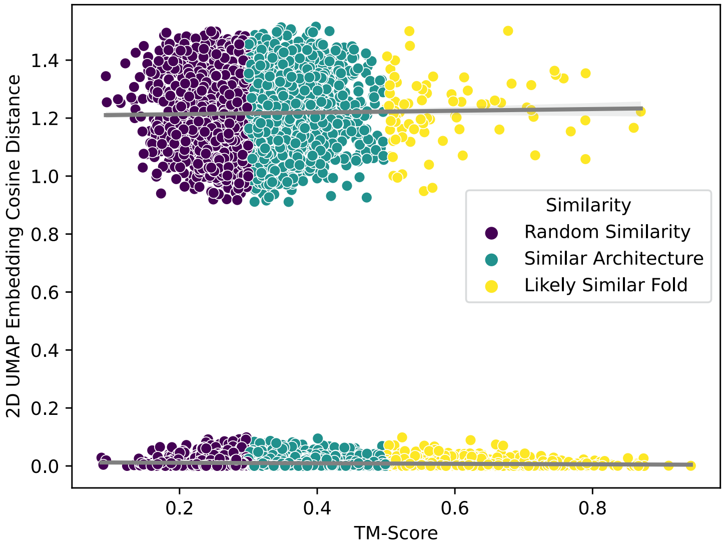

**(b)** 2D UMAP Embedding Cosine Distances vs TM-score

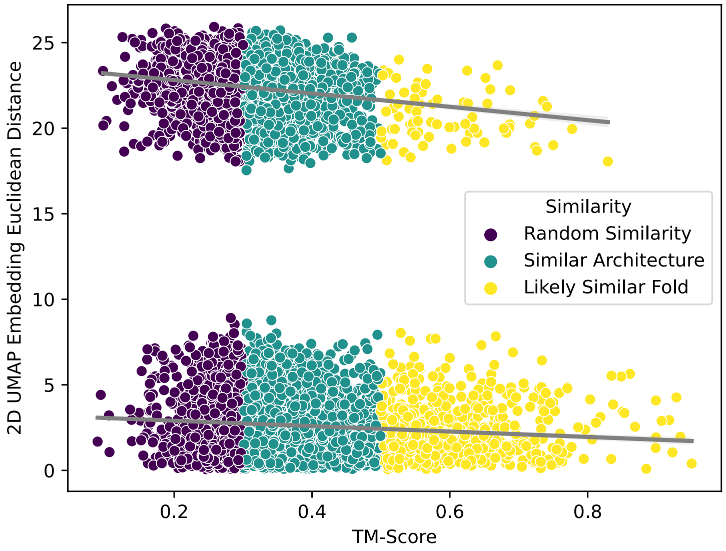

**(c)** 2D UMAP Embedding Euclidean Distances vs TM-score

**Supplementary Figure 25:** In both (a) raw and (b-c) UMAP-reduced embeddings distances, there is a somewhat weak negative correlation between the distances and TM-scores for the Global Model. We show both the cosine (b) and Euclidean (c) distances because the regression line of the cosine-distance–based graph was flatter compared to the Euclidean-distance–based graph; the Euclidean-distance–based calculation yields a slightly steeper slope (stronger correlation).

#### 1.6 Stochastic Block Modelling (SBM) of DeepUrfold-based Communities

##### 1.6.1 Radial Tree of Groupings/Communities

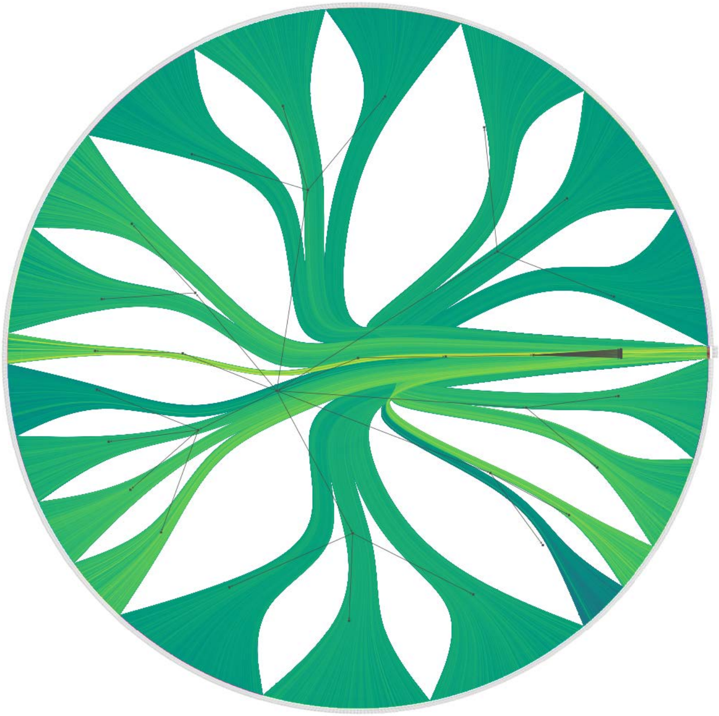

**Supplementary Figure 26:** This radial tree of computed groupings is based on applying SBM methods to DeepUrfold's –log(–ELBO)–weighted, all-by-all bipartite graph (Figure 1C). The –log(–ELBO) weight is shown as the edge color using the viridis scale, where larger values are more yellow and lower values more blue. A searchable, higher quality version of this tree can be found in our Zenodo archive (https://doi.org/10.5281/zenodo.13228646) under 'sbm_results'. Also of note, all SF-specific VAE models were found to group together in the same community, seen here at 3 o'clock (it is not included in circle-packing diagrams nor the # of clusters count [including this group would make for 21 circles]).

##### 1.6.2 Electrostatics

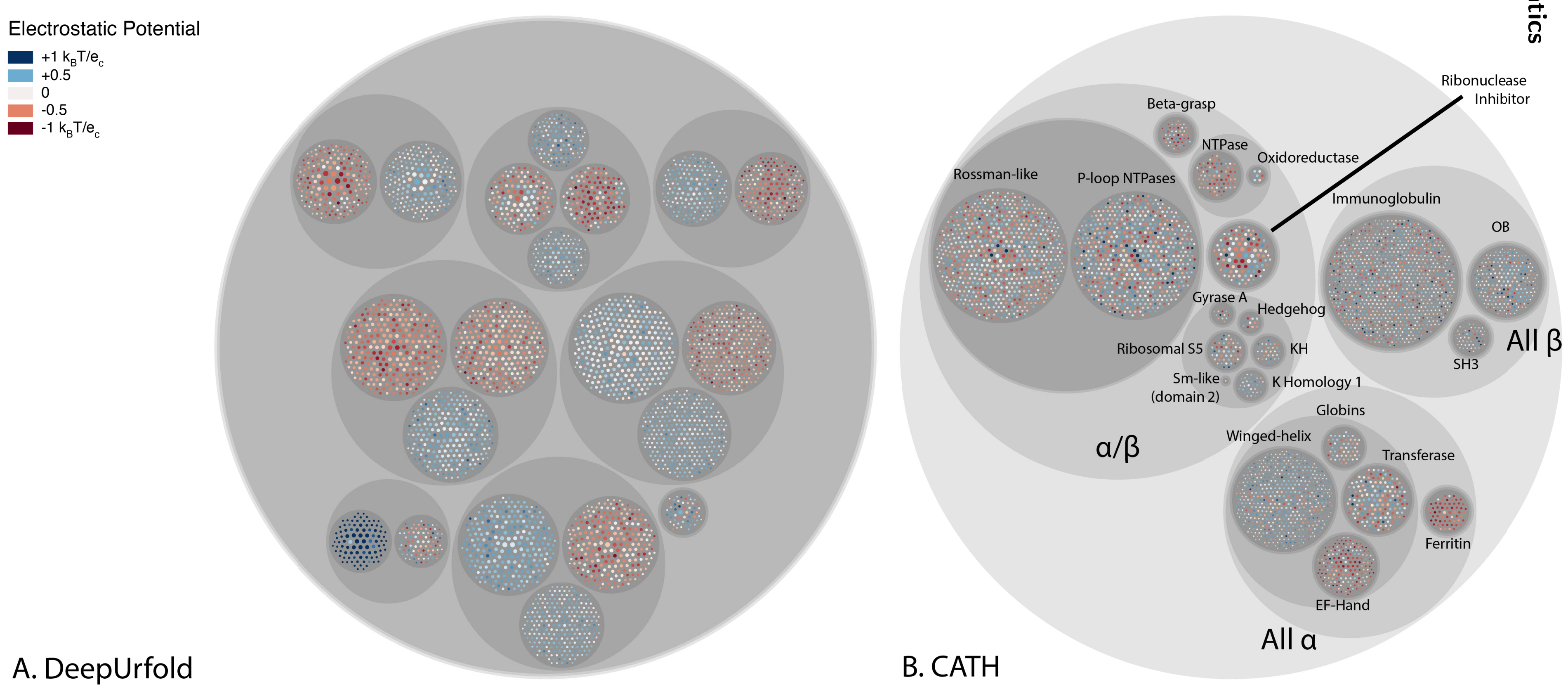

**Supplementary Figure 27:** Each atom is annotated with the Boolean feature is_electronegative. We sum up all of the electronegative atoms and take a fraction of the total number of atoms. Note an intriguing ‘cohesion’ exhibited by the DeepUrfold groupings (A), relative to CATH’s classifications (B).

##### 1.6.3 Charge

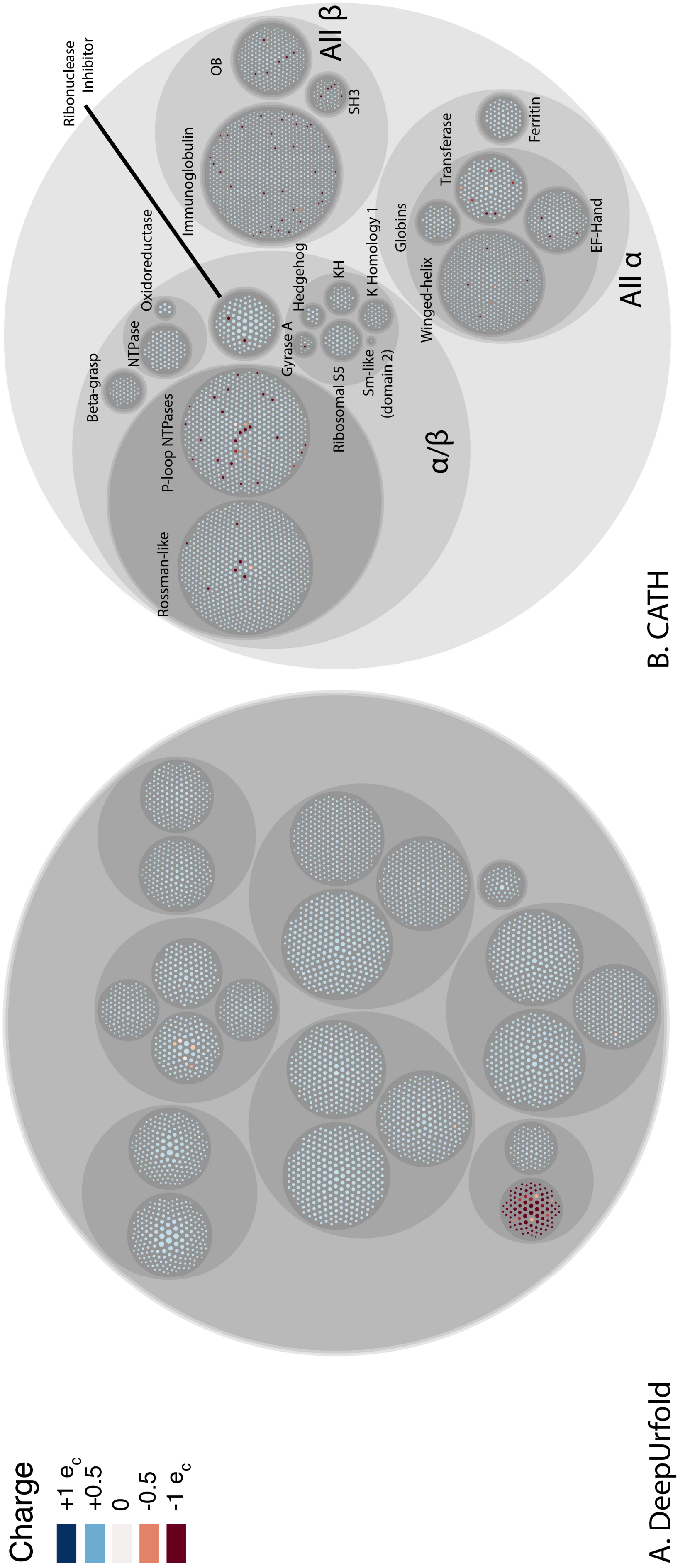

**Supplementary Figure 28:** Here, each atom is annotated with the Boolean feature is_positive. We sum up all of the positive atoms and take a fraction of the total number of atoms.

##### 1.6.4 Secondary Structure

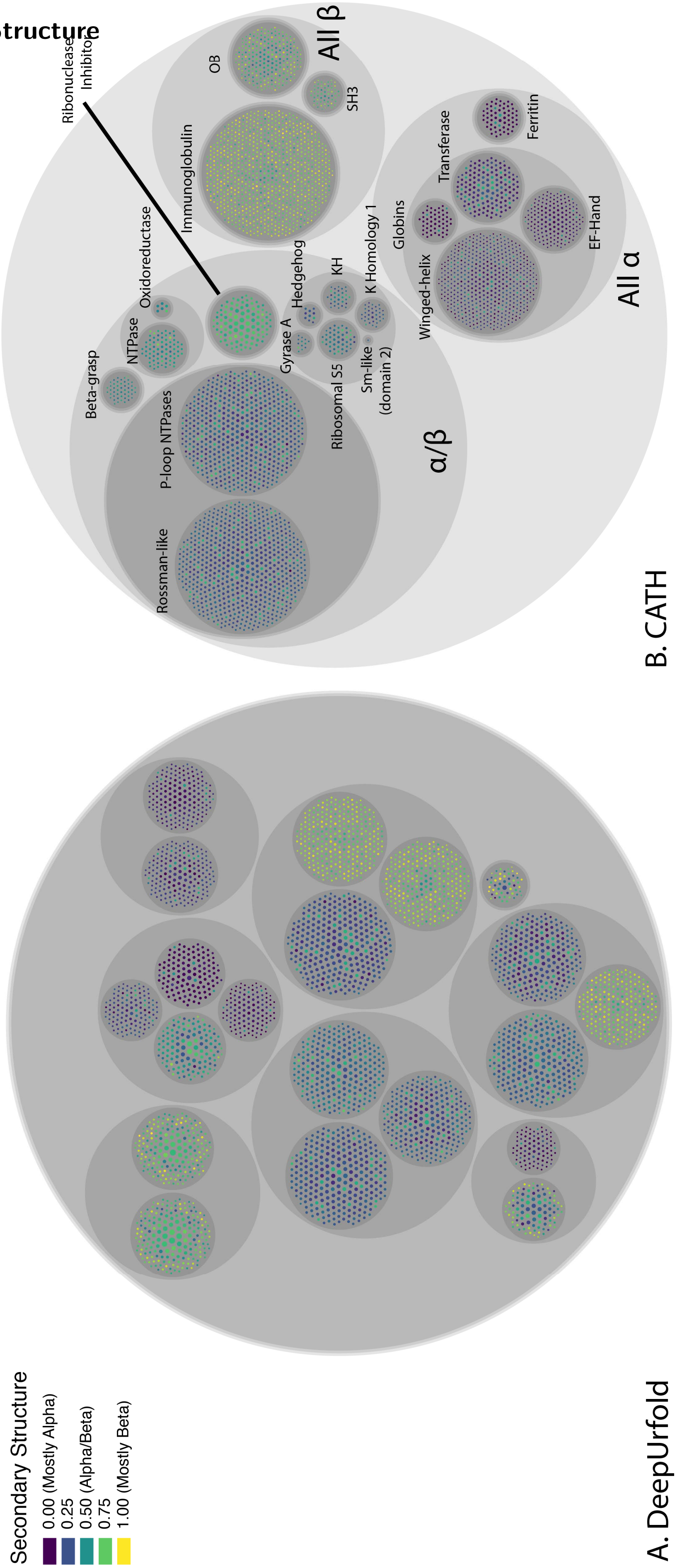

**Supplementary Figure 29:** To calculate the secondary structure score, we use the formula: (# beta atoms - # alpha atoms)/(2*(# beta atoms + # alpha atoms)) + 0.5.

##### 1.6.5 Gene Ontology

###### 1.6.5.1 Molecular Function

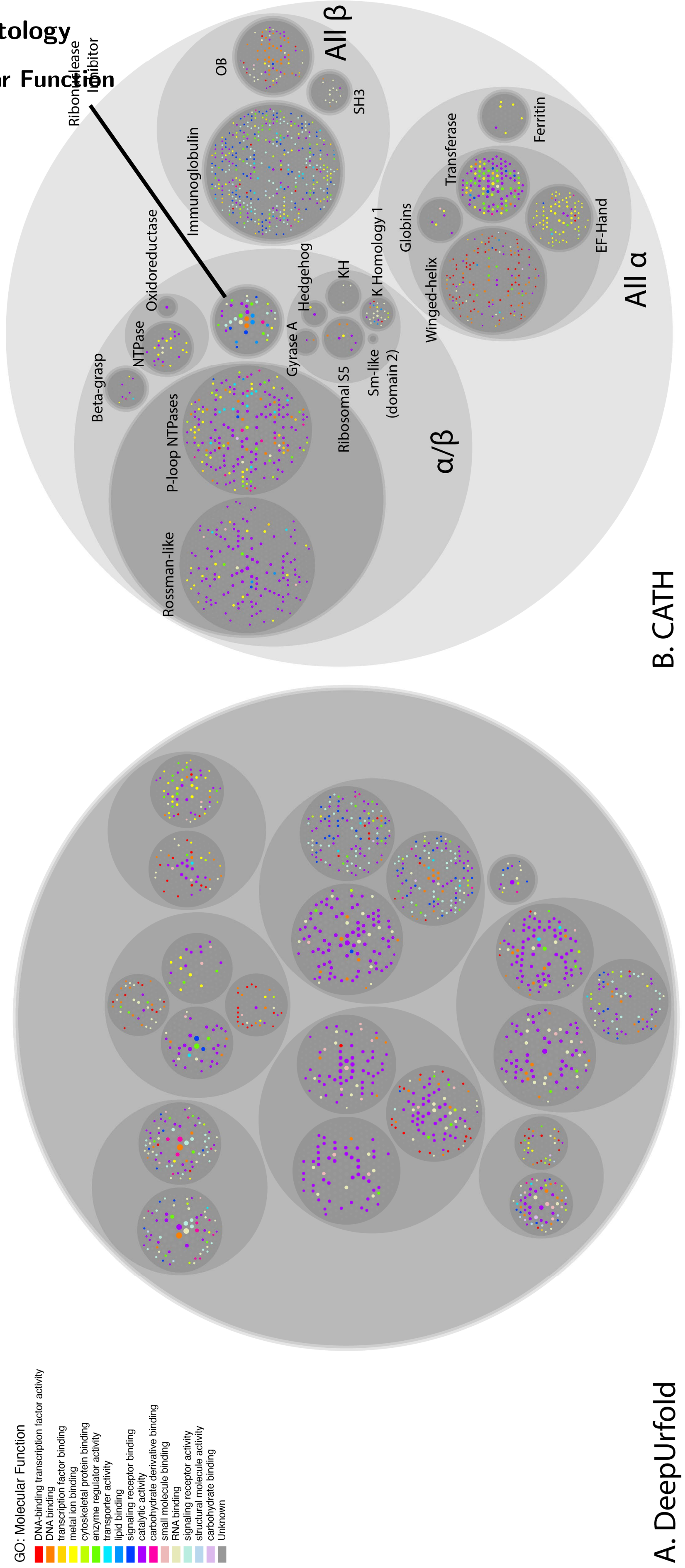

**Supplementary Figure 30:** We use GOATOOLS to calculate enrichment for each GO term from all domains in the predicted SBM community (leaf grouping only) using GO Slim terms from AGR. If the domain has a GO term that is enriched in its community ($p\_fdr\_bh \leq 0.05$), then it is colored for the associated term. If there are multiple enriched terms, only the first is used.

##### 1.6.5.2 Biological Process

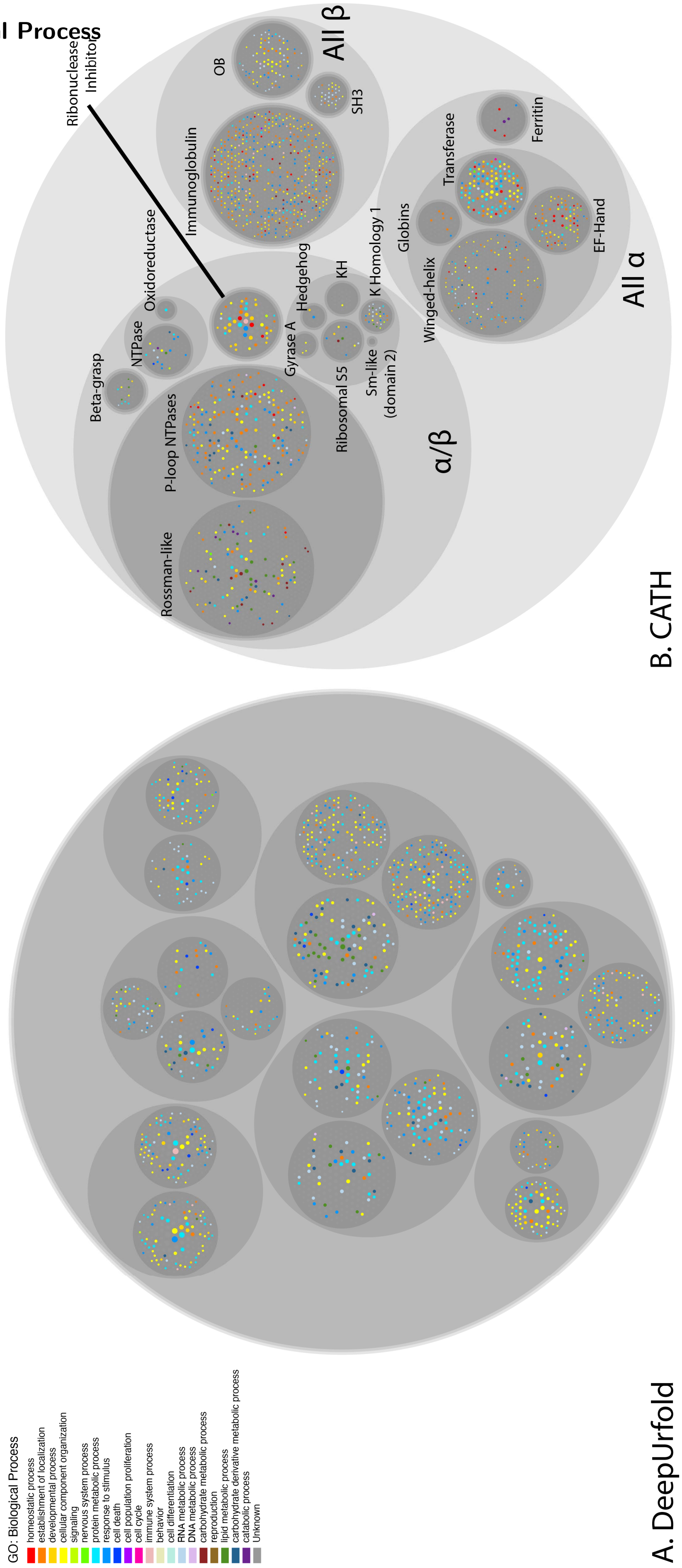

**Supplementary Figure 31:** We use GOATOOLS to calculate enrichment for each GO term from all domains in the predicted SBM community (leaf grouping only) using GO Slim terms from AGR. If the domain has a GO term that is enriched in its community (p_fdr_bh $\leq 0.05$), then it is colored for the associated term. If there are multiple enriched terms, only the first is used.

###### 1.6.5.3 Cellular Component

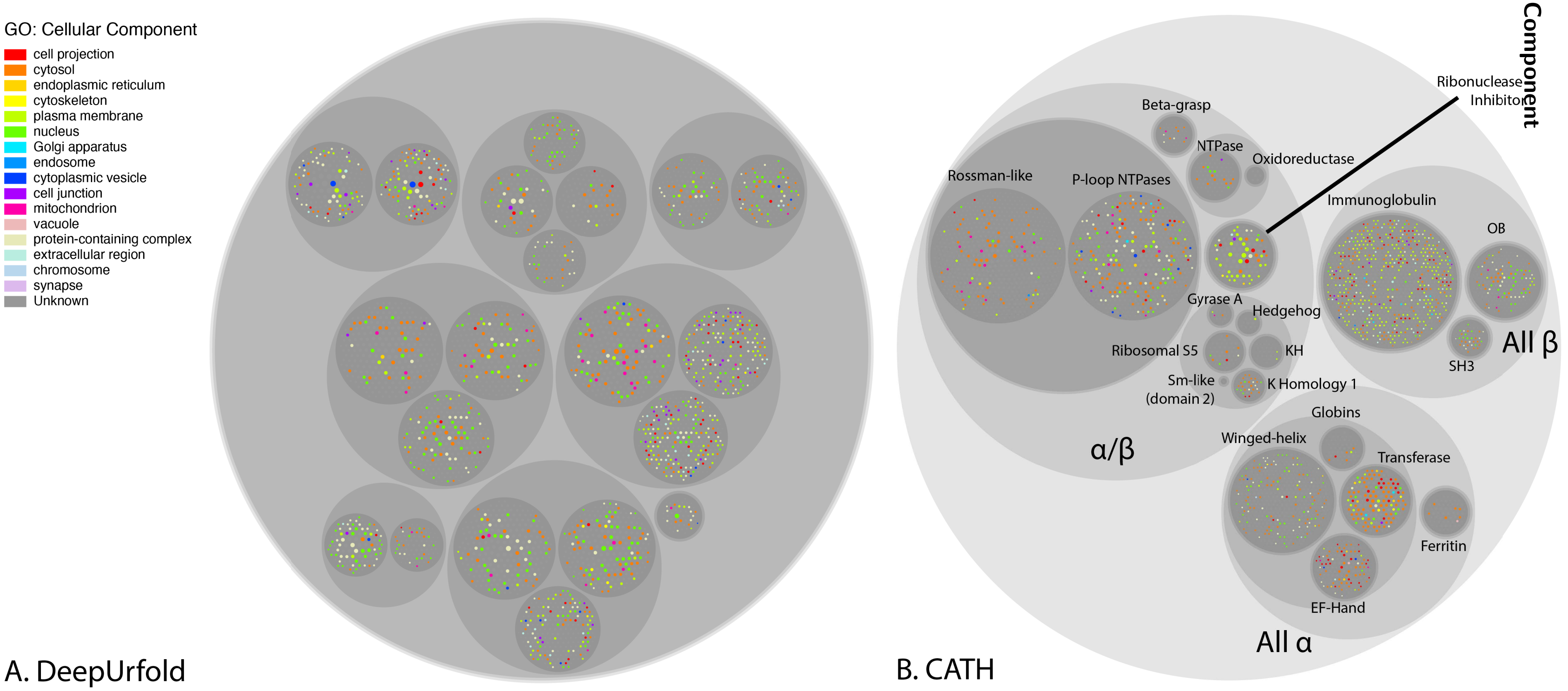

**Supplementary Figure 32:** We use GOATOOLS to calculate enrichment for each GO term from all domains in the predicted SBM community (leaf grouping only) using GO Slim terms from AGR. If the domain has a GO term that is enriched in its community (p_fdr_bh $\leq 0.05$), then it is colored for the associated term. If there are multiple enriched terms, only the first is used.

##### 1.6.6 A Downsampled SBM

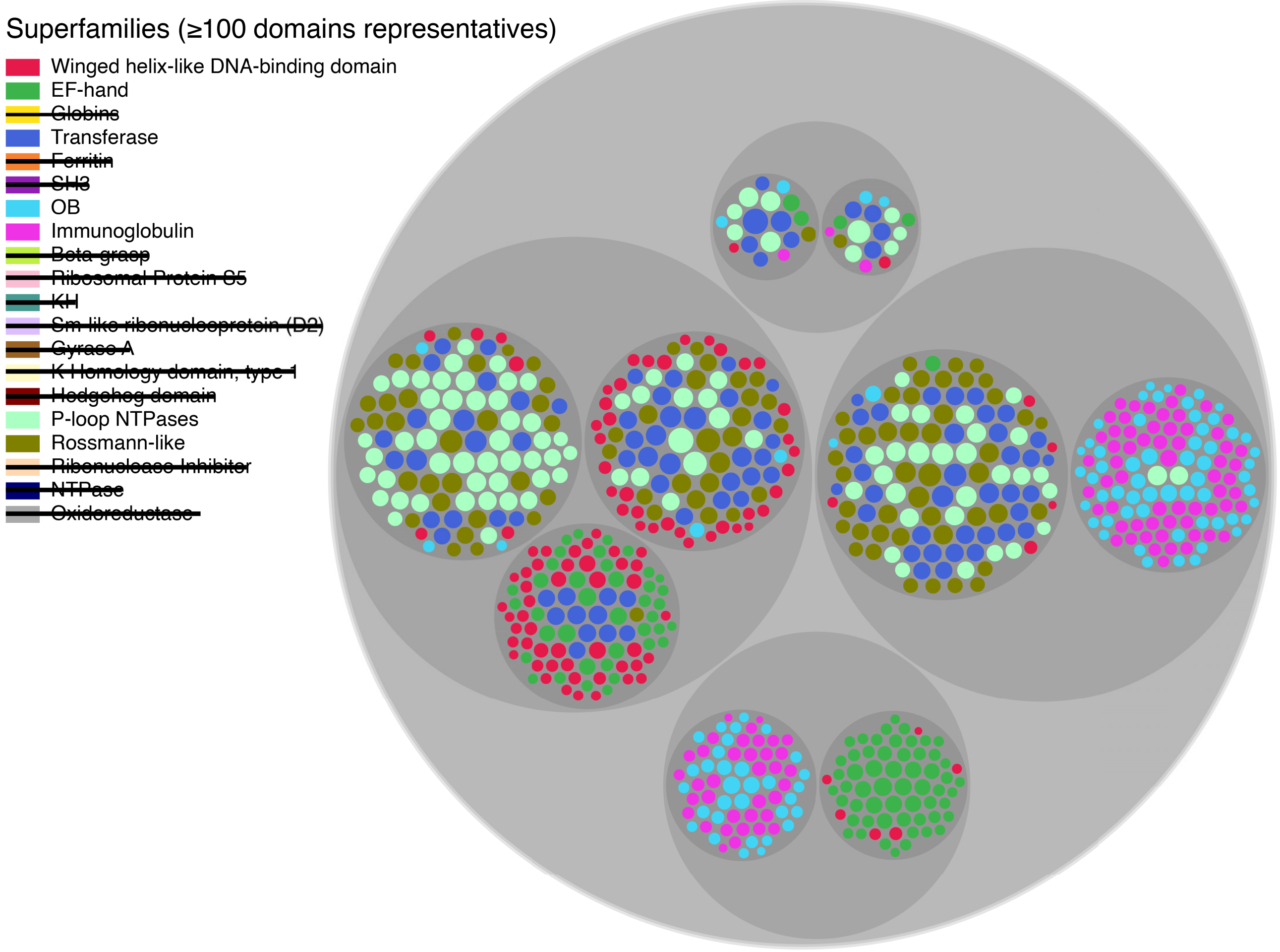

**Supplementary Figure 33:** In order to gauge how the SBM algorithm treats highly imbalanced classes in DeepUrfold, in the calculation results shown here we included only superfamilies that had $\geq$ 100 domain representatives, and we sampled 100 random domains from each. No immediate change can be detected here (versus the full SBMs shown above), and OB domains are still found in the same community as Immunoglobulins (consistent with the Urfold's model of 'intermixing' between superfamilies and higher-order structural levels [4]).

##### 1.6.7 An SBM with Random Weights

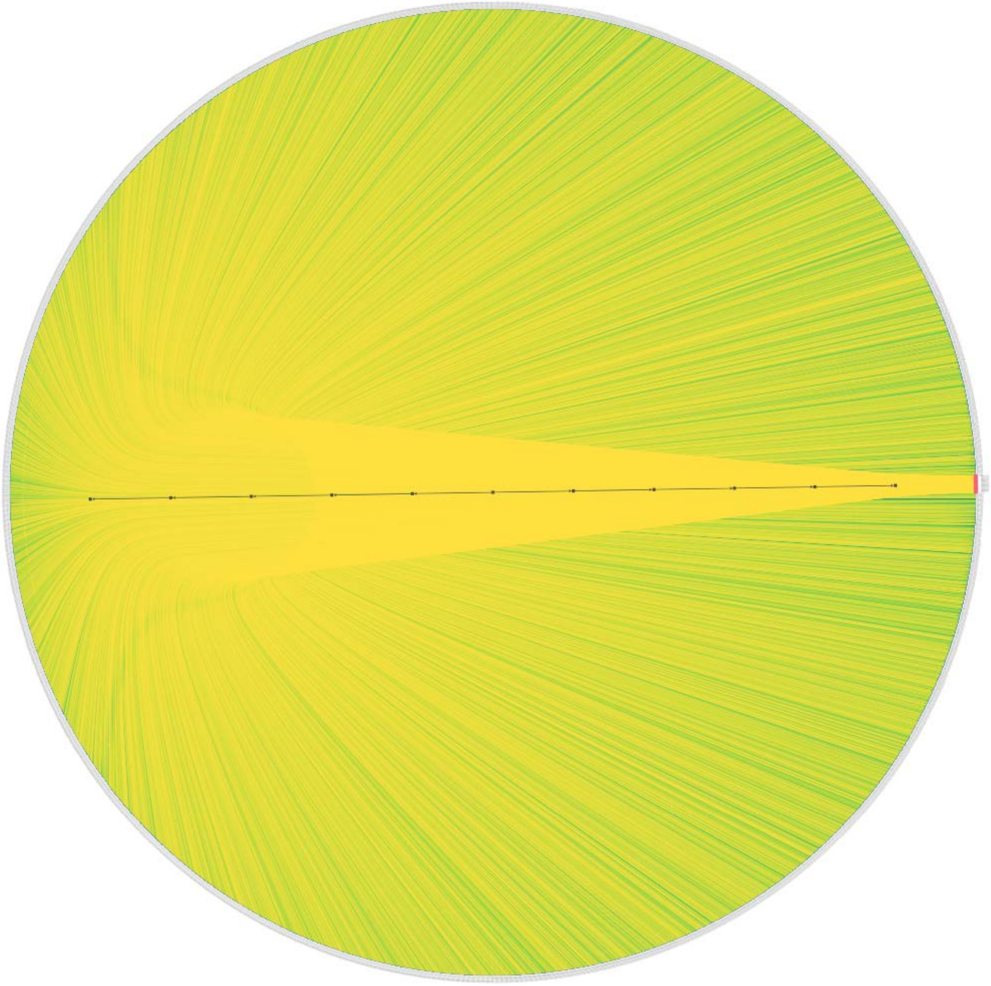

**Supplementary Figure 34:** This radial tree is for an SBM computed from a dataset of the same size as in our DeepUrfold calculations, but with randomly-weighted linkages as input. As a baseline, we sought to assess the SBM's community-detection performance given random weights (using a NumPy function like `np.random.rand(3674,20)`). Notably, this SBM calculation took over a week of wallclock time to converge, and when it did the solution contained only two groups: one for *all domains*, and another for *all superfamilies*—i.e., the exact bipartite nature of the $all_{\mathrm{SFs}} \times all_{\mathrm{domains}}$ input was recapitulated. This result indicates that the DeepUrfold model is better than random at detecting communities (see also Fig 5 in the main text): the model likelihoods from DeepUrfold might hold clues as to how we can group domains in a manner that better integrates structure and function information. A searchable, higher quality version of this tree can be found in our Zenodo archive (https://doi.org/10.5281/zenodo.13228646) under 'comparisons'.

### 2 Supplementary Tables

#### 2.1 Superfamilies Used in This Work

Supplementary Table 1: The 20 CATH superfamilies used in this study (see also Prop3D-20sf).

| CATH Code | Protein Name/Type | Description/rationale | # Domains | # Representatives | Urfold? (manual detection) |
|---|---|---|---|---|---|
| 1.10.10.10 | Winged helix–like DNA-binding domain superfamily/Winged helix DNA-binding domain | | 3444 | 524 | |
| 1.10.238.10 | EF-hand | | 1933 | 166 | |
| 1.10.490.10 | Globins | | 2891 | 52 | |
| 1.10.510.10 | Transferase (Phosphotransferase) domain 1 | | 7219 | 148 | |
| 1.20.1260.10 | Ferritin, core subunit, four-helix bundle | a major non-haem iron storage protein in animal, plants and microorganisms | 2985 | 60 | |
| 2.30.30.100 | SH3-type barrels (at T level [2.30.30]) | Includes Integrase, C-terminal domain superfamily, retroviral | 1545 | 56 | Small β-barrel (SBB) urfold |
| 2.40.50.140 | OB fold (at T level), "Nucleic acid-binding proteins" (at H level) | Dihydrolipoamide Acetyltransferase, E2P Nucleic acid-binding proteins | 2879 | 227 | SBB urfold |
| 2.60.40.10 | Immunoglobulins | | 31905 | 873 | |
| 3.10.20.30 | Beta-grasp domain | a core structure consisting of beta(2)-alpha-beta(2), which is similar to that found in ubiquitin. Domains with this type of structure are found in the 2Fe-2S ferredoxin family | 520 | 48 | Beta-grasp (Ub) |
| 3.30.1360.40 | Gyrase A; domain 2 (at T level) | | 160 | 13 | Sm-like ribonucleoproteins |
| 3.30.1370.10 | K Homology (KH) domain, type 1 | prevalent RNA-binding motif | 139 | 34 | RRM/RBD(ish) |
| 3.30.1380.10 | Hedgehog domain | | 101 | 10 | Sm-like ribonucleoproteins |
| 3.30.230.10 | Ribosomal protein S5; domain 2 | | 1274 | 45 | Sm-like ribonucleoproteins |
| 3.30.300.20 | KH domain | Could belong, together w/ the RRMs, in a new urfold? | 529 | 30 | RRM/RBD(ish) |
| 3.30.310.60 | Sm-like protein, C-term domain | C-terminal domain (TATA-binding) | 28 | 1 | Sm-like proteins |
| 3.40.50.300 | P-loop NTPases | P-loop containing nucleotide triphosphate hydrolases (NTPase; both 2AK3 and 1REB in same CATH ID) | 9233 | 561 | P-loop NTPases |
| 3.40.50.720 | NAD(P)-binding Rossmann-like domain | | 11728 | 647 | Rossmann-based urfold? |
| 3.80.10.10 | Ribonuclease Inhibitor | Leucine-rich repeat (LRR) protein; as a positive control, sanity check | 709 | 99 | |
| 3.90.420.10 | Oxidoreductase, molybdopterin-binding domain | | 58 | 6 | Beta-grasp (Ub) |
| 3.90.79.10 | Nucleoside Triphosphate Pyrophosphohydrolase | | 850 | 74 | Beta-grasp (Ub) |

#### 2.2 Metrics for SBM of DeepUrfold-based Communities

| | |
|---|---|
| **# of Clusters** | 20 |
| **Silhouette Score** | −0.0705 |
| **Davies-Boundin Score** | 33.1678 |
| **Overlap Score** | 0.2798 |
| **Rand Index** | 0.8542 |
| **Rand Index Adjusted** | 0.1977 |
| **Adjusted Mutual Information** | 0.4108 |
| **Homogeneity Score** | 0.4831 |
| **Completeness Score** | 0.3749 |

Supplementary Table 2: Provided here are the clustering metrics for DeepUrfold SBM groupings vs CATH. Note that these results also can be found in Supplementary Table 3; they are tabulated separately here simply for convenience and ease of reference.

#### 2.3 Comparison to Other Protein Sequence/Structure Similarity Tools

Supplementary Table 3: Comparison of Stochastic Block Modelling of a Bipartite graph of CATH Domains and Superfamilies with scores based on similar algorithms to DeepUrfold. The Silhouette Score and Davis-Boundin are self-measures and not comparable against CATH.

| | Method | Model | Train Data | Score | # Communities | Silhouette Score | Davis-Boundin | Overlap | Rand Index (Adjusted) | Adjusted Mutual Information | Homogeneity | Completeness |
|---|---|---|---|---|---|---|---|---|---|---|---|---|
| **Sequence-based** | | | | | | | | | | | | |
| Pairwise | UCLUST | Global Alignment | CATH S35 | ↑% Seq ID | 19 | 0.2863 | 2.4830 | 0.6148 | 0.9131 (0.6778) | 0.7703 | 0.7712 | 0.7791 |
| | | Local Alignment | CATH S35 | ↑% Seq ID | 38 | −0.2584 | 10.4408 | 0.3476 | 0.8338 (0.2730) | 0.4382 | 0.5074 | 0.4156 |
| Single Model | ESM-1b | Language Transformer | Uniref50 | ↓Euclidean Distance | 121 | 0.4299 | 0.7931 | 0.1676 | 0.8638 (0.0896) | 0.5843 | 0.9065 | 0.4311 |
| Superfamily-specific models | HMMER | HMM | CATH S35 | ↑Bitscore | 24 | −0.1715 | 5.3797 | 0.4122 | 0.7463 (0.2444) | 0.4747 | 0.4429 | 0.5416 |
| | | | EVcouplings alignment | ↑Bitscore | 14 | 0.5422 | 3.5902 | 0.3105 | 0.5546 (0.0905) | 0.2173 | 0.1734 | 0.3382 |
| | SeqDesign | Auto-regressive | EVcouplings alignment (Unaligned) | ↑Bitscore | 7 | 0.0410 | 3.6629 | 0.4095 | 0.7604 (0.2085) | 0.2736 | 0.2370 | 0.3235 |
| **Structure-based** | | | | | | | | | | | | |
| Pairwise | TM-Align | Structural Alignment | CATH S35 | ↑TM-Score | 42 | 0.1303 | 1.7138 | 0.3150 | 0.8734 (0.2380) | 0.6232 | 0.8247 | 0.5188 |
| | | | | ↓RMSD | 29 | 0.1593 | 1.8141 | 0.4200 | 0.8831 (0.3342) | 0.6628 | 0.8172 | 0.5706 |
| | | Circular Permutations | CATH S35 | ↑TM-Score | 42 | 0.1124 | 1.7407 | 0.2969 | 0.8702 (0.2162) | 0.6042 | 0.8005 | 0.5037 |
| | | | | ↓RMSD | 26 | 0.1801 | 1.6529 | 0.4164 | 0.8799 (0.3467) | 0.6740 | 0.8032 | 0.5929 |
| Superfamily-specific models | Struct2Seq | Graph Transformer | CATH (Backbone) | ↑Perplexity | 15 | 0.0563 | 2.5657 | 0.4724 | 0.8745 (0.3959) | 0.5434 | 0.5857 | 0.5196 |
| | ***DeepUrfold (ours)*** | 3D-CNN VAE | CATH (Full) | ↓-log(-ELBO) | 20 | −0.0705 | 33.1678 | 0.2798 | 0.8542 (0.1977) | 0.4108 | 0.4831 | 0.3749 |
| **Random** | | | | | | | | | | | | |
| | Uniform SBM | np.random. rand(3674,20) | | | 1 | – | – | 0.2376 | 0.1407 (0.0) | 0.0 | 0.0 | 1.0 |
| | Uniform Choice | np.random. choice(20, 3674) | | | 20 | −0.0179 | 25.7529 | 0.0743 | 0.8234 (0.0002) | 0.0003 | 0.0226 | 0.0171 |

### 3 Supplementary Notes

The following brief notes are intended to clarify some of the descriptions in the main text (wherein they are cited); they are provided here as supplementary notes so as to avoid cluttering or distracting the text:

- **Supplementary Note 1**: The term 'protein structure space' (PSS) means the set of all protein 3D structures, known and unknown; the term 'fold space' refers to the set of all protein folds. Though not strictly equivalent [4], we treat these terms interchangeably here unless noted otherwise.

- **Supplementary Note 2**: We use the capitalized term 'Urfold' to refer to the concept/theory/-model, as a general idea; the lowercase 'urfold' is used when we intend for that specific instance of the word to be limited to a specific case (e.g., "*the* SBB urfold"). Our goal is not to be dogmatic, but rather to be clear and precise as this new concept is being developed.

- **Supplementary Note 3**: As described in our guide to interpreting –(ELBO) distributions (Supplementary Figure 8 and Supplementary Discussion 5.2), this reasoning underlies the interpretation of Figure 2: Maximizing the ELBO equates to minimizing DeepUrfold's –(ELBO) loss function, which is why a shift leftwards along the horizontal axis, towards smaller (less positive) values in Figure 2, corresponds to 'better' models. Similarly, larger (more positive) values of the –(ELBO) quantity reflect poorer agreement between a domain structure and the VAE model it is being subjected to in an inference calculation (e.g., the single-tick marks in Figure 2 and Supplementary Figure 8).

- **Supplementary Note 4**: A 'consensus' model in the sense that DeepUrfold's likelihood-based scores can be viewed as measures of the goodness-of-fit of protein domain structures to VAE models that are learnt, against the variational objective, at the SF level.

- **Supplementary Note 5**: In some sense, these 'defining regions' may play analogous roles in protein domains as do *tokens* in natural language modelling and generation via large language models such as the generative pre-trained transformers (GPT-n series).

- **Supplementary Note 6**: As regards the semantics of this phrase '*sampling from*', note that a VAE's modelling/learning of this latent space distribution is what makes it a form of *generative* modelling: were one so inclined, the learnt distribution could be used to generate new instances/samples of the type of entity being modeled (a string of text, image data, etc.), in as optimal a manner as possible ('optimal' in terms of the match between statistical distributions of the generated entities relative to the observed data); more concretely, valid new entities could be created, for example, by interpolating between latent space embeddings. The generative approach contrasts with, e.g., more traditional *discriminative* models, wherein the likelihood of specific labels being associated with specific instances can be assessed and used to classify/discriminate between the different types of instances (versus spawning new ones). A benefit of generative models is that they develop a probabilistic framework that describes the statistics of the observed instance↔label mappings, thus enabling new entities to be created. Such approaches are powerful, e.g., in de novo protein design.

### 4 Supplementary Methods

#### 4.1 DeepUrfold's Model Architecture

```
DomainStructureVAE(
  (encoder): Encoder(
    (block1): Sequential(
      (0): MinkowskiConvolution(in=20, out=16,
          kernel_size=[3, 3, 3], stride=[2, 2, 2],
          dilation=[1, 1, 1])
      (1): MinkowskiSyncBatchNorm(16, eps=1e-05,
          momentum=0.1, affine=True,
          track_running_stats=True)
      (2): MinkowskiELU()
      (3): MinkowskiConvolution(in=16, out=16,
          kernel_size=[3, 3, 3], stride=[1, 1, 1],
          dilation=[1, 1, 1])
      (4): MinkowskiSyncBatchNorm(16, eps=1e-05,
          momentum=0.1, affine=True,
          track_running_stats=True)
      (5): MinkowskiELU()
    )
    (block2): Sequential(
      (0): MinkowskiConvolution(in=16, out=32,
          kernel_size=[3, 3, 3], stride=[2, 2, 2],
          dilation=[1, 1, 1])
      (1): MinkowskiSyncBatchNorm(32, eps=1e-05,
          momentum=0.1, affine=True,
          track_running_stats=True)
      (2): MinkowskiELU()
      (3): MinkowskiConvolution(in=32, out=32,
          kernel_size=[3, 3, 3], stride=[1, 1, 1],
          dilation=[1, 1, 1])
      (4): MinkowskiSyncBatchNorm(32, eps=1e-05,
          momentum=0.1, affine=True,
          track_running_stats=True)
      (5): MinkowskiELU()
    )
    (block3): Sequential(
      (0): MinkowskiConvolution(in=32, out=64,
          kernel_size=[3, 3, 3], stride=[2, 2, 2],
          dilation=[1, 1, 1])
      (1): MinkowskiSyncBatchNorm(64, eps=1e-05,
          momentum=0.1, affine=True,
          track_running_stats=True)
      (2): MinkowskiELU()
      (3): MinkowskiConvolution(in=64, out=64,
          kernel_size=[3, 3, 3], stride=[1, 1, 1],
          dilation=[1, 1, 1])
      (4): MinkowskiSyncBatchNorm(64, eps=1e-05,
          momentum=0.1, affine=True,
          track_running_stats=True)
      (5): MinkowskiELU()
    )
    (block4): Sequential(
      (0): MinkowskiConvolution(in=64, out=128,
          kernel_size=[3, 3, 3], stride=[2, 2, 2],
          dilation=[1, 1, 1])
      (1): MinkowskiSyncBatchNorm(128, eps=1e-05,
          momentum=0.1, affine=True,
          track_running_stats=True)
      (2): MinkowskiELU()
      (3): MinkowskiConvolution(in=128, out=128,
          kernel_size=[3, 3, 3], stride=[1, 1, 1],
          dilation=[1, 1, 1])
      (4): MinkowskiSyncBatchNorm(128, eps=1e-05,
          momentum=0.1, affine=True,
          track_running_stats=True)
      (5): MinkowskiELU()
    )
    (block5): Sequential(
      (0): MinkowskiConvolution(in=128, out=256,
          kernel_size=[3, 3, 3], stride=[2, 2, 2],
          dilation=[1, 1, 1])
      (1): MinkowskiSyncBatchNorm(256, eps=1e-05,
          momentum=0.1, affine=True,
          track_running_stats=True)
      (2): MinkowskiELU()
      (3): MinkowskiConvolution(in=256, out=256,
          kernel_size=[3, 3, 3], stride=[1, 1, 1],
          dilation=[1, 1, 1])
      (4): MinkowskiSyncBatchNorm(256, eps=1e-05,
          momentum=0.1, affine=True,
          track_running_stats=True)
      (5): MinkowskiELU()
    )
    (block6): Sequential(
      (0): MinkowskiConvolution(in=256, out=512,
          kernel_size=[3, 3, 3], stride=[2, 2, 2],
          dilation=[1, 1, 1])
      (1): MinkowskiSyncBatchNorm(512, eps=1e-05,
          momentum=0.1, affine=True,
          track_running_stats=True)
      (2): MinkowskiELU()
      (3): MinkowskiConvolution(in=512, out=512,
          kernel_size=[3, 3, 3], stride=[1, 1, 1],
          dilation=[1, 1, 1])
      (4): MinkowskiSyncBatchNorm(512, eps=1e-05,
          momentum=0.1, affine=True,
          track_running_stats=True)
      (5): MinkowskiELU()
    )
    (block7): Sequential(
      (0): MinkowskiConvolution(in=512, out=1024,
          kernel_size=[3, 3, 3], stride=[2, 2, 2],
          dilation=[1, 1, 1])
```

```
    (1): MinkowskiSyncBatchNorm(1024, eps=1e-05,
        momentum=0.1, affine=True,
        track_running_stats=True)
    (2): MinkowskiELU()
    (3): MinkowskiConvolution(in=1024, out=1024,
        kernel_size=[3, 3, 3], stride=[1, 1, 1],
        dilation=[1, 1, 1])
    (4): MinkowskiSyncBatchNorm(1024, eps=1e-05,
        momentum=0.1, affine=True,
        track_running_stats=True)
    (5): MinkowskiELU()
  )
  (global_pool): MinkowskiGlobalPooling(mode=
      PoolingMode.
      GLOBAL_AVG_POOLING_PYTORCH_INDEX)
  (linear_mean): MinkowskiLinear(in_features
      =1024, out_features=1024, bias=True)
  (linear_log_var): MinkowskiLinear(in_features
      =1024, out_features=1024, bias=True)
)
(decoder): Decoder(
  (block1): Sequential(
    (0): MinkowskiConvolutionTranspose(in=1024,
        out=1024, kernel_size=[2, 2, 2], stride
        =[2, 2, 2], dilation=[1, 1, 1])
    (1): MinkowskiSyncBatchNorm(1024, eps=1e-05,
        momentum=0.1, affine=True,
        track_running_stats=True)
    (2): MinkowskiELU()
    (3): MinkowskiConvolution(in=1024, out=1024,
        kernel_size=[3, 3, 3], stride=[1, 1, 1],
        dilation=[1, 1, 1])
    (4): MinkowskiSyncBatchNorm(1024, eps=1e-05,
        momentum=0.1, affine=True,
        track_running_stats=True)
    (5): MinkowskiELU()
    (6): MinkowskiConvolutionTranspose(in=1024,
        out=512, kernel_size=[2, 2, 2], stride
        =[2, 2, 2], dilation=[1, 1, 1])
    (7): MinkowskiSyncBatchNorm(512, eps=1e-05,
        momentum=0.1, affine=True,
        track_running_stats=True)
    (8): MinkowskiELU()
    (9): MinkowskiConvolution(in=512, out=512,
        kernel_size=[3, 3, 3], stride=[1, 1, 1],
        dilation=[1, 1, 1])
    (10): MinkowskiSyncBatchNorm(512, eps=1e-05,
        momentum=0.1, affine=True,
        track_running_stats=True)
    (11): MinkowskiELU()
  )
  (block2): Sequential(
    (0): MinkowskiConvolutionTranspose(in=512,
        out=256, kernel_size=[2, 2, 2], stride
        =[2, 2, 2], dilation=[1, 1, 1])
    (1): MinkowskiSyncBatchNorm(256, eps=1e-05,
        momentum=0.1, affine=True,
        track_running_stats=True)
    (2): MinkowskiELU()
    (3): MinkowskiConvolution(in=256, out=256,
        kernel_size=[3, 3, 3], stride=[1, 1, 1],
        dilation=[1, 1, 1])
    (4): MinkowskiSyncBatchNorm(256, eps=1e-05,
        momentum=0.1, affine=True,
        track_running_stats=True)
    (5): MinkowskiELU()
  )
  (block3): Sequential(
    (0): MinkowskiConvolutionTranspose(in=256,
        out=128, kernel_size=[2, 2, 2], stride
        =[2, 2, 2], dilation=[1, 1, 1])
    (1): MinkowskiSyncBatchNorm(128, eps=1e-05,
        momentum=0.1, affine=True,
        track_running_stats=True)
    (2): MinkowskiELU()
    (3): MinkowskiConvolution(in=128, out=128,
        kernel_size=[3, 3, 3], stride=[1, 1, 1],
        dilation=[1, 1, 1])
    (4): MinkowskiSyncBatchNorm(128, eps=1e-05,
        momentum=0.1, affine=True,
        track_running_stats=True)
    (5): MinkowskiELU()
  )
  (block4): Sequential(
    (0): MinkowskiConvolutionTranspose(in=128,
        out=64, kernel_size=[2, 2, 2], stride=[2,
         2, 2], dilation=[1, 1, 1])
    (1): MinkowskiSyncBatchNorm(64, eps=1e-05,
        momentum=0.1, affine=True,
        track_running_stats=True)
    (2): MinkowskiELU()
    (3): MinkowskiConvolution(in=64, out=64,
        kernel_size=[3, 3, 3], stride=[1, 1, 1],
        dilation=[1, 1, 1])
    (4): MinkowskiSyncBatchNorm(64, eps=1e-05,
        momentum=0.1, affine=True,
        track_running_stats=True)
    (5): MinkowskiELU()
  )
  (block5): Sequential(
    (0): MinkowskiConvolutionTranspose(in=64, out
        =32, kernel_size=[2, 2, 2], stride=[2, 2,
         2], dilation=[1, 1, 1])
    (1): MinkowskiSyncBatchNorm(32, eps=1e-05,
        momentum=0.1, affine=True,
        track_running_stats=True)
    (2): MinkowskiELU()
    (3): MinkowskiConvolution(in=32, out=32,
        kernel_size=[3, 3, 3], stride=[1, 1, 1],
        dilation=[1, 1, 1])
    (4): MinkowskiSyncBatchNorm(32, eps=1e-05,
        momentum=0.1, affine=True,
        track_running_stats=True)
    (5): MinkowskiELU()
```

```
)
(block6): Sequential(
  (0): MinkowskiConvolutionTranspose(in=32, out
      =16, kernel_size=[2, 2, 2], stride=[2, 2,
       2], dilation=[1, 1, 1])
  (1): MinkowskiSyncBatchNorm(16, eps=1e-05,
      momentum=0.1, affine=True,
      track_running_stats=True)
  (2): MinkowskiELU()
  (3): MinkowskiConvolution(in=16, out=16,
      kernel_size=[3, 3, 3], stride=[1, 1, 1],
      dilation=[1, 1, 1])
  (4): MinkowskiSyncBatchNorm(16, eps=1e-05,
      momentum=0.1, affine=True,
      track_running_stats=True)
  (5): MinkowskiELU()
)
(block7): MinkowskiConvolution(in=16, out=20,
    kernel_size=[1, 1, 1], stride=[1, 1, 1],
    dilation=[1, 1, 1])
(pruning): MinkowskiPruning()
)
)
```

### 5 Supplementary Discussion

#### 5.1 Superfamilies Used in This Work: A Brief Synopsis

We chose the particular 20 superfamilies that were the subject of this work (Supp Table 1) for a few reasons, delineated as follows:

- They are highly-populated and diverse superfamilies. Specifically, they are among the SFs with the most abundant structural representatives in CATH (e.g., immunoglobulins, globins, transferases, EF-hand domains, ferritin, etc.). Also, this collection of SFs is diverse, in terms of CATH (i) Classes (mostly-α, mostly-β and mixed α/β are included) and (ii) Architectures (e.g., 'orthogonal bundles' [1.10], 'up-down bundles' [1.20], β 'rolls' [2.30], 'barrels' [2.40] and 'sandwiches' [2.60] are represented, as are 2-layer α/β sandwiches [3.30], etc.).

- There is substantial evidence from past studies (albeit more qualitative, manual analyses by human experts) that some of these SFs house distinct urfolds—e.g., the SH3/Sm and OB, which together define the 'small β-barrel' (SBB) domain that was first identified and described as an '*urfold*' by Youkharibache et al. [5]; the Rossmann and P-loop NTPases [4]; the β-grasp; gyrase domains; Hedgehog domains; KH domains, and so on.

- The particular SF is included because it might help stress-test our models and overall DeepUrfold framework, or it serves as a 'positive control' of sorts, e.g., via inclusion of the leucine-rich repeats (LRRs; CATH ID 3.80.10.10) in our 'Prop3D-20sf' dataset [1].

Finally, we note that the DeepUrfold work reported here utilizes a Prop3D-20sf dataset that we recently created as a community resource—indeed, that dataset provides "a hand-crafted, already-featurized dataset of protein domains for 20 highly-populated CATH families", corresponding exactly to the 'top-20' SFs used in this work. Further details about the construction and processing steps involved in creating Prop3D-20sf, including its data splits for model training/testing/validation, factors such as (mitigating) evolutionary data leakage, and so on, can be found in our recent Prop3D article [1].

#### 5.2 Interpreting DeepUrfold's ELBO-based Scores & Histograms

In training DeepUrfold's VAE models, we formulate the optimization problem via an ELBO-based objective function that we seek to minimize: Specifically, we minimize a –(ELBO) loss function, which equates to maximizing the absolute value, |ELBO|. This, then, means that lower values of the –(ELBO) quantity (plotted as the abscissa in Figure 2, and in Supplementary Figures 8 and 10) are 'better', insofar as minimizing –(ELBO) corresponds to (i) maximizing the expected log-likelihood of our learned model and (ii) minimizing the relative entropy or 'distance' (Kullback–Leibler [KL] divergence, $D_{\mathrm{KL}}$, which is strictly nonnegative) between (a) the true underlying distribution of the data given a model, $p(x|\theta)$, and (b) our learned/inferred posterior distribution of latent parameters given the data, $q(z|x)$. More explicitly, DeepUrfold's VAE training employs a loss function that can be viewed in the following form (the left-hand side, –(ELBO) values, is what we plot as the abscissa in Fig 2 of the main text and elsewhere in this work):

$$-\mathcal{L}(\theta,\phi;x^{(i)}) = \underbrace{\omega \cdot D_{\mathrm{KL}}(q_\phi(z|x^{(i)})\,||\,p_\theta(z))}_{◐} + \underbrace{\mathbb{E}_{q_\phi(z|x^{(i)})}[-\log\ p_\theta(x^{(i)}|z)]}_{◑},$$

where $\mathcal{L}$ is the loss function; $\omega$ is a weight for the $D_{\mathrm{KL}}$ term; $\theta$ and $\phi$ are the statistical model parameters; $x^{(i)}$ denotes the $i^{\text{th}}$ data point; $p(z)$ is the prior distribution of $z$; $q_\phi(z|x^{(i)})$ is the distribution of $z$ given $x^{(i)}$ under the parameters $\phi$; and $p_\theta(x^{(i)}|z)$ is the probability of $x^{(i)}$ given $z$ under the parameters $\theta$.

This formulation of our variational objective shows the –(ELBO) as being constructed from the expected negative log-likelihoods (the ◑ term) along with a 'penalty term' (the $D_{\mathrm{KL}}$; ◐) that measures how far the approximate posterior ($q$) is from the exact/true prior ($p$). These ELBO and ELBO-based scores provide useful bounds in approximating likelihoods (floors or ceilings, depending on negative signs and whether minima or maxima are being optimized for). In the context of DeepUrfold, these likelihoods, in turn, are measures of the goodness-of-fit of protein domain structures to VAE models that are learnt, against the variational objective, at the superfamily level (other than for our monolithic 'Global Model', which is learned across all SFs).

The above reasoning underlies the interpretation of several plots in this work (e.g., Figure 2 and Supplementary Figure 8): Maximizing the ELBO equates to minimizing DeepUrfold's –(ELBO) loss function, which is why a shift leftwards along the horizontal axis in Figure 2 corresponds to 'better' fits between data and VAE models. Similarly, more positive values of the –(ELBO) quantity reflect poorer agreement between a domain structure and the VAE model it is being subjected to in an inference calculation (e.g., the single-tick marks in Figure 2 and Supplementary Figure 10); see also the analysis provided in Supplementary Figure 10 and the schematic rendition in Supplementary Figure 8.

#### 5.3 Latent Space Visualization and Embeddings

##### 5.3.1 Individual, Superfamily-level Feature Embeddings via UMAP

In this strategy for exploratory visualization of DeepUrfold-derived latent space embeddings, which can be viewed as a '*SF-level UMAPs*' approach, we consider a simple case: for an individual, SF-specific VAE model, such as we have presented in this work for 20 highly-populated SFs (Supplementary Table 1), one can 'cleanly' perform a UMAP-type dimensionality reduction of the latent space embeddings for that single SF's learned VAE model, without any complications stemming from concatenation or any other sort of 'aggregation' operation. While this tactic does not afford a holistic view of all of fold space, it is clean at the SF level. The lengthy, multi-panel Supplementary Figure 18 presents the results of such calculations for all 20 SFs. Reassuringly, the finer-grained, 'SF-resolved' UMAP results shown in that figure are consistent with the 'higher'-level results—i.e., from applying UMAP to either an aggregate of all 20 SF embeddings (Figure 3B's 'concatenated' embeddings approach), or to Figure 3C's 'Global Model'—insofar as a separation between α-rich and β-rich regions is evident in the individual SF-level panels, with a gradient between the regions (again, cf. Figure 3 of the main text).

In addition, some initial trends can be empirically detected across the various SFs shown in Supplementary Figure 18. Specifically, at least three phenomenological 'categories' or 'types' of UMAP patterns appear to emerge in Supp Fig 18's collection of results:

- A noticeably 'streaky' type of distribution, wherein the UMAP results feature a rather tight and anisotropic clustering of points, e.g., linear-shaped or crescent-shaped (panels c, d, h, p and q) Anecdotally, many of these CATH SFs appear to have exceedingly high sequence diversity (e.g., the Igs of `2.60.40.10` (panel h), the P-loop NTPases in `3.40.50.300` (panel p), and the Rossmann domains in `3.40.50.720` (panel q)).

- A more homogeneously 'smeared-out' pattern—that, nevertheless, still exhibits dispersion between the α and β classes—can be seen, e.g. in SFs `1.10.10.10` (panel a) and `3.30.1370.10` (panel n).

- A more notably 'bilobal' pattern, with a more intense separation between the α-rich and β-rich regions, can be seen to be exhibited by some SFs which are more extreme in their SSE content—e.g., (almost-)all-α, as with ferritin subunits (`1.20.1260.10`; panel e) and other 4-helix-bundle proteins, or else highly β-rich (e.g., the LRRs of the ribonuclease inhibitors, SF `3.80.10.10`; panel r).

While the above is a qualitative and anecdotal description thus far, we suspect that further exploration of these trends may be informative as regards elucidating the possible structural correlates/determinants of the learned feature embeddings.

##### 5.3.2 Patterns of Proximity among Latent Space Embeddings

Does the proximity of two given feature embeddings in DeepUrfold's latent spaces, corresponding to two given protein domains, 'track' with traditional measures of protein similarity—e.g., via 3D structure alignment, quantified as a pairwise TM-score? To explore this question, we initially computed and compared (i) a quantity based on UMAP embedding cosine distances and (ii) TM-scores for 3D structure alignment of the corresponding protein pairs. A linear regression plot of (i) versus (ii) was quite disperse (not shown here), with weak but nevertheless positive Pearson correlation coefficients. The imperfection of this trend is interesting in at least two ways: (i) First, the weakness of the correlation is consistent with a key theme of the DeepUrfold approach—namely that 3D geometric structure alone is insufficient in capturing protein interrelationships, as other characteristics (biophysical properties, phylogenetic information, etc.) help define the *sequence* ⇜⇝ *structure* ⇜⇝ *function* triad upon which evolution acts. (ii) Second, even if considering purely 3D structure were sufficient, the mappings (feature embeddings) learnt by DeepUrfold's SF-specific VAE models are free to operate at the level of sub-domain fragments, whereas the TM-score is a strictly domain-level metric. In other words, even if two domains were quite similar—in terms of belonging to the same urfold, featuring structurally-similar sub-domain fragments (urfolds) that covered, say, 2/3 of a domain (but were in different relative orientations), etc.—then these would give similar (nearby) feature embeddings in DeepUrfold, but would give only poor TM-scores. Finally, we note that our UMAP projections of the VAE latent space representations are purely for qualitative visualization purposes, and the embeddings themselves have not been optimized with, e.g., a vector quantized VAE (VQ-VAE) type of approach; therefore, one cannot draw too strong of inferences from these data.

Notwithstanding all of the above, we further assessed possible correlations between TM-scores and DeepUrfold embeddings by examining patterns of proximity between embeddings—for example, as raw embedding (cosine) distances, Euclidean distances, cosine distance quantities based upon UMAP reductions of the embeddings (versus raw embeddings), and so on. These calculations, described in Supplementary §1.5.6 and its accompanying Supplementary Figures 23-25, were done to try and directly compare (i) pairwise UMAP embedding distances versus (ii) the corresponding TM-scores between 3D structures, for all 3,764 domains in the Prop3D-20sf dataset. In the plots of such information, shown in Supplementary Figures 23-25, the data are randomly sampled across only 1,000 domains. While the correlation is weak, there is nevertheless a visible and significant inverse correlation between increasing TM-scores and decreased embedding distances (raw and from UMAP). In other words, greater 3D structural similarity corresponds to 'closer' embeddings. In those figures, note the concentration of data-points in the 'grey zone' of TM-scores $\approx$ 0.25-0.4, where two protein domain structures would not be considered as similar even though they may share some non-random elements of structural similarity.
